# Hippocampal place cells encode global location but not changes in environmental connectivity in a 4-room navigation task

**DOI:** 10.1101/2020.10.20.346130

**Authors:** Éléonore Duvelle, Roddy M Grieves, Anyi Liu, Selim Jedidi-Ayoub, Joanna Holeniewska, Adam Harris, Nils Nyberg, Francesco Donnarumma, Julie M. Lefort, Kate J. Jeffery, Christopher Summerfield, Giovanni Pezzulo, Hugo J. Spiers

**Author notes:** These authors contributed equally to this work (Éléonore Duvelle, Roddy M Grieves). Corresponding author(s): Éléonore Duvelle, Hugo Spiers.

## Abstract

Flexible navigation relies on a cognitive map of space, thought to be implemented by hippocampal place cells: neurons that exhibit location-specific firing. In connected environments, optimal navigation requires keeping track of one’s location and of the available connections between subspaces. We examined whether the dorsal CA1 place cells of rats encode environmental connectivity in four geometrically-identical boxes arranged in a square. Rats moved between boxes by pushing saloon-type doors that could be locked in one or both directions. While rats demonstrated knowledge of environmental connectivity, their place cells did not respond to connectivity changes, nor did they represent doorways differently from other locations. Importantly, place cells coded the space in a global frame, expressing minimal repetitive fields despite the repetitive geometry (global coding). These results suggest that CA1 place cells provide a spatial map that does not explicitly include connectivity.

## Introduction

Real-world navigation involves traversing complex environments, such as offices, subway systems or burrows that often have multiple connected subspaces. How are these represented in the brain? Environmental features can be categorised as contextual (e.g. colour, odour), topographic (e.g. angles, distances) or topologic (e.g. containment, connectivity). Hippocampal place cells, with their remarkable location-specific firing (O’Keefe and Dostrovsky 1971), are thought to form the neural basis of a spatial cognitive map underlying spatial memory and flexible navigation (O’Keefe and Nadel 1978). Place cells encode topographical and contextual information, as their firing reflects changes in parameters such as environment geometry (S. Leutgeb et al. 2005), colour (Jeffery and Anderson 2003), size (Fenton et al. 2008; Muller and Kubie 1987) and the spatial arrangement of connected compartments (Grieves et al. 2016). Recent studies suggest that topology is also encoded, *implicitly*, in the co-firing of place cells (Dabaghian, Brandt, and Frank 2014). For instance, in a track with changing topography but unchanging topology, the order of active place cells was stable (Dabaghian, Brandt, and Frank 2014); Chen and colleagues (2013), using a probabilistic modelling approach, recovered the topology of different environments from place cell firing; Wu and Foster (2014) found that reactivated place cell sequences (‘replay’) reflected both the length (topography) and structure (topology) of a Y-shaped maze. Here, we sought to test if place cells encode connectivity changes *explicitly* in their firing rate and firing location.

The insertion of barriers into an environment induces changes in firing rates and firing locations proximal to the barrier (Muller and Kubie 1987; Rivard et al. 2004; Alvernhe, Save, and Poucet 2011). Such local remapping is consistent with the predictions of the boundary vector cell (BVC) model, in which the spatial firing of place cells originates from the combined activity of neurons sensitive to local boundaries (Barry et al. 2006; Hartley et al. 2000). However, it has recently been proposed that such local remapping may be linked to changes in the connectivity, rather than purely the local sensory inputs. In this view, place cells form a predictive map or ‘successor representation’ of the environment (Stachenfeld, Botvinick, and Gershman 2017; see also Gustafson and Daw 2011). According to this model, altering connectivity should change the firing of place fields local to the change as this impacts possible future states. Related to this, past studies have found increased activity in doorways in multicompartmented environments (Spiers et al. 2013; Grieves et al. 2016), which could be explained by doorways being key bottlenecks connecting subspaces, i.e., have a special connectivity status; however, such overrepresentation can also be explained by the BVC model (Grieves, Duvelle, and Dudchenko 2018). Additionally, the representation of connected compartments can either be almost identical (place field repetition) or different depending on how compartments are oriented with respect to each other, even though the connectivity is not affected (Grieves et al. 2016). This is also explained by the BVC model of place cell firing (Grieves, Duvelle, and Dudchenko 2018). To our knowledge, no experiment has directly investigated hippocampal neural activity during changes in connectivity whilst maintaining consistent environmental geometry and controlling for parameters known to influence place cell activity, such as movement direction (Muller et al. 1994; Navratilova et al. 2012), running speed (McNaughton, Barnes, and O’Keefe 1983) and texture changes (Wang, Monaco, and Knierim 2020).

To address the question of how compartmented spaces and their connectivity are represented by place cells, we used an environment composed of four square boxes (60×60cm) connected in a 2 x 2 array by pushable doors and surrounded by distal cues. All boxes shared an identical geometry to facilitate cross-box comparisons; the connectivity between them could be seamlessly modified by locking doors without changing their appearance or the geometry of the environment. We trained rats to perform a guided foraging task in this environment, where rats alternated between goal-directed navigation to a briefly sound-cued box and pseudo-random foraging in that box. We compared behaviour and neural activity between baseline conditions where all doors were open and test conditions where some of the doors were locked. Specifically, we asked the following questions: i) are rats able to learn the locked or open status of individual doors and to optimally navigate in the environment? ii) does the firing rate or firing location of place cells change following connectivity changes? iii) to what extent do place cells encode position in a *local* reference frame, with repeating place fields in the same local place in all compartments, versus a *global* reference frame, with a different set of cells active in different positions in each compartment? We found that rats were sensitive to changes in connectivity and differentiated locked and open doors, but place cells did not, in any of the measures we used. This was not due to a failure of the cognitive map to discriminate between boxes: despite the repeating geometry, the population of place cells represented location globally, i.e. with a different map for each box.

## Results

### Performance in the 4-room navigation task

We first characterised the behaviour of rats in terms of door choices to determine if they had acquired knowledge of the changing connectivity. Behavioural data were obtained from 5 trained rats performing guided foraging in the ‘4-room’ environment (see Fig. 1, Sup. Fig.1, Sup. Video 1, *Methods - Training and Methods - Task*). Each rat provided 2 or more sequences of 5 sessions in two conditions: ‘Closed-Door’ (Fig. 1B) or ‘One-Way’ (Fig. 1C) for a total of 19 closed sequences and 11 One-Way sequences, excluding sequences where rats stopped performing the task before completing the full sequence (8 Closed-Door and 1 One-Way excluded). Each sequence started with two control sessions where all doors were unlocked, proceeding to two test sessions where either one of the doors was locked both ways (‘Closed-Door’ sequence) or all doors were locked one way (‘One-Way’ sequence), ending with a control session. Sessions lasted 23 minutes on average and ended after 12 trials; they were followed by a rest period on an elevated platform for around 10 min, with a total daily recording duration of about 2h30.

**Figure 1:**
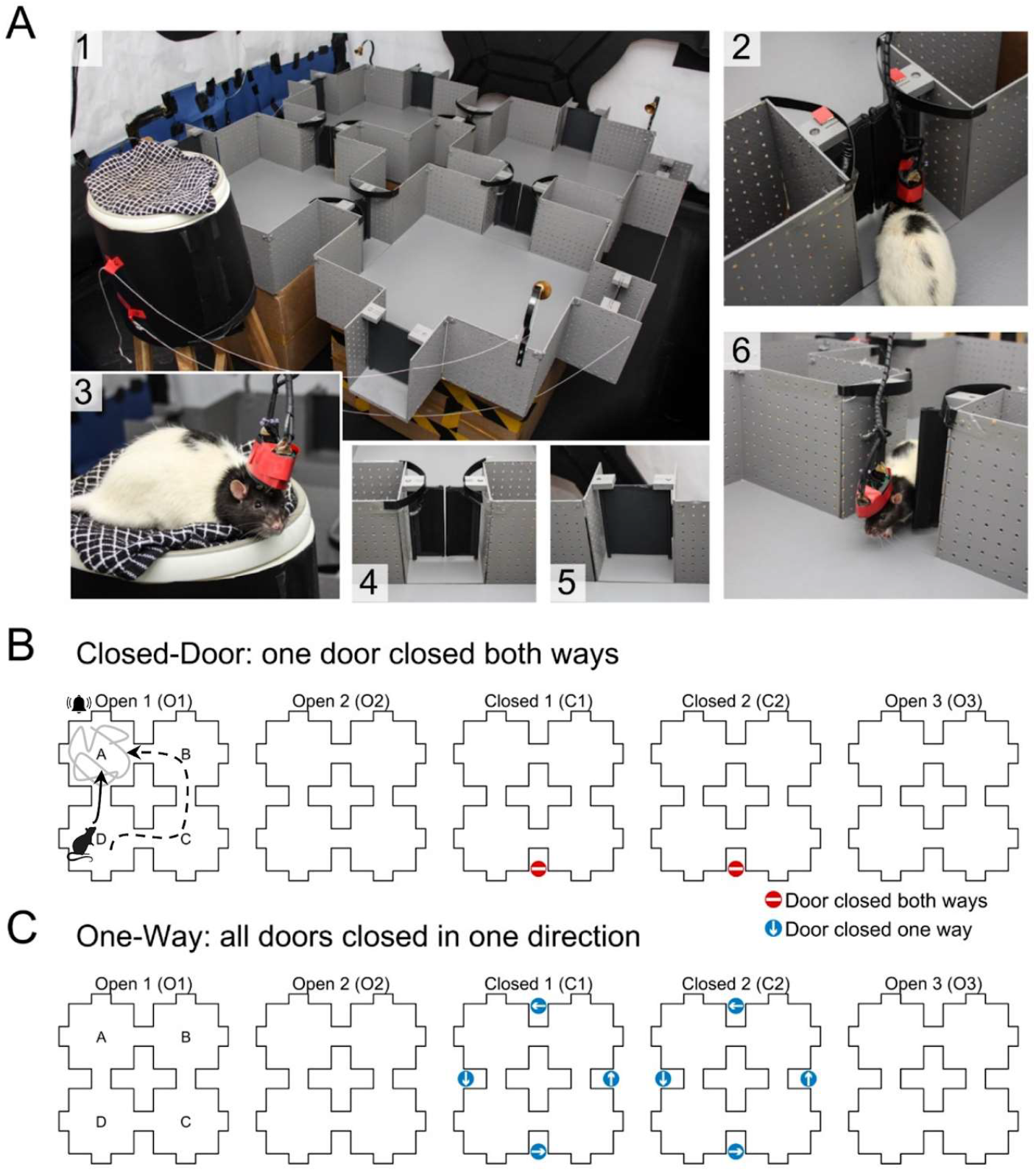
The four-room environment and task protocol. For a detailed protocol see Methods: Task. **(A)** Photographs of: 1) the maze: bells attached to the corner of each box can be activated by pulling on a thread to indicate the next goal box. For every trial, the bell of a chosen box was activated and cereal was thrown in that box. More food was thrown until the rat spent at least 30s in the box, then the next trial started for a total of 12 trials per session. A schematic and photos of the room and distal cues can be seen in Sup. Fig.1; 2) a rat about to push a one-way locked door; 3) the rat resting on the elevated platform, the headcap surrounding the headstage was used to facilitate door-pushing, 4) a normal door, 5) a dummy door with geometry similar to the normal door, 6) a rat passing through an open door. **(B)** Protocol of the ‘Closed-Door’ sequence of sessions. In sessions O1, O2 and O3, all 4 doors were open. In sessions C1 and C2, one of the doors was locked in both directions, indicated by a red stop sign. Possible goal-directed trajectories from D to A are shown in black, a mock foraging trajectory is shown in light grey, these were categorised automatically using duration of trajectories, occupancy and other criteria (*Methods - Behaviour discrimination*). The same goal box list was used for all 5 sessions of a sequence. (C) Protocol of the ‘One-Way’ sequence of sessions. The protocol was similar to Closed-Door except that in sessions C1 and C2 all four doors were locked one-way, allowing only clockwise or anticlockwise transitions. The open direction is indicated by a blue arrow next to the door.

### Rats rapidly learned spatial connectivity changes

To assess whether or not rats were aware of the connectivity of the environment (doors open or locked), we computed the number of door-pushing attempts for each session, normalised per door side and per minute (see Supplementary Video 1 for examples of open or locked door-pushing behaviour). Pushes on each side of a door were manually recorded as distinct events, whether they led to a door-crossing or not (*Methods - Event Flags*). In both the Closed-Door (Fig. 2A, C) and One-Way (Fig. 2B, D) sequences, rats pushed on all doors equally in the first and second sessions (O1 and O2) when all doors were open (see Table 1 for statistics), but significantly reduced their pushes on the locked doors in the third and fourth sessions when the connectivity was changed (C1 and C2). Importantly, although all doors returned to their open state in the 5th session (O3), rats still pushed less on the previously locked doors (Fig. 2A&B, Table 1), indicating long-term memory for door state. In addition, rats pushed significantly less on the locked side of doors in the C1 session of the One-Way condition compared to the Closed-Door condition, suggesting better knowledge of connectivity in One-Way (Table 1).

**Figure 2:**
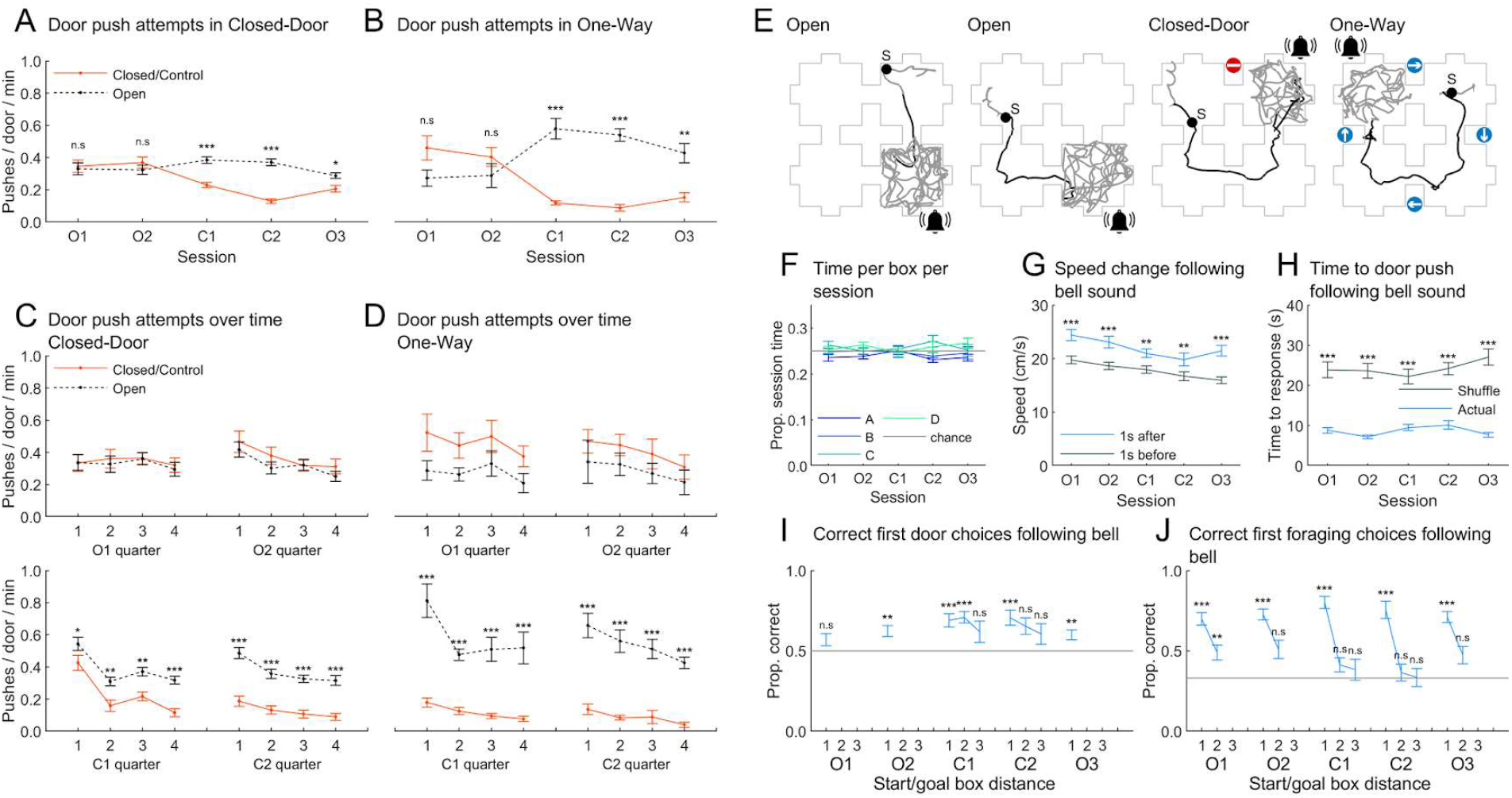
Rat behaviour indicates knowledge of the connectivity and partial knowledge of the task. Rat-averaged data is shown in Sup. Fig.2; Table 1 shows all statistical results. **(A)** Normalised number of door pushes for either the control doors (in O1-3, doors that will be or were locked), locked doors (same doors but in C1 and C2), or open doors, in the Closed-Door sequence, averaged over sessions. **(B)** Same, for the “One-Way” sequence. We usually locked the direction preferred by the rats if they had a bias, which is why the numbers of pushes on the control side seem higher in O1 and O2. **(C)** Same as A but separated by session quarters and without O3. **(D)** Same as C for the One-Way sequence. **(E)** Example trajectories: circular black markers indicate rat position at the time of a bell sound with the bell symbol indicating the origin of the bell sound. Thin grey lines show the outline of the maze. Thick grey lines show trajectory data classified as “foraging”. Thick black lines show “goal-directed” trajectories. **(F)** Average time spent in each box for each session type. For F-J, Closed-Door and One-Way sessions are combined. **(G)** Average movement speed one second before or after the bell sound. Rats significantly increased their speed after the bell sound. **(H)** Average time between a bell sound and the next door push. Rats pushed a door following a bell sound faster than chance (Methods - Response to bell sounds). **(I)** Proportion of correct/optimal first door push after all bell sounds. Chance level is 50% (2 doors per box). Values separated by distance between the current box and the goal box (distance of 1 = adjacent box). Distances of 2 and 3 are only available when a door was locked. (J) Proportion of correct first foraging choices (where the rat forages first after a bell sound). Chance level is 33% when excluding the initial box. Rats always foraged in the correct box for a goal distance of 1, otherwise the proportion was not always significantly different from chance. See Sup. Fig.3 for plots of I and J per sequence type, *Methods - Response to bell sounds and Methods - Correct door pushes and correct foraging*. Here and later, error bars indicate standard error of the mean and * :p < .05, ** : p < .01, ***: p < .001.

**Table 1:**
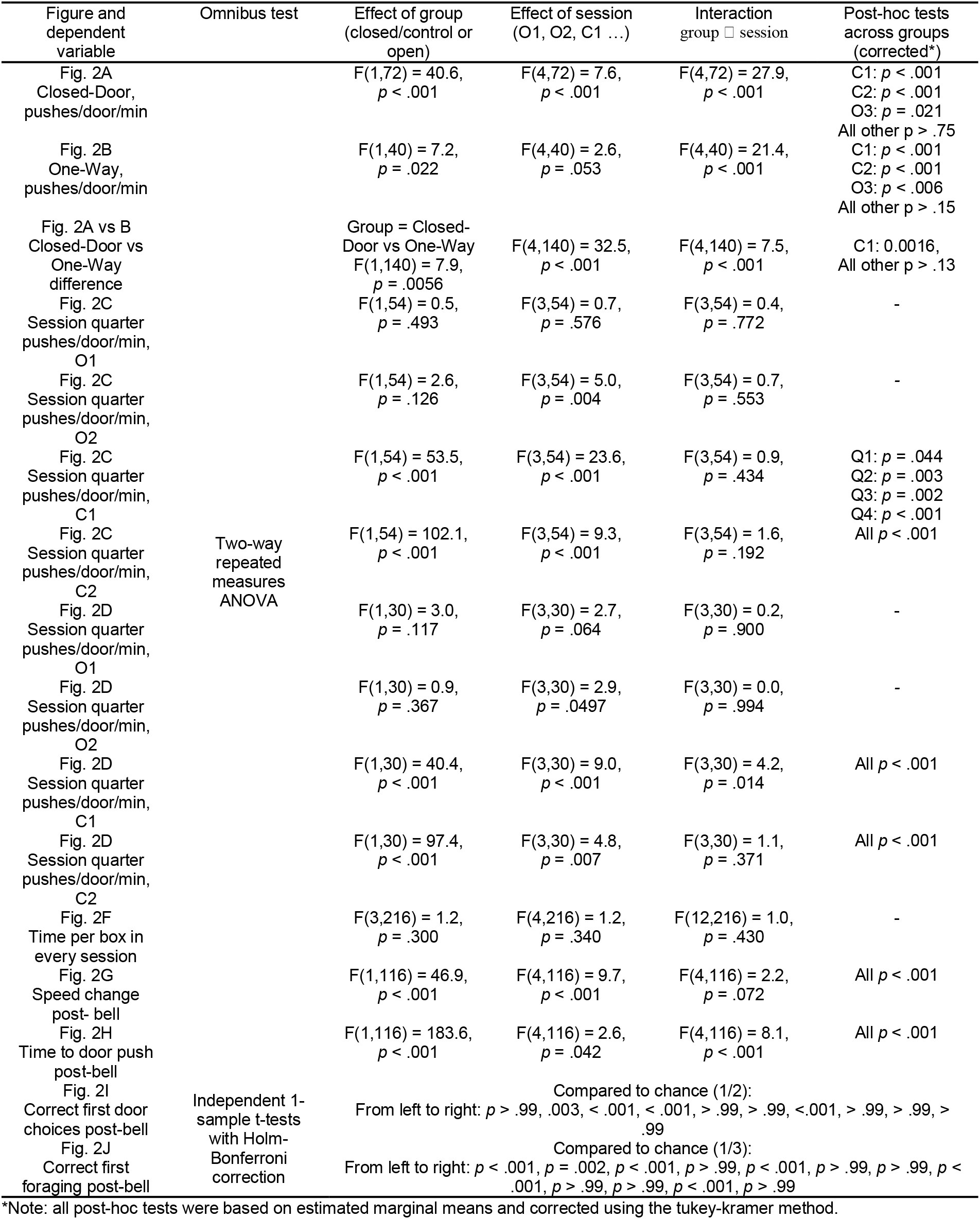
Knowledge of the door status and partial knowledge of the task. Statistical test results corresponding to Fig. 2.

To investigate connectivity learning at a finer timescale, we divided each session into quarters and compared the number of pushes on locked and open doors as before. In both conditions, rats discriminated the locked doors from open doors from the first quarter indicating a very fast learning process (Fig. 2C,D, Table 1). These effects also hold across individual rats (Sup. Fig.2).

### Rats associated the bell with a change in goal but did not always locate its origin

We then asked whether rats were making optimal door choices in the maze and started foraging in the correct location, to evaluate their knowledge of the task and of the global structure of the maze. Our ‘guided foraging’ protocol encouraged rats to alternate between pseudo-random foraging within a box and goal-directed trajectories towards a different goal box, cued by remotely ringing a bell located at the corner of that box. This was done in order to keep the rats engaged, allow for homogeneous spatial sampling and avoid the use of a response strategy - relying on repetitive trajectories - which would not necessarily engage the hippocampus (Packard and McGaugh 1996), but see (Gasser et al. 2020). The two types of trajectories were automatically classified using criteria such as time spent inside a given compartment, occupancy and door-pushes (Fig. 1B, Fig. 2E; *Methods - Behaviour Categorisation*); specifically, any trajectory between boxes was categorised as goal-directed. We note that rats did not seem to see or smell the reward unless it was very close as they would sometimes leave the current box during a foraging phase; we also anecdotally observed detour trajectories (Fig. 2E, right) suggesting the use of a global map to navigate.

Session duration did not significantly vary with session position in the sequence or door condition, either for Closed-Door or One-Way (*p* > .05 in all cases, univariate ANOVA) and rats spent the same time in each box (Fig. 2F, Table 1). Rats’ speed consistently increased following a bell sound (Fig. 2G, Table 1) after which they pushed on a door (whether open or locked) more quickly than would be expected by chance (Fig. 2H, Table 1), confirming that they associated the sound with shifting their current goal to a different box. The door rats first interacted with following a bell sound tended to be the optimal one, but this was not statistically significant for all conditions (Fig. 2I, Table 1). Finally, rats selectively stopped to forage in the rewarded box only significantly more than chance when it was immediately connected to the start box (Fig. 2J). This suggests that rats would often start foraging in the box they entered, regardless of whether it was rewarded or not. This could be because checking for reward took very little effort, or because they were unsure of the goal location. Data separated by sequence type are shown in Sup. Fig.3. In summary, rats seemed aware that the bell sound indicated a new goal and usually, but not always, chose the optimal path to reach it.

Because rats would rarely attempt to push on locked doors, running speed, movement direction and other behavioural parameters known to influence place cell firing would be very different around locked doors compared to open ones when rats were transitioning from one box to another (see average dwell maps in Sup. Fig.4). To control for this, we focused most of the neural analyses on the foraging behaviour where these parameters were consistent across conditions.

### Place cell activity in the 4-room environment

We recorded CA1 extracellular activity from the same 5 rats and 30 sessions used for the behavioural analyses. Only putative place cells were analysed: well-isolated, putative pyramidal cells with spatial firing, active in at least two of the 5 sessions (*Methods - Unit classification and Methods - Firing rate maps and spatial information*). All recordings were histologically confirmed to originate from the dorsal CA1 field of the hippocampus (*Methods - Histology* and Fig. 12). In Closed-Door sequences, we recorded 261 place cells (out of 325 neurons or 80.3%), while in One-Way sequences, we recorded 161 place cells (out of 232 neurons or 69.4%). Table 2 summarises the number of recording sequences, place cells and place fields per rat and sequence type. Cluster quality measures can be found in Sup. Table 2 (*Methods - Cluster quality measures*). Spike plots and rate maps for all place cells are shown in Sup. Fig.10 for Closed-Door and Sup. Fig.11 for One-Way.

**Figure 12:**
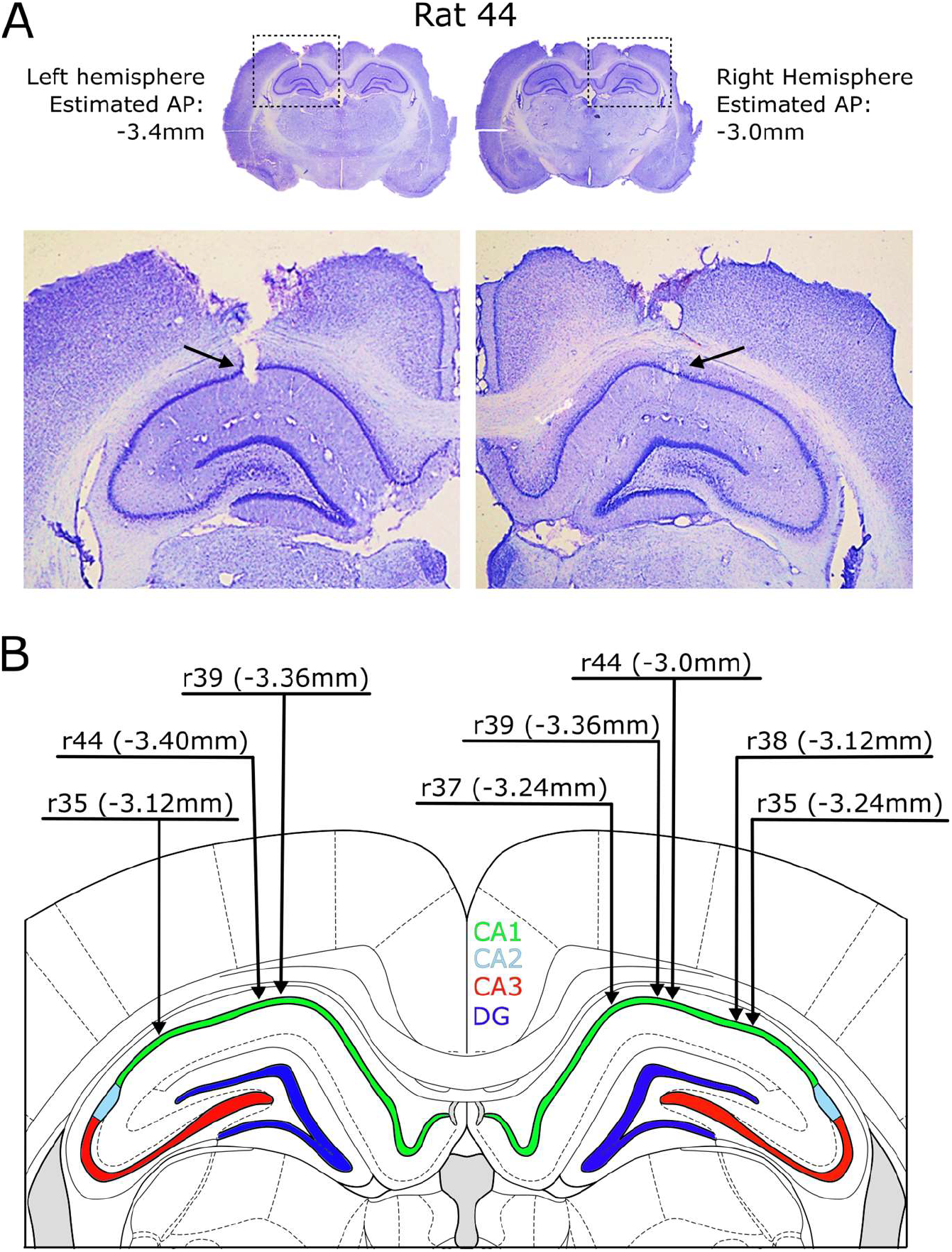
Histological verification of electrode recording sites. **(A)** Top: histological slices for r44 showing the position of the left and right hemisphere electrodes. Text gives the estimated position of the electrodes relative to bregma in the anterior-posterior (AP) axis. Bottom: close-up view of the slices in the region outlined by a dotted line. Black arrows denote the position where the electrodes crossed the CA1 cell layer. **(B)** Brain atlas section with superimposed electrode angles and positions estimated for all animals (black arrows). Arrowheads give the position at which the electrodes crossed the CA1 layer. Text gives the rat number and estimated AP position of the electrode tract. Note that the brain atlas section is for an AP of -3.84mm relative to bregma (the target AP) but estimated APs vary from this position.

**Table 2:** Numbers of sequences, place cell and place field numbers per rat and condition.

| Rat | Number of full sequences |  | Place cells (% of all neurons) |  | Place fields * |  |
| --- | --- | --- | --- | --- | --- | --- |
|  | Closed-Door | One-Way | Closed-Door | One-Way | Closed-Door | One-Way |
| r35 | 6 | 3 | 73 (74.5) | 43 (75.4) | 525 | 267 |
| r37 | 3 | 2 | 29 (80.6) | 26 (83.9) | 91 | 95 |
| r38 | 4 | 2 | 32 (94.1) | 12 (70.6) | 93 | 53 |
| r39 | 3 | 2 | 47 (79.7) | 35 (58.3) | 254 | 358 |
| r44 | 3 | 2 | 80 (81.6) | 45 (67.2) | 412 | 236 |
| Total | 19 | 11 | 261 (80.3) | 161 (69.4) | 1375 | 1009 |

\*Note: Place field numbers here were detected in the session with the highest mean firing rate (the session used when categorising units as putative place cells): thus these represent an upper bound and are not representative of the average session.

The activity of a selected set of example cells for Closed-Door and One-Way sequences is shown in Fig. 3 and Fig. 4, showing the variety of responses to the compartmented space and to connectivity manipulations. Cells had a variety of field numbers, repeating or not. While their spatial activity remained generally stable across consecutive sessions, place fields sometimes appeared and disappeared in a way not specifically linked to the changes in connectivity. These observations are quantified in the following results.

**Figure 3:**
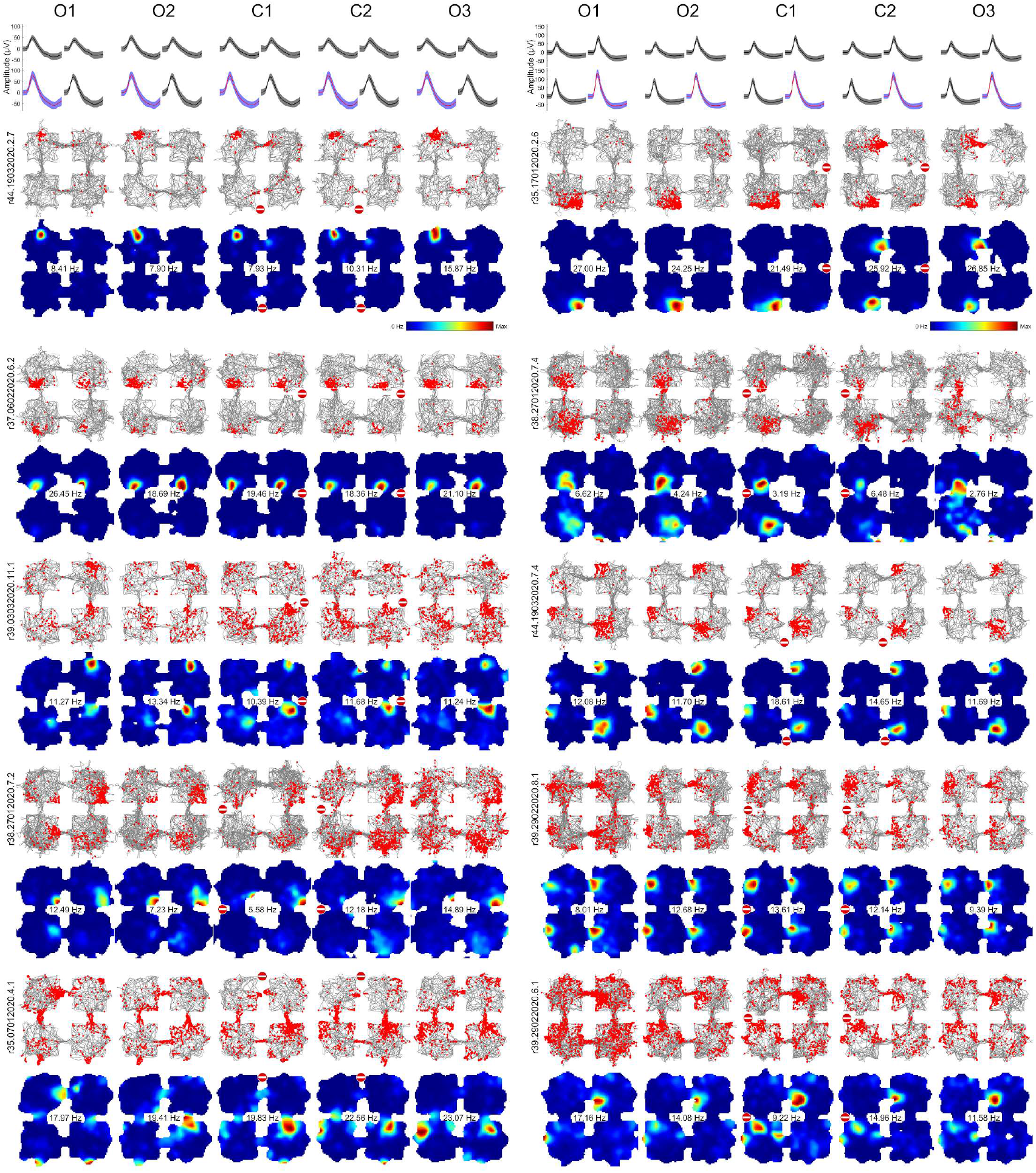
Variety of place cells responses in the Closed-Door sequence. First row: average ± standard deviation of waveform amplitudes on each tetrode channel for the first two cells, the largest amplitude is shown in blue. Below, each row pair shows the spike plots (top; trajectory in grey, spikes in red) or speed-filtered rate maps (bottom; number indicates maximum firing rate; see *Methods - Firing rate maps and spatial information*) for a given place cell recorded in the five sessions of a sequence. Cells were selected to show a range of place field numbers (increasing from 1 at the top to 5 or more at the bottom) and response types across different rats. Closed doors are indicated by signs as in Fig. 1.

**Figure 4:**
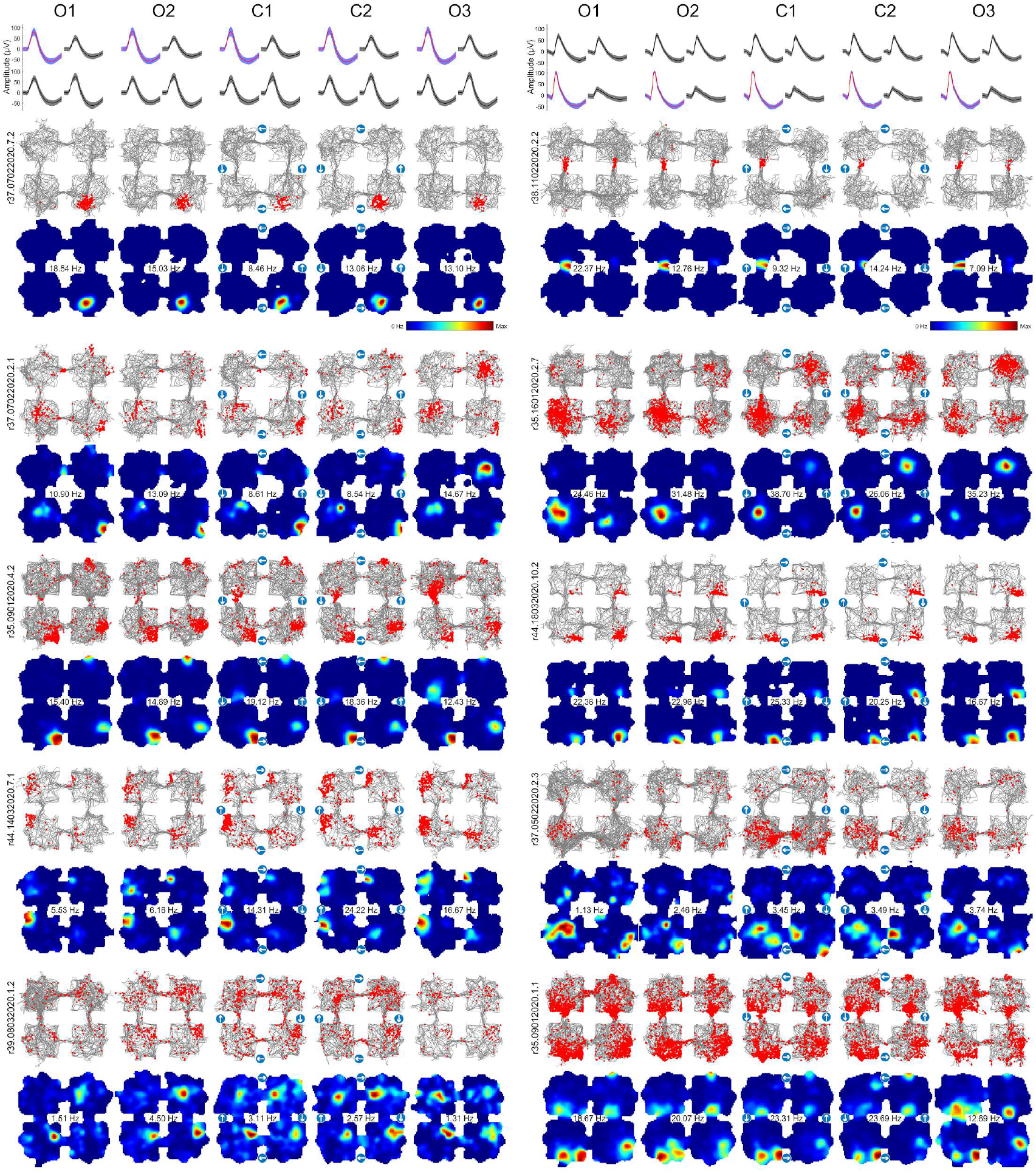
**Variety of place cells responses in the One-Way sequence,** organised as in the previous figure.

### Place cells did not respond to changes in spatial connectivity

Our main hypothesis was that CA1 place cells, as the neural substrate of a spatial cognitive map, explicitly encode the connectivity of environments. For the neural data, we analysed only foraging periods unless otherwise specified. This excludes ‘goal-directed’ periods of transition between boxes as behaviour during these would strongly differ for different connectivity conditions which might in turn spuriously influence place cell firing. This partition of the data was performed automatically (*Methods - Behaviour discrimination*), see Fig. 2E for examples and Sup. Fig.4 for cumulated occupancy maps in the different conditions. Note that the firing location of place cells did not significantly change between goal-directed and foraging, as will be presented later (*Results – Place cells did not spatially remap between task phases*).

We first asked whether place cells globally encoded connectivity status by comparing firing rate maps between sessions, as connectivity changed between sessions Open2 - Closed1 (O2-C1) and Closed2 - Open3 (C2-O3, Fig. 5B&H, see Methods - Rate map correlations). Average correlations between consecutive sessions were always highly positive and O2-C1 correlations, where connectivity changed, were not different from O1-O2 which had identical connectivity (Fig. 5C&I, Table 3 for statistics). Instead, correlations significantly decreased with time (Fig. 5D&J), coherent with past findings (Mankin et al. 2015). We next focused on activity close to the connectivity changes by only using data within a 25 cm radius of the doors (Fig. 5A&G). There were no significant differences between O1-O2 correlations and O2-C1 correlations, either for Closed-Door (Fig.5E, Table 3) or One-Way (Fig.5K, Table 3).

**Figure 5:**
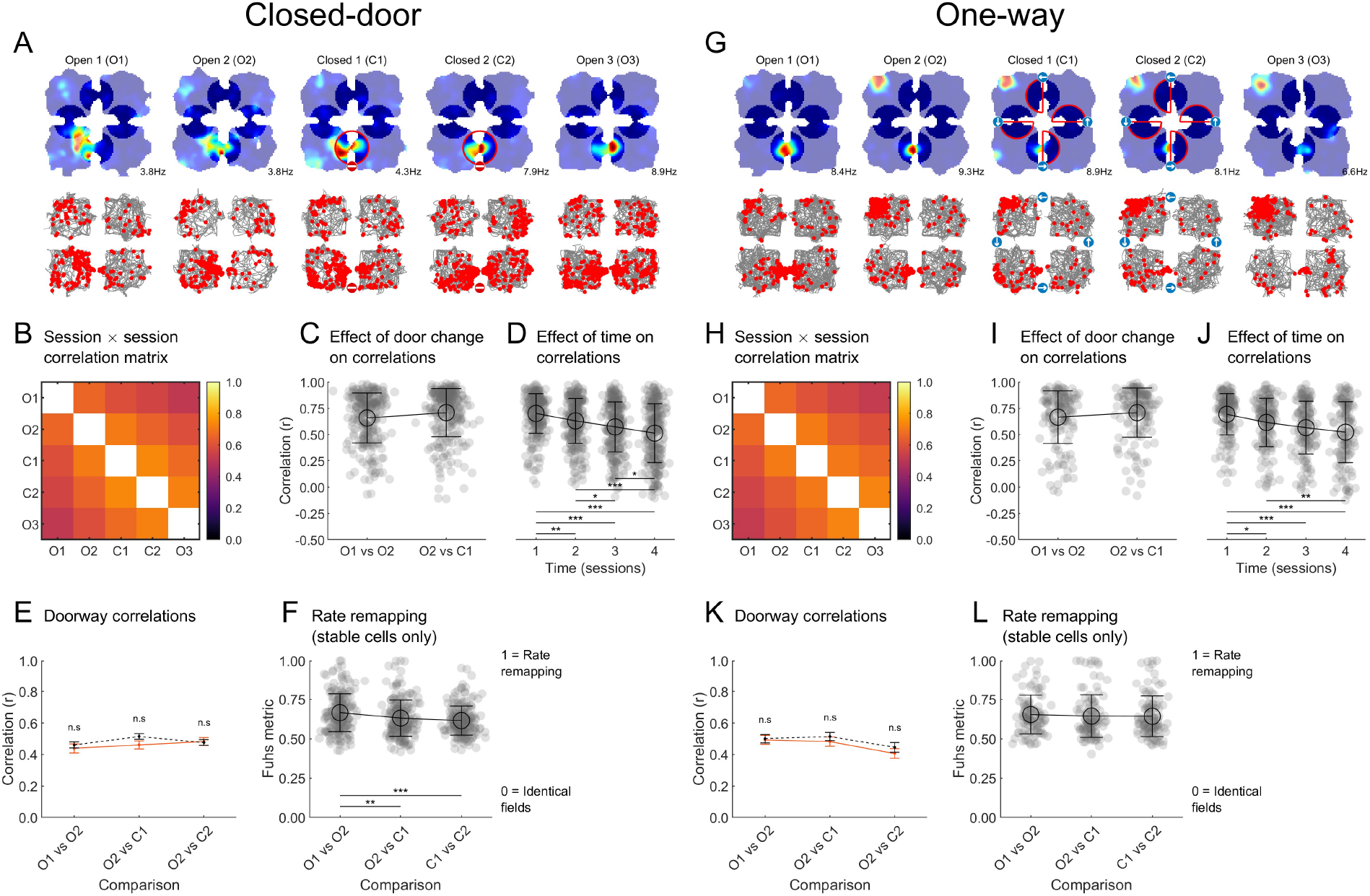
Place cell firing was not altered by changes in the connectivity. See Table 3 for all statistical results. All data uses foraging-only behaviour. **(A)** Example cell recorded in a Closed-Door sequence, presented as in Fig. 3. *top*, foraging rate maps; non-shaded areas show the data used in the doorway correlation analysis in E, red circled areas in C1 and C2 show the region used for the locked door data, the equivalent area is used as control data for O1, O2 and O3, the other regions are used for the open door data. *bottom*, foraging spike maps. Note that even when the door was locked, this cell kept firing on both sides of the door and in the same location. **(B)** Matrix of cell-averaged rate map correlations between all sessions. All values are relatively high (>0.4) but drop as the time between sessions increases (i.e. towards the top right and bottom left corners). **(C)** Comparison of the correlations between two open sessions (O1 vs O2) and one open and one locked (O2 vs C1). Remapping following door closure would trigger a drop in correlations but instead, the two distributions are not different. **(D)** Correlations between rate maps of a recording sequence as a function of the time distance between them, 1 = successive sessions, 2 = separated by 1 session, etc. Correlations decreased over time. **(E)** Correlations of activity within a 25-cm radius of the doors, as indicated in A. No effect was observed, demonstrating that closing a door does not trigger remapping, even locally. **(F)** Rate remapping index between the same session pairs as E; high values indicate increased rate remapping. Values significantly decreased for the comparisons with the connectivity change indicating that there was even less remapping when the connectivity was modified compared to between two sessions of unchanged connectivity. **(G-L)**: same presentation as A-F for the One-Way sequence, with similar results.

**Table 3:** Statistical test results corresponding to Fig. 5.

| Figure and dependent variable | Omnibus test | Effect of group (closed/control or open) | Effect of session | Interaction group □ session | Post-hoc tests across groups (corrected*) |
| --- | --- | --- | --- | --- | --- |
| Fig. 5C<br>Correlations<br>O1-O2 vs O2-C1 | one-way ANOVA | - | $F(1,509) = 6.0$,<br>$p = .015$ | - | - |
| Fig. 5D<br>Correlations over sessions/time | one-way ANOVA | - | $F(3,1017) = 30.4$,<br>$p < .001$ | - | 1v2: $p = .003$<br>1v3: $p < .001$<br>1v4: $p < .001$<br>2v3: $p = .030$<br>2v4: $p < .001$<br>3v4: $p = .020$ |
| Fig. 5E<br>Doorway correlations | two-way ANOVA | $F(1,1238) = 1.6$,<br>$p = .212$ | $F(2,1238) = 1.7$,<br>$p = .190$ | $F(2,1238) = 0.9$,<br>$p = .400$ | - |
| Fig. 5F<br>Fuhs rate remapping metric | one-Way ANOVA | - | $F(2,598) = 11.0$,<br>$p < .001$ | - | O1vsO2 vs O2vsC1:<br>$p = .004$<br>O1vsO2 vs C1vsC2:<br>$p < .001$<br>All other $p > .30$ |
| Fig. 5I<br>Correlations<br>O1-O2 vs O2-C1 | one-way ANOVA | - | $F(1,509) = 6.0$,<br>$p = .015$ | - | - |
| Fig. 5J<br>Correlations over sessions/time | one-way ANOVA | - | $F(3,602) = 13.8$,<br>$p < .001$ | - | 1v2: $p = .022$<br>1v3: $p < .001$<br>1v4: $p < .001$<br>2v4: $p = .005$<br>All other $p > .25$ |
| Fig. 5K<br>Doorway correlations | two-way ANOVA | $F(1,774) = 1.2$,<br>$p = .274$ | $F(2,774) = 4.2$,<br>$p = .015$ | $F(2,774) = 0.2$,<br>$p = .849$ | - |
| Fig. 5L<br>Fuhs rate remapping metric | one-way ANOVA | - | $F(2,330) = 0.2$,<br>$p = .78$ | - | - |

\*Note: all post-hoc tests were based on estimated marginal means and corrected using the Tukey-Kramer method.

Thus, place cells’ firing location did not consistently change with connectivity, either locally or globally. We also examined rate remapping, i.e. change of firing rate without a change of spatial selectivity (S. Leutgeb et al. 2005) in the full foraging firing rate maps using a normalised index (Fuhs et al. 2005). To focus on rate and not position remapping - already assessed by the previous analysis - we excluded cells that were not spatially stable from this analysis (*Methods - Firing rate remapping*). The Fuhs metric equals 0 for identical fields and approaches 1 when fields are mis-aligned or differ in firing rate (Fuhs et al. 2005). Rate remapping did not increase when the connectivity changed, i.e when comparing O2-C1 to O1-O2 (Closed-Door; Fig. 5 F & L), indicating that place cells did not encode a change of connectivity in their firing rates.

### Individual place cells did not significantly remap due to the change in spatial connectivity

Place cell remapping at the individual neuron level might not have been visible in the previous population-level analyses; perhaps a small, undetected subpopulation of cells was encoding the door change. To address this we computed individual remapping indices between consecutive sessions for each cell. This was done by comparing the cross-session correlation values between foraging maps to a distribution of correlations obtained from shuffled maps; correlations above the 95th percentile of the shuffle distributions indicated stable cells (*Methods - Individual remapping between sessions*). Each cell’s remapping pattern was reduced to a series of numbers where 0 represents no remapping and 1 represents above-chance remapping (inset Fig. 6 B). For example, a cell remapping due to changes in connectivity would have a *0101* pattern (remapping each time the connectivity changes); given the uncertainty of connectivity knowledge for the last session (rats sometimes still avoiding pushing on the previously locked door; Fig. 2 A,B) we also considered cells with a *0100* pattern as putative connectivity cells.

**Figure 6:**
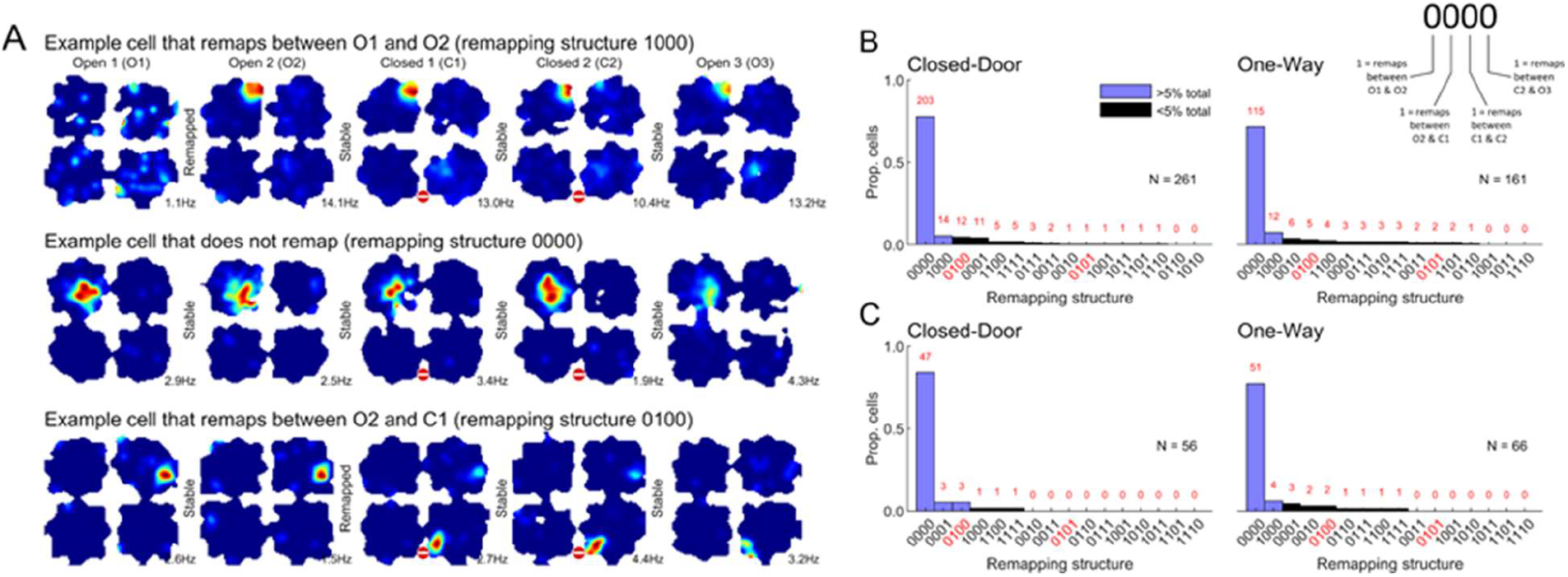
Individual place cells do not preferentially remap following connectivity changes. **(A)** Example rate maps (foraging only) from which the individual cell remapping is assessed. This is done by comparing the map correlations of a given place cell between two consecutive sessions to those from shuffled (random cells) maps, and if the data value is lower than the 95^th^ percentile of the shuffle data, the cell is considered to remap. (B) Histograms for each protocol showing the distribution of remapping patterns for all place cells (*1* = remapping, *0* = significantly stable, according to the key given in B). Red x-axis labels highlight the 2 conditions where remapping could be attributed to a change of connectivity. Few cells individually remapped selectively due to the connectivity change (<5% of cells). (C) Similar to B but only for cells with one of their fields with their centroid closer than 25cm of the locked door(s); the same pattern of results was found.

In the Closed-Door condition, only 13 cells (4.5% of place cells) had a pattern indicative of remapping due to connectivity changes (12 cells for *0100* and 1 for *0101*). The majority of cells were significantly more stable than chance (203 cells for *0000*). Similar results were obtained in the One-Way sequence with even less connectivity-encoding cells (Fig. 6B, right). These numbers were equivalent to - or lower than - those of cells spontaneously remapping between O1 and O2, thus, they do not appear to reflect any connectivity-specific remapping. To exclude the possibility that the few connectivity-encoding cells are most of the cells active next to an affected doorway, we restricted the results to cells with a field in the vicinity of locked doors (Fig. 6C) and found similar results (3/56 connectivity-remapping cells for Closed-Door, 2/66 for One-Way).

### Spatial connectivity changes did not alter place field number, location, size or firing rate

We next focused on individual place fields (detected on speed-filtered, foraging maps *Methods - Place fields*). All detected place field centroids are shown in Fig. 7A&E. The number of active place fields (Fig. 7B,F) or fields in the vicinity of the locked doors (Fig. 7C,G) did not change across sessions for either the Closed-Door or One-Way sequences (see Table 4 and Table 5 for all statistics).

**Figure 7:**
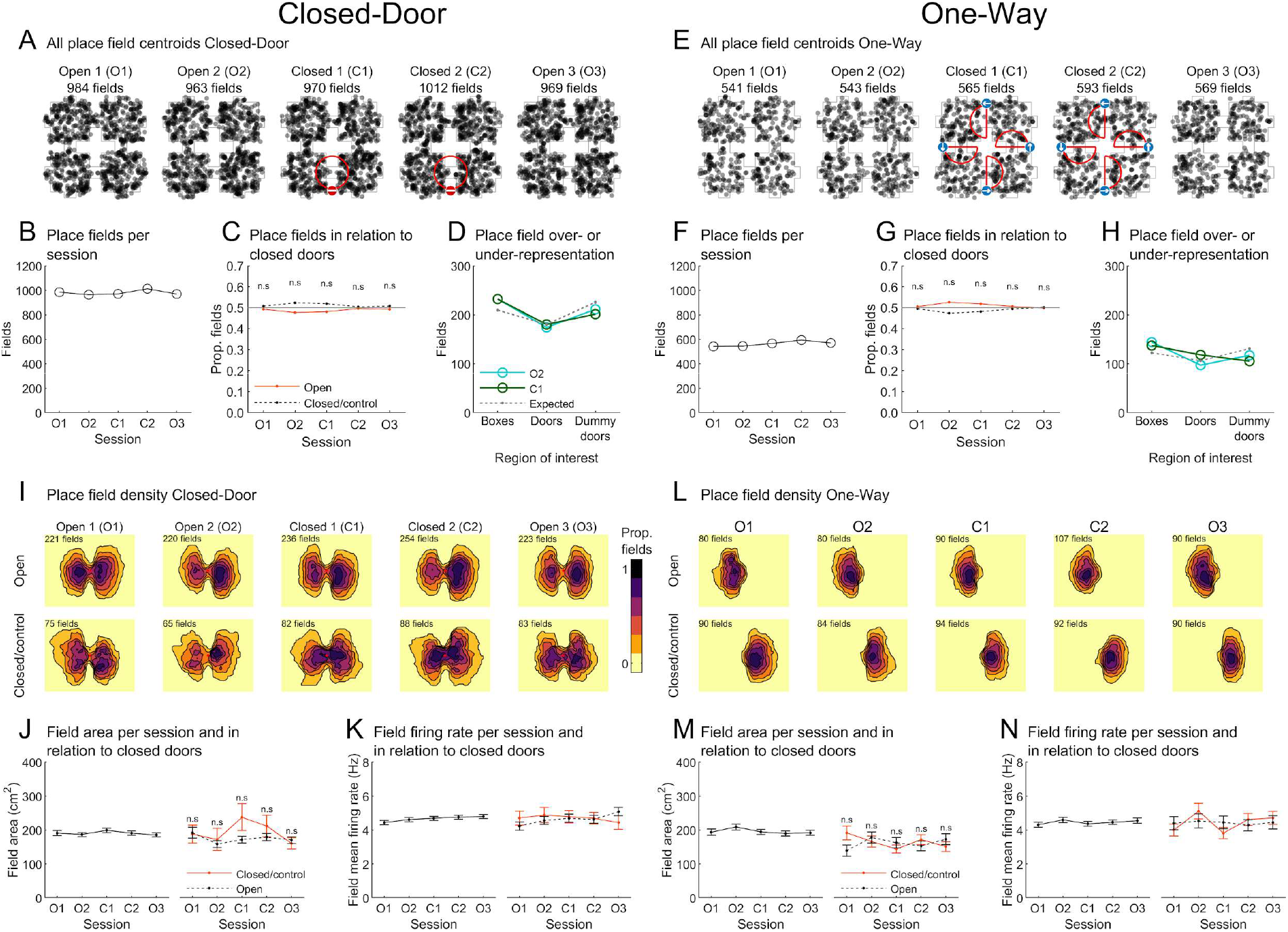
Absence of door overrepresentation or change in place field number, position or size around the locked door(s). See Table 4 and Table 5 for all statistical results. **(A)** Spatial distribution of all place field centroids for each session of the Closed-Door sequence. Numbers of place fields are given at the top. **(B)** Total number of place fields detected in each session. **(C)** Of place fields around a door (centre of mass <25 cm to door), proportion around closed or open/control doors. No effect was found. **(D)** Number of detected place fields around doors, dummy doors or inside a box. The number of expected fields, assuming a uniform distribution, is also shown for each area. Fields are not different from being uniformly distributed. **(E-H)** Same as A-D but for the One-Way condition. **(I)** Cumulated contour maps of place fields around a door (centre of mass <25 cm to a door) with number of place fields shown for each plot. **(J)** Field area either for all data (left) or only in the vicinity of a door (right). There was no significant difference of field size between locked and open doors, although there was a trend towards an expansion of place field size around the locked door in the sessions of interest (C1 & C2). **(K)** Mean infield firing rate across sessions for all data (left) or only door place fields (right). No significant difference or session effect was observed. **(L-N)** Same as I-K but for One-Way with similar results.

**Table 4:**
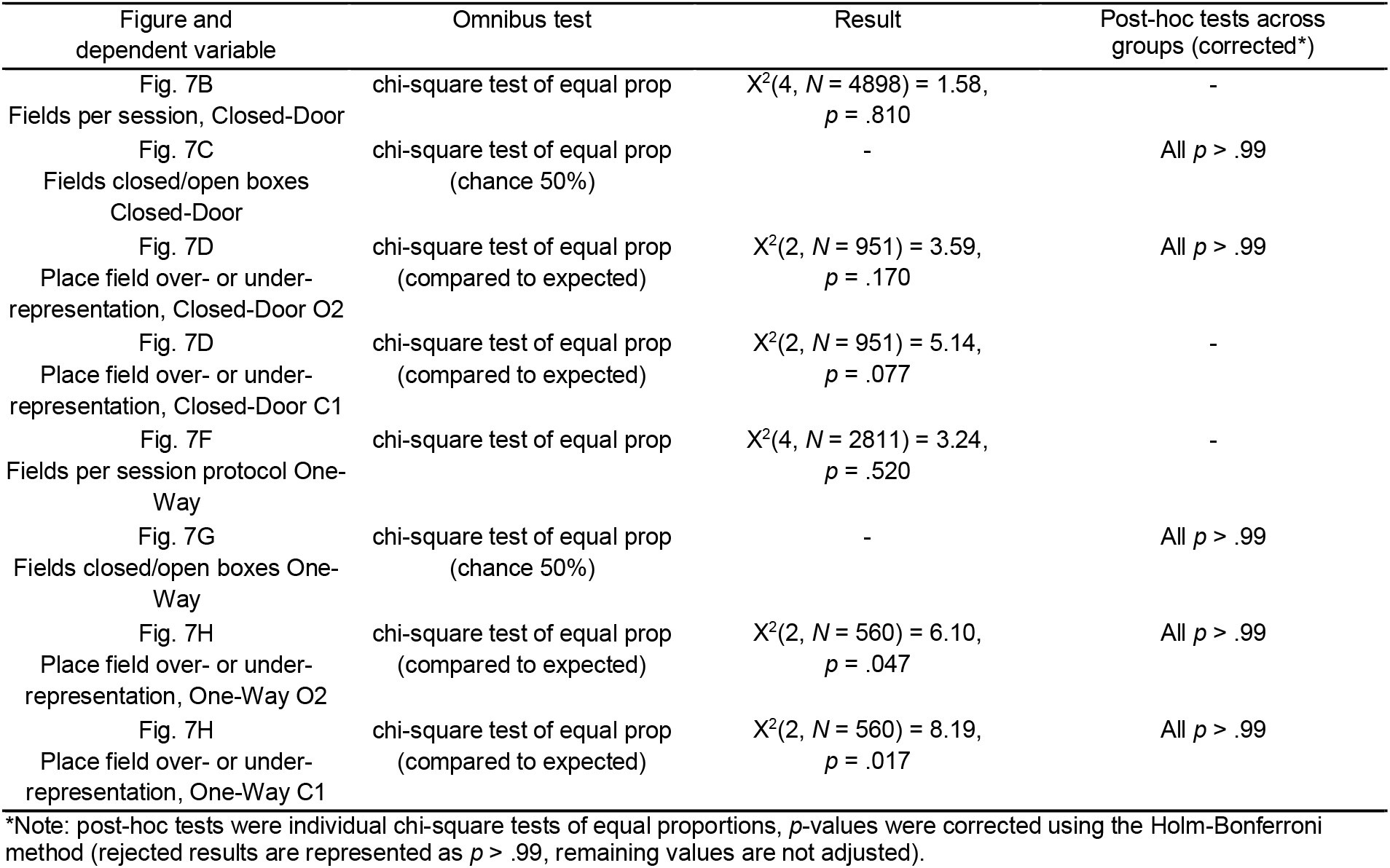
Statistical tests results corresponding to Fig. 7 panels B, C, D, F, G, H

**Table 5:** Statistical tests results corresponding to Fig. 7 panels J, K, M, N.

| Figure and dependent variable | Omnibus test | Effect of session | Effect of group | Interaction<br>group × session |
| --- | --- | --- | --- | --- |
| Fig. 7J left<br>Field area between<br>sessions, Closed-Door | one-way ANOVA | $F(4,4893) = 0.7,$<br>$p = .590$ | - | - |
| Fig. 7J right<br>Field area per session per<br>doorway, Closed-Door | two-way ANOVA | $F(4,1537) = 1.8,$<br>$p = .120$ | $F(1,1537) = 2.8,$<br>$p = .100$ | $F(4,1537) = 1.4,$<br>$p = 0.230$ |
| Fig. 7K left<br>Field frate between<br>sessions, Closed-Door | one-way ANOVA | $F(4,4893) = 1.4,$<br>$p = .227$ | - | - |
| Fig. 7K right<br>Field frate per session per<br>doorway, Closed-Door | two-way ANOVA | $F(4,1537) = 0.3,$<br>$p = .910$ | $F(1,1537) = 0.1,$<br>$p = .770$ | $F(4,1537) = 0.8,$<br>$p = 0.510$ |
| Fig. 7M left<br>Field area between<br>sessions, One-Way | one-way ANOVA | $F(4,2806) = 0.9,$<br>$p = .470$ | - | - |
| Fig. 7M right<br>Field area per session per<br>doorway, One-Way | two-way ANOVA | $F(4,887) = 0.4,$<br>$p = .840$ | $F(1,887) = 0.1,$<br>$p = .710$ | $F(4,887) = 1.9,$<br>$p = 0.110$ |
| Fig. 7N left<br>Field frate between<br>sessions, One-Way | one-way ANOVA | $F(4,2806) = 0.7,$<br>$p = .610$ | - | - |
| Fig. 7N right<br>Field frate per session per<br>doorway, One-Way | two-way ANOVA | $F(4,887) = 1.1,$<br>$p = .360$ | $F(1,887) = 0.0,$<br>$p = .900$ | $F(4,887) = 0.9,$<br>$p = 0.440$ |

Note: no multiple comparisons across groups are reported here because no omnibus tests indicated a significant effect of group.

As the doors must have a particular significance for the animal in this environment, they could have been specifically represented by place cells, for example with ‘door cells’ that would be active only around all the doors but nowhere else. Additionally, cells could represent the connectivity status of a doorway, for example by firing only at locked doors. However, we did not observe this using a cell-by-cell observation of individual rate maps, even within the population of cells rejected for the analysis (non-place cells, data not shown). We did observe some cells encoding two parallel doors (see for example top right example of Fig. 4); interestingly, these observations never extended to non-parallel doors, but sometimes included dummy doors, indicating a probable geometric origin (see also prior to last example on the right of Fig. 3). Alternatively, place fields might over-represent doors, as reported previously for entryways into compartments (Spiers et al. 2013). We compared the density of place fields located around the doors, dummy doors and compartments to the number expected if fields were distributed uniformly throughout the mazes and found that place fields did not significantly overrepresent any of these locations (Fig. 7D, H, *Methods - Place field overrepresentation*).

We also investigated whether connectivity changes could affect the size of place fields (*Methods - Place field firing rate and area*), illustrated in Fig. 7I and L by contour plots of the cumulated density of all place fields with a centroid less than 25cm from the doors. When looking at all fields, sizes did not significantly change across sessions and fields next to a locked door did not change in size more than those next to unchanged doors (Fig. 7J&M). However, there was a non-significant trend for increased place field area around the locked door during session C1 in the Closed-Door condition only, but this was not reproduced in the One-Way condition. Similarly, there was no difference in local firing rate (Fig. 7K&N). Finally, we investigated if place fields moved in response to a locked door by tracking the absolute shift of their centroids across sessions. Similar to previous results, fields were found to shift more between O1 vs O2 (no connectivity change) than between O2 and C1 (connectivity change), leading us to conclude that connectivity changes do not specifically induce field displacement (see *Supplementary Methods - Field distance to doorway* and Sup. Fig.5).

Finally, we asked whether fields generally extended across doors and whether this changed when the doors became locked (*Methods - Place field extent across doors*). Focusing only on fields in the vicinity of changing doors (weighted centroid less than 25cm away from a door that is or will be locked), we computed a ‘bridge index’ as the difference between the area on each side of a door as a fraction of the total area (Fig. 8A). This index ranges from 0 (100% of a field is on one side of a door) to 1 (50% of a field is on each side of a door). For visualisation only, we also extracted the length of the field extent on the axis orthogonal to the door axis (Fig. 8B, E). For this analysis exceptionally, we used the combined foraging and goal-directed firing rate maps to allow for maximal detection of a possible effect and ensure that the same field would not be artificially divided because of missing data at the door. In both configurations, we found that the vast majority of place fields did not extend through a doorway. Instead, most fields were limited to within a single compartment as indicated by very low bridge indices (Fig. 8C, F), the majority of which were zero in both configurations (Fig. 8D, G). More precisely, there were no significant differences in bridge index between sessions (Closed-Door and One-Way: X^2^(4,406) = 1.6, *p* > .80 and X^2^(4,917) = 1.7, *p* > .70; Kruskal-Wallis tests) indicating that the number of fields extending through a door remained constant when the door was locked (Fig. 8C, F). The proportion of fields exhibiting a bridge index of 0 in O2 (66.2%) and C1 (69.1%) did not differ significantly (z = -0.39, *p* = .70, z-score test of two population proportions, Fig 8D). Similarly, in One-Way, these proportions were not different between conditions either (74.7% in O2, 69.6% in C1, z = 1.06, *p* = .29, z-score test of two population proportions, Fig 8G).

**Figure 8:**
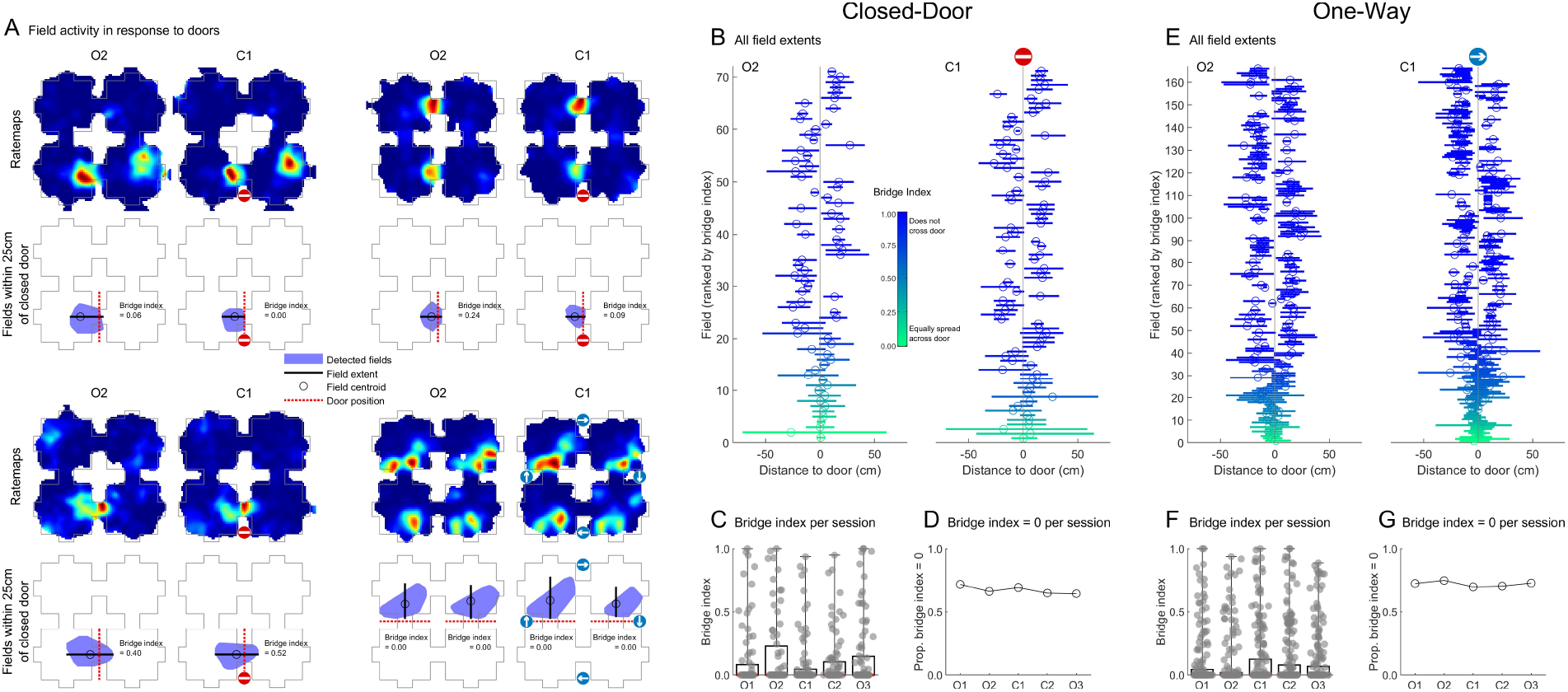
Most place fields do not extend across doors and field extent is not affected by closing the doors. All data (foraging and goal-directed firing rate maps) were used for these analyses. **(A)** Example place cells recorded in the One-Way (bottom right) and Closed-Door (top row, bottom left) conditions with a place field within 25cm of a door that was locked in C1. The cell’s firing rate maps for O2 and C1 are given, below these are schematics of the maze with place fields represented by a blue polygon. Only fields with a centroid within 25cm of a manipulated door are shown and analysed. Doorways within 25cm of a field centroid are represented by a dashed red line. Text gives the bridge index of each field. **(B)** For visualisation, all fields within 25cm of the locked door in the Closed-Door condition. Lines show the extent of place fields along the axis perpendicular to the door, circles give the centroid of the field along this axis. Fields are sorted from bottom to top from a high bridge index (extending across the door) to a low bridge index (entirely in one compartment). **(C)** Bridge indices for these fields in all sessions. In most cases, fields did not cross the nearest doorway and their bridge index was close to zero. The bridge index did not significantly vary between O2 and C1. **(D)** The total proportion of fields with a bridge index of 0 in each session. No change **(E-G)** Same as B-D but for the one-way condition. In this condition all four doors were locked in one direction, thus all fields within 25cm of any door were used. The arrow indicates the open direction in C1.

In summary, our results so far indicate an absence of detectable encoding of changes in spatial connectivity by CA1 place cells. While we have shown that rats were indeed aware of the connectivity of the environment, these results could be explained by a lack of ability to represent each compartment differently from the others (in other words, place cells encoding space only locally and rats confusing boxes) or, since we excluded goal-directed trajectories from most of the analyses, by an effect only visible in those trajectories. These possible explanations are explored below.

### Place cells encoded position in a global reference frame

In previous experiments that had 2 or 4 connected compartments with identical orientation and geometry, almost all place cells appeared to have firing fields in the same local space in each compartment (Skaggs and McNaughton 1998; Spiers et al. 2013; Grieves et al. 2016). In the conditions where such place field repetition (or local encoding) was observed, rats were unable to discriminate between compartments as assessed by an odour discrimination task (Grieves et al. 2016). If place cells encoded position locally in our paradigm, not discriminating compartments, this could interfere with any encoding of compartment-specific connectivity. Additionally, knowing whether place cells encode position in a global or local reference frame (or perhaps both simultaneously) in our paradigm would provide insights into the inputs used by place cells.

To test the type of coding used by place cells in our experiment we first looked at the number of place fields per cell: in our experiment, fewer than 50% of the place cells had 4 or more detected place fields (Fig. 9A) indicating that place field repetition was likely uncommon. Using an analysis that did not rely on place field detection criteria, we computed the correlation between every box in each session, averaged over all cells (Fig. 9B and Table 6 for all statistics of this figure). The observed diagonal bands of high correlations indicate that representations of boxes were highly self-similar across sessions but not between different boxes (Fig. 9B, Table 6). Correlations were also not higher for adjacent boxes than diagonally opposite ones (Fig. 9C, D, Table 6; see *Methods - Firing rate maps and spatial information*).

**Figure 9:**
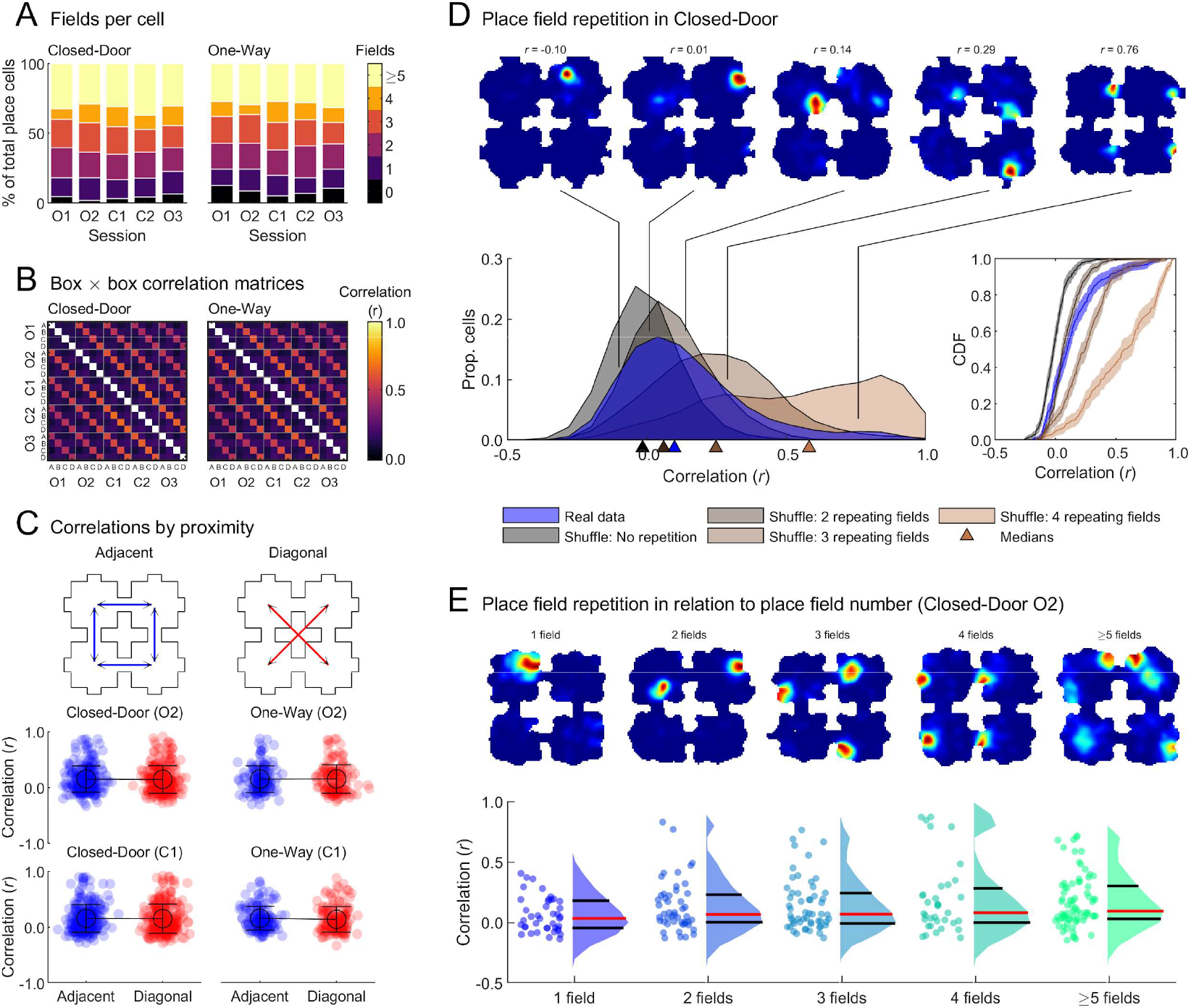
Most place cells do not repeat in the 4-room maze. All statistics can be seen in Table 6. **(A)** Percentage of place cells grouped by number of detected place fields. A perfect local coding would be evidenced by most cells having 4 place fields (or multiples of 4). Instead, most cells have fewer than 4 fields. **(B)** Cross-box correlation matrices averaged over all place cells, computed on speed-filtered, foraging maps, separated by sequence type. Higher values on the diagonals demonstrate that activity is more similar for same over different boxes. **(C)** Paired rate map correlations either for adjacent or diagonal boxes showing that boxes closer in space (adjacent) do not have more similar representations than boxes that are farther away (diagonal). **(D)** D-E show data only for the Closed-Door condition, the One-Way condition can be seen in Sup. Fig.7. *top*, example rate maps (all data, speed-filtered, red indicates high firing rate, dark blue indicates low firing rate) with indicated correlation values on the data (blue) distribution. *Bottom left*, blue distribution represents real data cross-box correlation values for session O2. All other distributions represent different shuffles designed to simulate the correlations we would expect if cells did not have any repeating fields or repeated fields in 2, 3 or 4 compartments. Medians for all distributions are indicated by corresponding markers along the x-axis. *bottom right*, same data as D but showing the empirical cumulative distribution functions, shaded areas represent lower and upper confidence intervals. The data most closely match the distribution expected if cells had 2 repeating fields on average. (E) only for session O2, top, example rate maps for each category of place field number (as described in D). *bottom*, box correlations scores for all data, separated by number of fields; dots shown individual place cell data, the data distribution is shown as violin plot, with median in red and quartiles in black. High correlations can be observed for only a small population of cells.

**Table 6:**
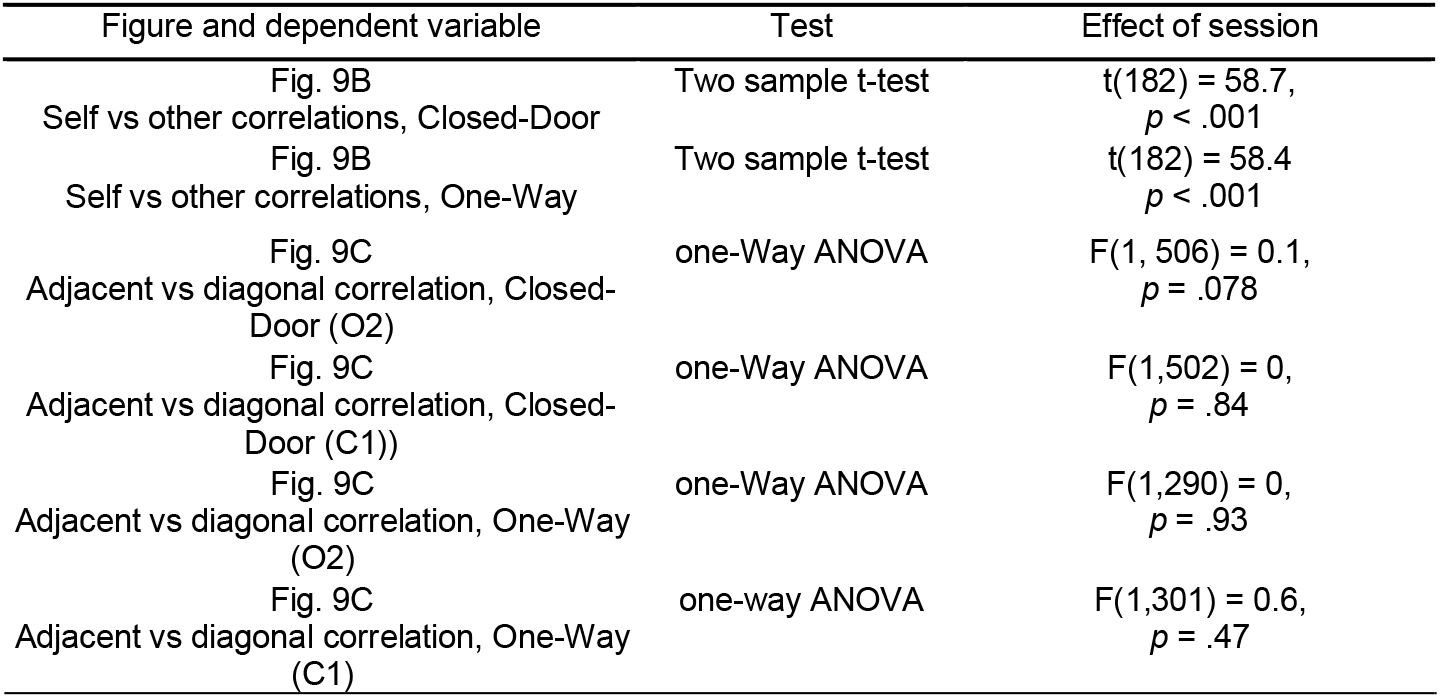
Statistical test results corresponding to Fig. 9.

To further quantify the degree of place field repetition, we computed the average cross-correlations between rate maps of boxes (*Methods - Place field repetition*) and compared them to cross-correlations from shuffled data (Sup. Fig.6). We first observed that the data distribution was skewed towards low values (all medians between 0.07-0.10) in contrast to past experiments reporting place field repetition where cross-compartment correlation distributions were strongly skewed towards high values (>0.5) (Spiers et al. 2013; Grieves et al. 2016; Harland et al. 2017). Observed values differed from the shuffled distributions in all connectivity conditions (*p*<.0001 in all cases), showing that there was more place field repetition than chance, and they did not differ between session types (open or locked) (*p*>.05 in all cases, two-sample Kolmogorov-Smirnov tests) in line with our other results finding no effect of a connectivity change. However, observed medians did not exceed the 95^th^ percentile of the corresponding shuffles and most cells (64.7%) had a cross-box correlation lower than this value. These results suggest that place cells represented compartments more similarly than chance, but nowhere near to the same degree as other multicompartment experiments. To investigate this further we designed shuffles to simulate the correlations expected under different outcomes: place cells representing all compartments uniquely or representing 2, 3 or all 4 compartments identically (*Methods - Place Field Repetition*). We found that the data median (0.10) was closest and extremely similar to the value expected if cells represented two compartments similarly on average (0.06); this distribution was also the only one that did not differ significantly from the observed values (Fig. 9D; *p*>.05, *p*<.001 in all other cases, two-sample Kolmogorov-Smirnov tests).

Did many cells actually repeat their fields in two compartments, or were these values skewed towards high correlations by a few cells repeating in multiple compartments? To investigate this, we classified cells according to their number of detected place fields and plotted their cross-box correlation values (Fig. 9E, using only data from O2 session in Closed-Door). We observed a bimodality in all the distributions with more than one field, revealing a small subpopulation of cells with repeated spatial firing in each category. The results for the One-Way sequence are very similar and are shown in Sup. Fig.7. These results indicate that place field repetition is not an all-or-nothing phenomenon but is instead on a continuum: while the majority of recorded place cells did not repeat, some did in varying subsets of the four boxes. This stands in contrast with past experiments in compartmented spaces where most place cells had repeating fields in all 4 compartments, with occasional cells not fully repeating (Spiers et al. 2013; Grieves et al. 2016). Interestingly, cells exhibiting repeating fields were often co-recorded with non-repeating ones (see for example the bottom right two examples of Fig. 3 from r39, or the 3rd from the top and the bottom right cells from r35 in Fig. 4) and some cells could even have repeating as well as non-repeating fields (see fourth cell in Fig. 9D or third cell in Fig. 9E). To determine if there were enough non-repeating place fields to allow for a fully global coding of space, we next attempted to decode position from place cell firing using Bayesian decoding.

To assess whether place cells indeed implemented a global coding of space, defined as differentiating boxes as well as positions within them, we asked if their activity discriminated enough between boxes to allow for an accurate decoding of individual box quadrants using Bayesian decoding (Kaefer et al. 2020; Zhang et al. 1998), see *Methods - Bayesian decoding of box quadrants*. Briefly, each box was divided in 4 identical quadrants (Fig. 10 A) and rate maps using all data (foraging and goal-directed) from simultaneously-recorded place cells were used to train a Bayesian memoryless decoder. Only sessions with 15 or more simultaneously recorded place cells were used (n = 9 for Closed-Door and n = 3 for One-Way). To avoid overfitting (van der Meer, Carey, and Tanaka 2017), we used the maps from O1 to decode the instantaneous firing in O2, and maps from O2 to decode firing in C1. A mean ‘confusion matrix’ (probability of decoding to a box quadrant given the actual box quadrant) was computed for each type of decoding (O2 Closed-Door, O2 One-Way, C1 Closed-Door, C1 One-Way). We asked if decoding to the correct quadrant in the correct box was more likely than chance and if other possibilities such as decoding to the correct box but incorrect quadrant, or the correct quadrant in incorrect boxes, were also more likely than chance - the former would indicate a quadrant-scale resolution of spatial coding while the latter would indicate place field repetition. We first observed a clear diagonal in all the confusion matrices (Fig. 10 B), indicating accurate decoding to the correct box and quadrant. We also did not observe other diagonals of high-value in the matrices, which would have indicated decoding of the correct compartment in an incorrect box, i.e., local coding of space. We then computed the average decoding probability values for the different possibilities mentioned above as well as chance probability: only the correct quadrant, correct box group was clearly above chance (Fig. 10C). Individual session confusion matrices can be found in Sup. Fig.8 demonstrating the reproducibility of this pattern for most sessions. This result, taken together with our previous analyses, converge towards a clear finding of encoding of global position by the place cell population in the 4-room environment.

**Figure 10:**
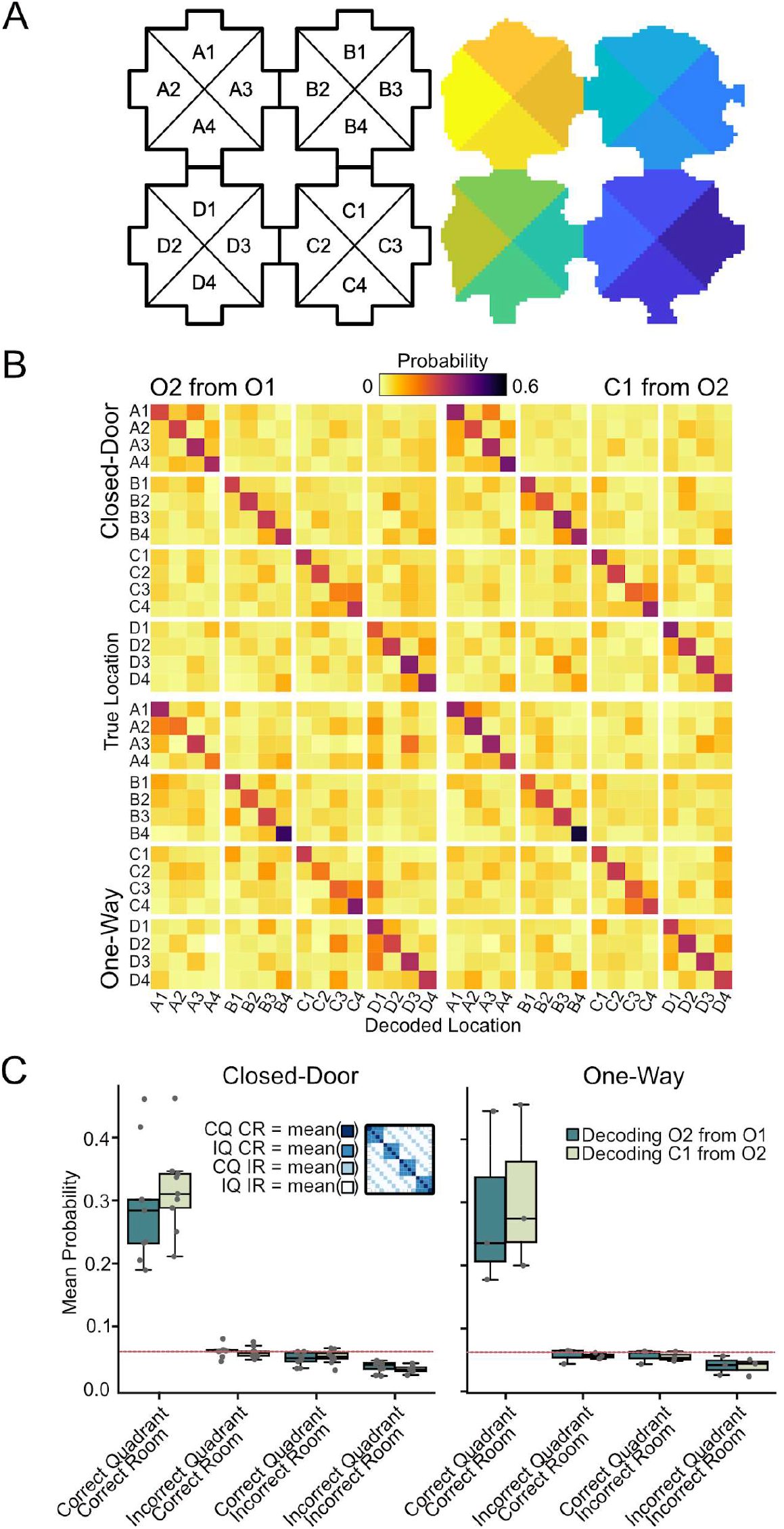
Global position can be decoded from place cell firing using Bayesian decoding. **(A)** Left: schematised division of rooms and quadrants. Right: example discretisation in binned position data. **(B)** Mean confusion matrices from sessions with at least 15 simultaneously recorded place cells from Closed-Door (n = 9) and One-Way (n = 3) conditions. Separate matrices are shown for decoding of positions in session O2 using O1 training rate maps and decoding of positions in C1 using O2 training rate maps. (C) Average decoding performance to different types of quadrants / boxes. Individual points are average probabilities per quadrant for each response category from an individual session. Red line indicates chance performance (1/16). The average probability of decoding to a correct quadrant in the correct box appears much higher than chance, by opposition to all other decoding possibilities which are at or below chance level. Connectivity status did not affect the probability of decoding to the correct quadrant. Inset: schematic of individual session confusion matrix and the matrix cells that are averaged together for each response category.

Additionally, to detect any effects of connectivity changes on decoding, we compared the probability of decoding to the correct quadrant, correct box in the O1-O2 condition to the same probability in O2-C1 condition, for Closed-Door, reasoning that if the activity in the locked condition was different from activity in the open one, the performance of the decoder would drop when using O2 maps to decode C1 data (Fig. 10C). However, we found no significant difference (t(8) = -1.60, *p* = .147, paired t-test), confirming our earlier findings that connectivity changes do not affect place cell firing even at the population level. No test was run on One-Way given the small sample size (3 sessions) but the data distribution appears similar.

In summary, this second set of analyses demonstrates that the majority of the population of place cells represented position in the 4-room environment globally, with a unique map for each room. In addition, we see a small subset of cells expressing repetitive place fields in all boxes. This suggests that global and local encoding of location are not mutually exclusive, and that both codes can be simultaneously supported by the hippocampus, although the global code might be more prominent as in the present case. These results also confirm that the absence of connectivity encoding by place cells is not due to a poor representation of space, and suggests that rats knew in which compartment and which position in that compartment they were, coherent with our findings that they could discriminate locked from open doors even though all doors looked identical.

### Place cells did not spatially remap between task phases

Do place cells remap depending on the task at hand? This has been shown to be the case under some conditions, usually if tasks are performed in separate blocks (Markus et al. 1995; Wiener, Paul, and Eichenbaum 1989; Spiers, Olafsdottir, and Lever 2018), but not when tasks are intermingled within the same experimental session (Trullier et al. 1999; Duvelle et al. 2019). In the present experiment, assessing whether this is the case would indicate if our results observed mostly during foraging do, or do not, extend to the goal-directed parts of the task. We automatically categorised behaviour into ‘foraging’ or ‘goal-directed’ epochs (*Methods - Behaviour discrimination*), see Fig. 11A for an example. As expected, the running speed was higher for goal-directed epochs than foraging (Fig. 11B, Table 7). Incidentally, we found an effect of session on speed with slower speeds in later sessions, probably indicative of decreasing motivation as time passes (Fig. 11B).

**Figure 11:**
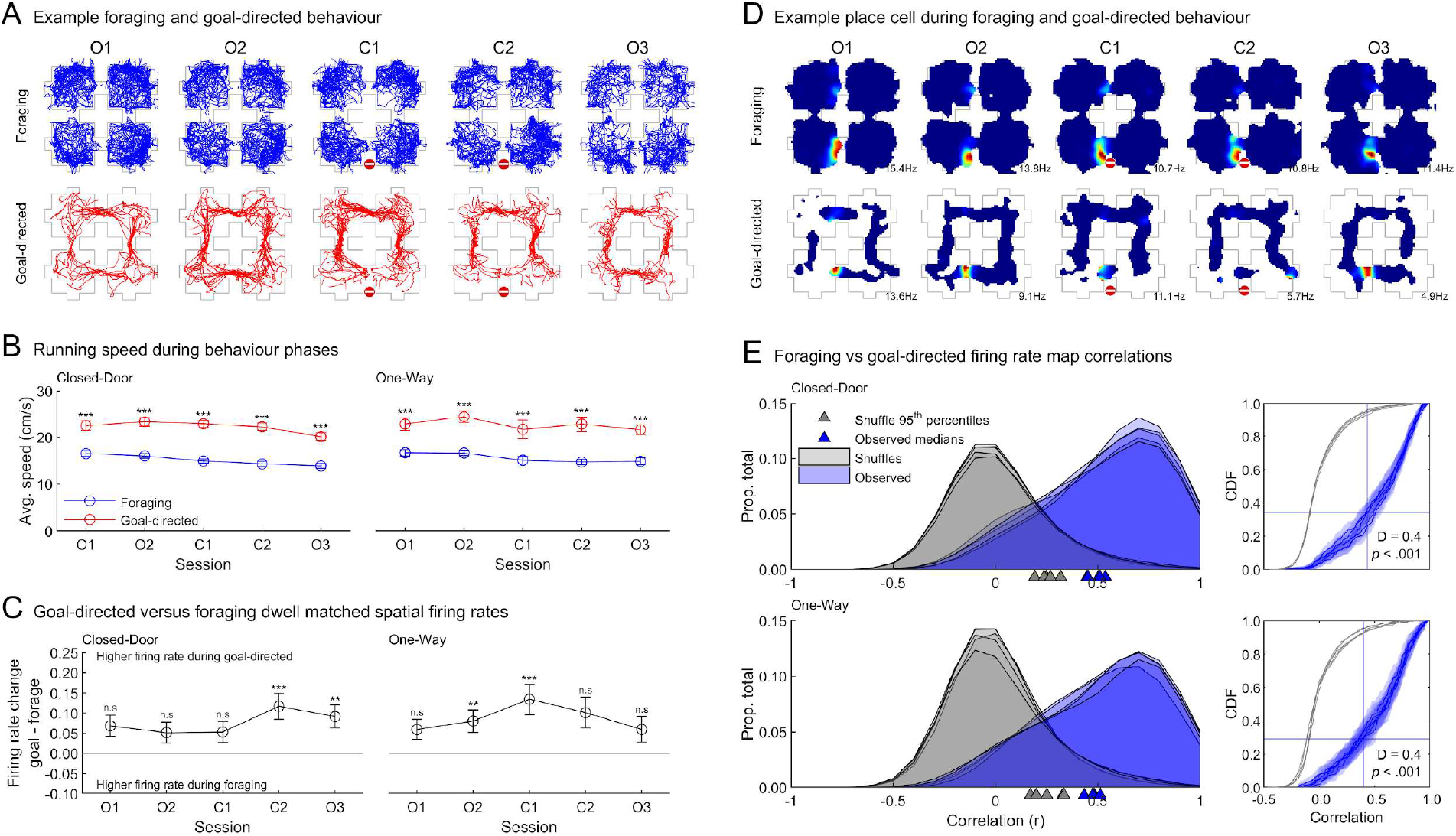
No spatial remapping between foraging and goal-directed epochs. See Table 7 for all statistical results. **(A)** Example of trajectory discrimination between foraging (top) and goal-directed (bottom) for one Closed-Door sequence. Note that foraging trajectories are comparable between sessions while goal-directed ones are strongly affected by the door closure; see Sup. Fig.4 for dwell maps averaged over all sessions. **(B)** Average running speed over sessions for the different task phases. Speed during goal-directed movement was always higher. **(C)** Comparison of firing rate between task epochs, computed over common spatial bins as the difference of goal-directed and foraging average firing rates, averaged over place cells. Positive numbers indicate increased firing rate during goal-directed epochs. **(D)** Example rate maps of a cell recorded in the same session as A in a Closed-Door sequence, divided between foraging (top) and goal-directed (bottom). Note that the spatial activity of this cell remains stable between the two types of behaviours. Colour scale is the same as previous rate maps, number indicates peak firing rate. **(E)** *left*, distributions of the correlations between goal-directed and foraging rate maps for all sessions (O1, O2, C1, C2, O3) in blue and shuffled data in grey. *Right*, cumulative distribution function of each group. The 95^th^ percentile of the shuffle distribution (O2) is indicated by a vertical blue line and the intersection of this with the O2 data distribution is indicated by a horizontal line, showing that more than 60% of the cells are more stable than chance. Text gives the result of a two-sample Kolmogorov-Smirnov test comparing the two distributions in session O2.

**Table 7:** Statistical test results corresponding to Fig. 11.

| Figure and dependent variable | Omnibus test | Effect of group (closed/control or open) | Effect of session | Interaction group × session | Post-hoc tests across groups (corrected*) |
| --- | --- | --- | --- | --- | --- |
| Fig. 6B left<br>Foraging vs goal-directed running speed Closed-Door | two-way repeated measures ANOVA | $F(1,72) = 156.2$,<br>$p = 2.618067\text{e-}10$ | $F(4,72) = 6.8$,<br>$p = 1.027453\text{e-}04$ | $F(4,72) = 4.2$,<br>$p = 4.298650\text{e-}03$ | All $p < .001$ |
| Fig. 6B right<br>Foraging vs goal-directed running speed One-Way | two-way repeated measures ANOVA | $F(1,40) = 47.5$,<br>$p = 4.224901\text{e-}05$ | $F(4,40) = 3.9$,<br>$p = 8.931908\text{e-}03$ | $F(4,40) = 1.2$,<br>$p = 3.190040\text{e-}01$ | All $p < .001$ |
| Fig. 6C left<br>Foraging vs goal-directed firing rates One-Way | Independent one-sample t-tests with Holm-Bonferroni correction | Compared to chance (0):<br>From left to right: $p > .99$, $p > .99$, $p > .99$, $p < .001$, $p = .002$ | | | |
| Fig. 6C right<br>Foraging vs goal-directed firing rates One-Way | | Compared to chance (0):<br>From left to right: $p > .99$, $p = .004$, $p < .001$, $p > .99$, $p > .99$ | | | |

\*Note: all post-hoc tests were based on estimated marginal means and corrected using the Tukey-Kramer method.

We next examined the difference in the average firing rate of place cells between foraging and goal directed behaviour epochs, comparing only locations occupied in both conditions (Fig. 11C). The difference of firing rates was always positive, indicating that cells fired more during goal-directed periods, although significantly so only for a subset of sessions (two-sided signed rank tests with Holm-Bonferroni correction). This is coherent with previous reports that place cell firing is modulated by speed (McNaughton, Barnes, and O’Keefe 1983).

We then computed separate rate maps for each epoch (*Methods - Remapping between foraging and Goal-directed behaviour*); examples can be seen in Fig. 11D and the distributions of correlations between epochs for each session type are shown in Fig. 11E. In all sessions, the median of the observed correlations exceeded the 95^th^ percentile of a corresponding chance distribution, demonstrating that the data from both task phases were more similar than chance and evidencing an absence of global remapping between foraging and goal-directed phases in this paradigm (Fig. 11E).

To conclude, we found that: *i*) place cells do not encode connecting points such as doors differently from other locations; *ii*) their firing location and firing rates are unaffected, whether locally or globally, by changing the connectivity of the environment; *iii*) place cells’ activity patterns are consistent with a mixed global and local representation where the global representation of position dominates despite the repetitive geometry of the environment; *iv*) these cells do not spatially remap between the foraging and the goal-directed alternating phases of the task.

## Discussion

We tested whether hippocampal CA1 place cells encode changes in spatial connectivity in their firing rate patterns in an interconnected 4-room environment. We found that rats rapidly learned the changes in connectivity as shown by their preference in using open doors over locked doors. Based on past experimental results and current models of hippocampal place cell dynamics, we predicted that place cells would over-represent doorways and show changes in their firing rates or field locations when doors were changed from open to locked. We found no evidence of such coding, and this was not due to poor spatial localisation by the rats or their place cells as these encoded each compartment differently. Thus, after controlling for possible internal and external confounding factors, our findings suggest that dorsal CA1 place cells do not explicitly encode connectivity or its changes, raising the question of which other brain region or mechanism might encode this information. Additionally, we observed a dominant global code across the population, despite the repetitive geometry of the environment. Still, there were clear examples of cells expressing local coding. Such a mixed population code provides the potential for greater flexibility in navigation: from the global code, an animal can learn that specific, unique locations are to be approached or avoided while from the local, repeating, code, an animal can generalise across the environment to avoid or approach some combination of features, e.g. doorways to the north. These findings and their implications for current models of place cells are discussed below.

### Navigation in connected spaces

Rats and humans can flexibly adapt to noticeable changes in the connectivity of an environment such as inserted barriers or canyon-like drops (Tolman 1948; Alvernhe, Save, and Poucet 2011; Poucet and Herrmann 2001; Alvernhe et al. 2008; Alvernhe, Sargolini, and Poucet 2012; Grieves and Dudchenko 2013; Javadi, Patai, Marin-Garcia, Margolis, et al. 2019; Javadi, Patai, Marin-Garcia, Margois, et al. 2019; Simon and Daw 2011; de Cothi et al. 2020). Extending these results, we found that rats were able to learn and remember the status of doors being locked or unlocked, in the absence of directly perceivable changes other than the direct feedback when pushing on a door. To our knowledge, only one other similar result has been reported (Okaichi 1996) where rats used their knowledge of available unlocked doors to select the optimal path to a visible goal through a maze, see also (Grieves and Dudchenko 2013)). In addition to this, we show that rats are able to separately learn the connectivity status of the two sides of a door. Importantly, the bias towards open doors persisted into the last session, after a pause outside of the maze, showing that the effect of locking doors on behaviour was not due to working memory or to obvious sensory cues. This is similar to previous reports (Alvernhe, Save, and Poucet 2011) where the behavioural and neural effects of introducing a transparent barrier were still visible once the barrier was removed.

Did rats rely on a global representation of their environment and a map-based strategy to reach the goal box (Arleo and Rondi-Reig 2007; O’Keefe and Nadel 1978)? We observed direct, detour-like trajectories when a door was locked (ex, Fig. 2E), showing that at least in those cases rats appear to rely on a global map of the environment. Rats tended to choose the door corresponding to the shortest path to the goal, suggesting knowledge of the goal location and of the maze layout, but this was not always significantly different from chance. Rats were only significantly better than chance at choosing the correct box to forage when it was immediately connected to the start, which may be explained by the low cost of quickly checking for food on the way to the goal. In summary, our results are consistent with rats using a map-based strategy to navigate this maze, but future experiments would help confirm this. For example, employing a targeted training of the sound-goal association would ensure localisation of the goal, and use of different experimental conditions (light, dark, cue rotations) would help determine which strategy and cues the rats used to navigate.

### Place cells do not explicitly encode spatial connectivity

Environmental connectivity has long been thought to be an integral part of the cognitive map (Tolman 1948), a view later reinforced by behavioural experiments and theories of cognitive maps; for reviews, see (Poucet 1993; Epstein et al. 2017). We anticipated that place cells would encode connectivity explicitly (in the firing of individual cells), especially locally to a change, but this was not the case. A very small proportion of place cells remapped when one or several doors became locked, but this was less than the number of cells remapping between two sessions of unchanged connectivity, suggesting that this was not specific to connectivity updating but perhaps underlying latent time coding as has been observed previously (Mankin et al. 2015; 2012). Overall, we did not find any observable effects of changing the connectivity of the environment on CA1 place cell firing, investigated in terms of rate maps correlations, average firing rate, place field locations, place field size, firing rate or extent through doors, either for all activity or only locally to the changed doors. We also did not encounter any ‘door’ cells (active at all 4 doors), nor did we find doors to be overrepresented, unlike past studies in multicompartment environments in which open doorways connected compartments (Spiers et al. 2013; Grieves et al. 2017).

Thus, after controlling for confounding factors, such as occupancy, speed and movement direction (by only analysing comparable behaviour between conditions), time (by comparing pairs of sessions separated by a similar time difference) and geometry (by locking existing doors instead of introducing a new barrier), we found no evidence of explicit encoding of connectivity by CA1 place cells. In the study of Alvernhe et al. (2011) where remapping was found after introducing a barrier to create a detour, this experimental manipulation also changed the rats’ behaviour as well as the geometry of the environment, both of which could have impacted the firing of place cells. Indeed, boundary vector cells (BVCs) in the subiculum, thought to contribute to place cell firing (Barry et al. 2006; Hartley et al. 2000), have informally been observed to fire in response to plexiglass walls (personal communication by Colin Lever, July 2019). Place cells themselves can change their firing in the vicinity of a transparent barrier (Rivard et al. 2004). However, in a shortcutting situation (Alvernhe et al. 2008), remapping near the geometric change was found in CA1 and CA3 place cells, as well as far from the change for CA3 cells only, indicating that opening a connecting node and closing one might be encoded differently or that environmental connectivity could be encoded in CA3. Other nearby regions could be encoding spatial connectivity instead of the hippocampus, such as the subiculum and its high concentration of BVCs and border cells (Brotons-Mas et al. 2017; Lever et al. 2009), or grid cells in the medial entorhinal cortex, the activity of which is affected by geometric changes in the environment (Krupic et al. 2016; Wernle et al. 2018). Alternatively, a more remote region could encode environmental connectivity, such as the prefrontal cortex, in which activity levels vary depending on the available connections from possible future paths in humans (Javadi et al. 2017; Spiers and Gilbert 2015). Finally, as previously mentioned, connectivity could be encoded implicitly, either i) in the co-firing of place cells (Dabaghian, Brandt, and Frank 2014), a view not strongly supported by our findings of unchanged place field extent through doors when they became locked, or ii) in the non-local, sequential ‘replay’ of place cells (Wu and Foster 2014; Pezzulo, Kemere, and van der Meer 2017). Future experiments with many simultaneously recorded cells (Pfeiffer and Foster 2013) could examine the effects of connectivity modulations on non-local firing or co-firing in the absence of confounding factors.

### Implications for models of place cell dynamics

Our results may help the development of the current computational models of hippocampal place cells. The ‘Successor Representation’ (SR) model proposes that place cells instantiate a predictive map, representing each state, or position in an environment, in terms of its successor states, or possible future positions available from that place (Stachenfeld, Botvinick, and Gershman 2017; de Cothi and Barry 2019). This model predicts changes in the firing of place cells when transitions become blocked in an environment (see Figures 2, 3 and S3 in (Stachenfeld, Botvinick, and Gershman 2017)). However, we found no evidence of such changes in our task. One interpretation of the SR model is that it might predict changes in the representation of the environment between the goal-directed periods of our task and the foraging periods, due to the change in policy. Akin to other studies where two tasks or behaviours are alternated in the same environment (Trullier et al. 1999; Duvelle et al. 2019), we found no change in spatial firing between goal-directed and foraging phases. One possibility is that the patterns of place cell dynamics best explained by the SR model are observed in tasks requiring stereotyped, repetitive behaviours linked to more consistent policies.

We found that doorways are not overrepresented and that the majority of place cells provide a unique code for each room, encoding position in a global reference frame. These findings appear to contradict the BVC model of place cell firing, which predicts both fully repeating place fields and overrepresentation of doorways in previous paradigms with repeating environments (Barry et al. 2006; Grieves, Duvelle, and Dudchenko 2018). It is also intriguing that we did not see remapping following door-locking as this should technically change the status of a door from passable to being a boundary. However, predictions of this model in our environment rely on knowing how biological BVCs would respond to doors, which - unlike corridors - obstruct the view and slow down passage even when not locked, which is currently unknown. If doors were represented as walls, our results could be explained by a modified version of the BVC model: the contextual gating model (Jeffery et al. 2004; Hayman and Jeffery 2008). By this view, BVC inputs (which are purely spatial) are “gated” by non-spatial cues such as colour or door of the environment, task demands, combinations of distal cues or local cues, such that different BVCs drive the cells in different contexts. In the present case, if different boxes were considered different contexts, this model would predict remapping between boxes. The small number of neurons showing full place field repetition (in all 4 boxes) or those showing repetition in only a subset of boxes could be explained as cells whose geometric inputs were not always gated by any of the available contextual cues. However, it is not clear why each box would be considered a different context in our case but not in previous cases showing place field repetition (Spiers et al. 2013; Grieves et al. 2016; Harland et al. 2017).

Additional factors that might explain differentiation of compartments are the use of a navigation task, the greater salience of distal landmarks or the presence of uncontrolled local cues such as self-deposited odours or rats being able to visually differentiate real and dummy doors. However, several experiments with strong polarising distal cues have also found evidence of place field repetition, indicating that distal cues may not minimize repetitive place fields (Singer et al. 2010; Grieves et al. 2016; Derdikman et al. 2009; see also supplementary data in Grieves 2015). It is also unclear that either the addition of a spatial task or use of self-deposited odours could explain our results as rats were impaired in a spatial discrimination task taking place in a parallel version of a 4-room maze, in which place field repetition was also observed; additionally, if self-deposited odours were sufficient to help discriminate compartments, place field repetition would not have been observed (Grieves et al, 2016). Another explanation for global coding of position is that the different orientation of the two entry points into each box may have helped disambiguation or pattern separation of the boxes (J. K. Leutgeb et al. 2007). Indeed, the point of entry into a single compartment has been shown to affect place cell firing (Keinath et al. 2020; Sharp, Kubie, and Muller 1990) and a key feature of past experiments evidencing place field repetition is that each compartment was entered in the same environmental direction.

## Conclusion

We recorded dorsal CA1 place cells in the hippocampus of rats alternating pseudo-random foraging and goal-directed trajectories in an environment composed of four connected boxes. When the connectivity of the environment was modified by locking doors, rats rapidly adapted their navigation behaviour, showing knowledge of the updated connectivity, but place cells did not specifically modulate their firing activity, either in terms of firing location or firing rate, even in the vicinity of the affected doors. Place cells did not spatially remap between the foraging and goal-directed phases of the task, but their firing became generally decorrelated with time. The large majority of place cells encoded position in a global reference frame, i.e. with a unique combination of active fields in each box; this was intermixed with a subset of place cells showing local coding of the environment. Our results help clarify the extent to which CA1 contributes to representing connected environments and their structure, which will aid the refinement of models of hippocampal mechanisms and function.

## Material & Methods

### Overall summary

Upon arrival in the rat colony, rats were handled daily for at least a week, implanted and allowed to recover for another week. They were then food-restricted, trained, and screened for place cells. Once rats had passed all training phases and signals from putative place cells (> 4 simultaneous cells) were detected, they were recorded in two different phases of the ‘4-room navigation task’ (1 phase per day), in baseline conditions (all doors open) and ‘Closed-Door’ or ‘One-Way’ conditions (1 door locked both ways or all doors locked only one-way). After successful recordings in each phase of the task, the tetrodes were lowered in an attempt to detect new cells and recordings were repeated in each phase, usually with a new door condition (e.g. a different door was locked or all doors were locked the other way); this process continued until fewer than 4 cells were detected simultaneously. Rats were then perfused transcardially with saline followed by 9% formalin and their brains were extracted and stained to confirm the location of recording tetrodes. These methods are detailed below.

### Subjects

Five Lister-Hooded male rats weighing approximately 300-600g and aged 3-6 months at the start of the experiment were used. They were first housed in pairs at 20 ± 2°C under a 12/12h light/dark cycle starting at 12 AM. They were provided with ad libitum water, food, environmental enrichment and daily handling from the experimenter. After implant surgery, rats were housed individually and allowed to recover for one week before food-restriction started to maintain their weight at 90-95% of free-feeding body weight. One of the rats had prior experience with a spatial task in the same experimental room and another one had prior experience with a linear track task in a different room; the other 3 rats were naive. All procedures complied with the national [Animals(Scientific Procedures) Act, 1986 United Kingdom] and international [European Communities Council Directive of 24 November 1986 (86/609/EEC)] legislation governing the maintenance of laboratory animals and their use in scientific experiments.

### Room and experimental environment

The experimental room was equipped with distinct distal visual cues on each wall and dim ceiling lighting. A schematic of the apparatus and room cues as well as photos is shown in Sup. Fig.1. The experimenter, the recording system and the computer desk were in the same room. The custom-built recording apparatus consisted of four 60×60 cm grey-painted wooden boxes connected to each other via four 16×16 cm dark grey door systems. Each box also had two ‘dummy doors’ (same dimensions and colour as the real doors, but made of one panel instead of two) appended to their external sides (Fig. 1). All walls were 20 cm high and were made of painted hardboard with small (∼4mm diameter) perforations. On the corner of each box, a bell was placed which could be activated remotely by pulling a string from the experimenter’s desk. The bell system acted as a sound cue to inform the rat of the next rewarded box. The door mechanism consisted of two vertical panels each glued to a rotating wooden rod and equipped with one spring each, to ensure that the door would remain shut unless it was being pushed. Curved black plastic stripes were added on the top of the doors to guide the recording cables. A slot present at the top of each door panel allowed for the insertion of 4 small metallic locks that could block the panel on one or both sides. Since the locks were inserted at the top, the rats could not see them unless they were in a rearing position. Thus, the doors would look and feel the same from a rat’s point of view whether locked or not, the only difference being that locked doors could not be pushed open. Note that we indifferently use “locked” or “closed” to mean a door that cannot be pushed. The 4-room environment was placed on top of 60 cm-high cardboard boxes. A padded headcap was added around the drives and headstages to help rats with door-pushing and absorb possible shocks on the implants. An overhead camera centred on the environment provided video input to the tracking system. Tracking and electrophysiology data were collected using a 64-channel recording system (DacqUSB, Axona, St. Albans, UK). Video recordings were sometimes made using the DacqTrack system (Axona, St. Albans, UK). During screening, recording and most of the training, the animal was connected to the recording system via 4 flexible cables attached to the ceiling by elastic bands.

An elevated rotating platform (80 cm high) was placed next to the environment where rats could rest before and after screening / training / recording sessions. For screening sessions (i.e. monitoring brain signals to decide whether tetrodes were in the hippocampal cell layer or not and move the tetrodes accordingly), a plastic 120x×120 cm black square with 20 cm high walls was placed on top of the 4-room where rats could freely forage.

### Task

The final task, used during all recordings and most of the training, consisted of separate sessions which each contained an exploration phase followed by several ‘trials’ starting at a bell sound and ending before the next bell sound. The food reward used during the experiment was either rice krispies / coco pops (Kelloggs, Michigan, US) or rice pops / choco pops (Waitrose and partners, London, UK). The reward type could change across days but would remain the same throughout a given recording sequence.

During the **exploration phase**, the rat was placed in a given compartment with no food provided and allowed to explore all 4 boxes. After this, the **task trials** started. On a given trial, the bell of a specific box was rung and food was thrown there by the experimenter; once the rat reached the rewarded box, more food would be thrown or placed in the box at regular intervals. During this specific step, the experimenter would sometimes come closer to the box to place food in specific places; attention was drawn to placing food in front of doors (real or dummy), in corners, and in any under-sampled locations. The ‘trial’ ended after a given time passed in the same box (at least 30s, excluding time spent out of the rewarded box). Then, a new trial started using the next box, selected from a predetermined goal list, until all boxes of the sequence had been visited. The **goal list** was created pseudo-randomly by the experimenter with the following constraints: each box (A, B, C and D, Fig. 1 B) should be used 3 times as goal, making up 12 trials in total; all four boxes should be used before repeating a box; distances between successive boxes should vary. An example goal list would be ABDC BACD CBAD. A new list was generated for each recording sequence.

A daily recording **sequence** was divided into 5 sessions, each using the same goal list. The first two sessions were considered the **baseline**, with all four doors open both ways. The next two sessions were the **test sessions**, using either a ‘Closed-Door’ configuration (1 door locked both ways) or a ‘One-Way’ configuration (all 4 doors open only one way). The last session of the sequence was back to baseline, i.e. all doors open. In between sessions, the rat was placed on the elevated platform with access to water and allowed to rest for 5 or 10 min. During this time, the experimenter manipulated all 4 doors to minimise possible olfactory biases and locked or opened the appropriate doors.

In summary, there were two possible sequences:

- **‘Closed-Door’ sequence** = all open, all open, closed, closed, all open (Fig. 1B).
- **‘One-way’ sequence** = all open, all open, one-way, one-way, all open (Fig. 1C).

A ‘Closed-Door’ sequence was generally followed on the next day by a ‘One-Way’ sequence or was repeated with a different locked door. Tetrodes were advanced by at least ∼25µm to sample new cells only when a rat had successfully completed one of each sequence type. In some cases the same sequence type was recorded without advancing the tetrodes, meaning that the same sample of place cells could have been recorded, but the order of rewarded boxes and the chosen locked door(s) were always different in each session. The maze was cleaned with alcohol spray between animals, and urine traces or faeces were locally cleaned on the rare occasions when they appeared during the experiment. No attempt was made to remove other possible olfactory cues left by the rats between sessions of a recording sequence.

### Training

All rats were pre-trained post-surgery in the four-room environment for 2 to 5 sessions every weekday, as detailed below. Two of the rats were also used before implant surgery to pilot the door system, one of those in a different room and apparatus. Training in the 4-room took 6 to 15 days (median = 10 days) which depended on how fast the rats reached the final criteria, during which they were also screened for hippocampal cells. Training aimed to familiarise rats with: i) pushing doors in order to transition from box to box, ii) foraging efficiently in each box and iii) running for food for long periods of time (the experiment lasting on average 2.5h each day). No explicit training of the association between the goal and the bell sound was done. Undesirable behaviours like climbing on walls or doors and chewing maze parts were discouraged from the start of training by either making a loud clapping noise or pushing the rats. Training data were not analysed.

#### 1. Familiarisation

Rats were allowed to freely explore the maze without food. After 5 min, rats were taken out of the environment and placed onto the elevated platform. This was repeated once per box.

#### 2. Door training

With food present in all 4 boxes, rats were released in a random box and taken out of the environment after 10 min or once all food had been eaten, whichever came first. After 5 min, if the rat hadn’t gone spontaneously through a door, the experimenter would encourage door-pushing (e.g. going through a door with a hand, holding the door half-open, attracting the rat to the other side of a door with noises or food). This phase was repeated until rats were able to easily use doors to move from box to box. Interestingly, most rats started to use the doors spontaneously from early maze exposure.

#### 3. Task learning

Rats were plugged in to the recording system from the third session of this phase. Here, rats were doing the task as described above (*Methods – Task*), except that each training session lasted 20 min maximum. For three of the rats (r35, r37 and r38), locked doors (either one closed door or one one-way door) were introduced once they had mastered the baseline condition of the task. For the other two rats, closed-door sessions were added every 2-4 ‘all-open’ sessions. For all rats except r35, only two of the doors were used as closed or one-way doors during training so that the other two, new doors could be closed in priority during the actual recording. Rats were considered ready for recordings once they could do 11 trials (as defined in *Methods – Task*) in 20 min for two out of the last three training sessions, except for r44 for whom the criterion was 11 trials in 25 min. Rats’ behaviour was generally quite disturbed upon their first closed-door encounter (i.e. rats might try to pull on doors, push harder, climb, or avoid all doors altogether) but stabilised after a few more encounters.

### Microdrive and implantation surgery

Axona drives (MDR-xx, Axona, UK) were loaded with 8 tetrodes made of four 17 μm of diameter platinum-iridium wires (California Fine Wire, Grover Beach, CA). Tetrode tips were gold-plated (Non-Cyanide Gold Plating Solution, Neuralynx Inc., MT) using a NanoZ system (White Matter LLC) to reduce the impedance to 180–250 kΩ at 1kHz. 2 drives were implanted on each rat (one per hemisphere), above the CA1 field of dorsal hippocampus, using standard stereotaxic procedures under isoflurane anaesthesia and sterile conditions. The coordinates relative to Bregma were as follows: AP: -3.5 to 4.0 mm; ML: ±2.4mm; DV: 1.3 to 1.5 mm from dura surface. Both drives shared the same ground wire, connected to a ground screw above the cerebellum. 6 jewellers’ screws helped anchor the drive to the skull, together with one layer of Super-Bond C&B (Sun Medical, Shiga) followed by several layers of dental cement (Simplex Rapid, Kemdent®). A long-acting analgesic (Carprofen) and saline solution were given subcutaneously at the start of surgery. Post-surgery, rats were provided with another analgesic (Meloxicam) in their food for 3 days.

### Screening and recording

Daily or twice-daily screening sessions (spaced by at least 4h) started 1 week post-surgery, during which the animal rested on the elevated platform then foraged for the same reward used in the experiment in the square plastic environment for 8-16 min. Signals were screened for signs of sharp-wave ripples and pyramidal cells. If no hippocampal activity was detected, tetrodes were lowered by approximately 25 or 50μm. Extracellular activity was collected with DacqUSB, the signal was first sampled at 50kHZ, amplified then band-pass filtered between 300 and 7000 Hz, digitized at 48 kHz and could be further amplified 10–40 times at the experimenter’s discretion. Local Field Potential (LFP) data was obtained by sampling signals from selected channels at 4.8 kHz. Note that LFP data was not used in the current study.

### Histology

When no more than 4 putative CA1 signals were observed in a given rat and sharp-wave ripple amplitude was observed to decrease on both drives, small electrolytic lesions were created by passing a positive current (5 µm for 10s) through chosen electrodes while the animal was deeply anaesthetised. The rats were then overdosed with pentobarbitone before being transcardially perfused with a saline solution followed by a formalin solution (10%). The brains were preserved in formalin for at least 2 days, then, optionally into a 30% sucrose solution for another 2-3 days. Brains were then frozen and sectioned in 40 microns slices stained with Cresyl Violet. The electrode tracks and lesions signs were detected under a microscope to confirm recording sites. All recording sites used in the analysis were confirmed to be in hippocampus CA1 on both hemispheres (see Fig. 12).

## Behavioural analyses

### Position tracking

The head position of animals was tracked continuously at 50Hz, using an infrared LED affixed to the recording headstage and custom tracking software (DacqTrack, Axona Ltd., St. Albans, UK). For segments of missing tracking data, we simultaneously interpolated and smoothed the existing data using an unsupervised, robust, discretized, n-dimensional spline smoothing algorithm (Matlab function *smoothn*, (Garcia 2010; 2011)). For each sequence of sessions, we manually fitted a wire-frame to the position data, from which we extracted the maze boundary and doorway positions.

### Event flags

Experimenters recorded behavioural events such as door pushes and bell sounds online during recording by pressing on a miniature wireless keyboard. The recording system stored the time and type of keypress synchronised with neural and position data. Although the majority of these manual event flags were correct and utilised in priority, incorrect flags were corrected programmatically offline based on the animal’s tracked position in combination with trial-specific data. These corrections apply only to the behavioural measures related to bell sounds (*Methods - Response to bell sounds*), correct door pushes and correct foraging (*Methods - Correct door pushes and correct foraging*). Bell sound events recorded by the experimenters were never corrected. However, a minority of trials were rejected because the rat was tracked in the rewarded compartment synchronously with the bell sound and thus no first door push or foraging choice could be determined. In further trials a door push was recorded that was not possible given the animal’s location and these were replaced with the first door through which the animal moved after the bell sound. The majority of these two error types were due to short time lags between the animal’s behaviour and registering the event flag key press. Lastly, in a subset of trials, no door push was recorded and the missing value was filled using the first door through which the animal moved after the bell sound.

### Behaviour discrimination

Animals engaged in two different modes of behaviour which we termed ‘foraging’ or ‘goal-directed’. When animals moved through the maze in a fast and direct way (for instance after a bell sound but before they reached the food), marked by rapid and direct locomotion, we classified the behaviour as goal-directed. Other periods were spent freely searching for food (for instance after arriving in a rewarded box) and were marked by slower and circuitous locomotion, this was categorised as foraging. To differentiate these behavioural modes we looked at each visit an animal made to a box. The visit was categorised as foraging if:

1. the distance covered in the visit was more than 120cm and;
2. the animal covered more than 20% of the box area during the visit.

Distance was calculated as the total distance travelled along the visit path in non-overlapping 1s windows, coverage area was calculated by dividing boxes into 100 bins (unsmoothed) and counting the proportion of bins containing more than 1 position data sample. For comparison, the minimum distance between two doorways along a circular arc would be approximately 50cm and cover approximately 10% of a box. Conversely, rats were considered to be in a goal-directed mode if:

1. moving through a door (1s before to 1s after) or;
2. pushing on a door (2s before to 1s after) or;
3. not foraging.

### Response to bell sounds

To test whether rats were aware of and responded to the bell sounds, we first compared their running speed in the period 1s before and 1s after each bell sound. Instantaneous speed was calculated for every position data point as the sum of distances travelled between the preceding position data point and the following one divided by the time between them.

Next, we looked at the time between a bell sound and the next door push and compared these values to a shuffle. For the shuffle we generated *N* random time points uniformly throughout each session where *N* was the number of real door push events. We then calculated for each bell sound the time between that and the next random time point. This procedure provides the time expected between bell sounds and door pushes if the two events were completely dissociated.

### Correct door pushes and correct foraging

We next sought to determine whether rats navigated to rewarded boxes using optimal paths. For each session we generated a graph with directed edges (*Matlab digraph*) representing the nodes and possible routes given the maze structure and connectivity. Using this graph, we calculated the minimum number of doors that would need to be crossed when moving between any two boxes and the optimal door sequence that would need to be used. For example, with all doors open the minimum distance between boxes A and B would be 1 - moving through the door directly between them. However, if this door was locked the minimum distance would instead be 3 as the rat would have to travel via boxes D and C.

With this approach, for every bell sound we looked at the first door push the rats made following the bell and assessed whether this door belonged to the optimal path to the rewarded box given the maze connectivity. We then analysed the performance of rats depending on the distance from the start box to the goal box. In this analysis, when all doors were open (sessions O1, O2 and O3) we discarded trials where the rat was diagonally opposite the correct box (distance of 2) as in this case both doors and paths would be equally optimal. Similarly, in the open sessions the rat could never be 3 or more doors away from the correct box. In the sessions with a connectivity change (C1 and C2), optimality at all possible distances was assessed.

Regardless of the route taken after a bell sound we also sought to determine whether rats preferentially started to forage in the rewarded box. For this we looked for the first box in which the rat foraged after each bell sound (Methods: *Behaviour discrimination*) and assessed whether this was the rewarded box or not. This analysis was independent of the optimality analysis described above, meaning that a rat could take a non-optimal path or push on locked doors but still forage first in the rewarded box.

### Neural activity analyses

Single-unit data were first processed using an automated spike-sorting algorithm (Klustakwik v3.0,(Kadir, Goodman, and Harris 2014)) using the first three principal components and peak waveform amplitude as parameters. Manual refinement of the classification was then done using the TINT spike-sorting software (Axona, St Albans, UK). Only well-isolated putative pyramidal cells were kept (pyramidal waveforms, no or few spikes in the refractory period).

### Firing rate maps and spatial information

Spike and dwell time maps were constructed as bivariate histograms of the spike and position data respectively (2cm square bins, Matlab: *hist3*) after speed-filtering the data to remove periods where the animal’s running speed was less than 5 cm/s (to avoid contamination of the data by possible reactivation events). These maps were then smoothed with a two-dimensional Gaussian kernel (standard deviation: 2.5 bins, kernel size: 9×9 bins, Matlab: *imgaussfilt*). Firing rate maps were calculated by dividing spike maps by the corresponding dwell map for that session. In all maps, bins visited by the animal for less than 0.05s were considered empty. This procedure was repeated using i) all spike and position data, ii) data filtered to include only foraging behaviour and iii) data filtered to include only goal-directed behaviour, always using only speed-filtered data (Methods: *Behaviour discrimination*).

Spatial information content in bits/second was calculated using a method reported previously (W. E. Skaggs et al. 1996) as:

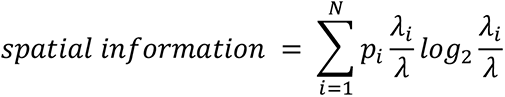

where the environment is divided into non-overlapping spatial bins *i* = 1,…, N, *p_i_* is the occupancy probability of bin *i*, *λ_i_* is the mean firing rate for bin i, and λ is the overall mean firing rate of the cell. We compared these spatial information values to a shuffle distribution using a method reported previously (Jercog et al. 2019); briefly, spike sequences were time-shifted by a random time interval (between 20s and the duration of the session minus 20s) in a circular manner. This method maintains their temporal order and total number whilst decoupling the spike times from the animal trajectory. For each shuffle we reconstructed a firing rate map and recalculated its spatial information content as described above. This procedure was repeated 100 times for each neuron. A neuron was defined as significantly spatially modulated if the observed spatial information content exceeded the 95^th^ percentile (Matlab: *prctile*) of its shuffled distribution.

Whenever activity between compartments was compared (e.g. place field repetition analysis, doorway rate map correlations analyses), the rate maps for individual boxes were smoothed separately to avoid spuriously smoothing data between neighbouring boxes. We constructed box-specific maps by cutting position and spike data to within the boundary (minimum enclosing rectangle) of each box. Firing rate maps were then constructed as described above. From these box-specific maps we also extracted 25cm radius regions around each doorway which were used in the doorway specific analyses. In all other cases, i.e. when whole maps of the environment were compared to each other, the entire map was smoothed as a whole.

### Unit classification

Units were classified as putative place cells if their mean firing rate was greater than 0.1Hz in at least 2 sessions and if, in the session with the highest firing rate:

1. Average firing rate was greater than 0.1 Hz and less than 5 Hz and;
2. Spatial information content was greater than 0.5 bits/second and;
3. Spatial information content exceeded the 95^th^ percentile of a spike train shuffle and;
4. The cell’s width of waveform (peak to trough) was greater than 300µs.

A cluster’s width of waveform was defined as the time between its maximum and minimum amplitude (peak to trough). We extracted this value from the channel with the greatest amplitude in the session with the highest firing rate. All further analyses were conducted only on those cells passing these criteria, which aimed at removing possible interneurons, silent cells and non-spatial cells. Cells were analysed separately in the two sequences of conditions (which may contain repeated recordings of the same cell). Within the same condition, it is possible that the same cell was recorded more than once, although we would often lower the tetrodes by at least 25µm between two recordings of the same type and we manipulated a different door in consecutive recordings of the same type. Finally, all analyses were repeated on one session per rat – the one with most simultaneously recorded cells - with similar results and conclusions (data not shown).

### Rate map correlations

We correlated the spatial activity of place cells at three different levels of specificity, between sessions, between boxes and between doorways. All correlations were pairwise Pearson correlations (Matlab *corr*) and we only correlated a pair of firing rate maps when at least one map had a peak firing rate greater than 1Hz (Grieves et al. 2016). Using this process we compared every session to every other session within a sequence, every box to every other box within a sequence and every doorway to every other doorway within a sequence.

For session comparisons we extracted the correlations between sessions O1 and O2 (first two sessions with all doors open) which gives a baseline measure of the stability of place cells in an unchanging environment. We then extracted the comparisons between O2 and C1 (the last open door session and the first closed door session); if cells changed their firing in response to a change in connectivity we would expect these correlations to be lower than the baseline. We also looked at the change in firing over time by extracting correlations between sessions separated by increasing durations (i.e. O1 and O2 are consecutive while O1 and O3 are separated by 3 sessions or approximately 90 minutes).

We also looked more specifically at activity around the doorways. For this we looked at each side of every door separately and we categorised these into two groups: the ‘closed/control’ group consisted of the doors that were locked during sessions C1 and C2, the ‘open’ group consisted of all other doors. Note that in the One-Way sequence a single doorway will contribute one side to the closed/control group and one side to the open group. Next, we extracted the correlations between sessions O1 and O2 separated into closed/control and open doorway values. These were then averaged so that every cell contributed one value to each group. As before, this acts as a baseline and we compared it to the same values extracted from correlations between sessions O2 and C1.

### Individual remapping between sessions

To more broadly categorise remapping between the different sessions, for every individual cell we looked at if, where and when remapping occurred between consecutive sessions. First, for each place cell we correlated the firing rate maps for consecutive sessions (O1 & O2, O2 & C1, C1 & C2 and C2 & O3). Remapping was defined as a change of spatial firing pattern at an above-chance level where chance was determined using a shuffle procedure. For this, for every sequence of sessions with more than 10 simultaneously recorded place cells (12 sequences for Closed-Door, 9 for One-Way) we correlated the rate maps of random cells between each session and the next, a thousand times. These shuffled distributions were very similar and thus combined to give four distributions, each one describing the chance of remapping between consecutive sessions. We then compared the between-session correlation values of all place cells to these combined shuffles; when the value for a given cell was lower than the 95^th^ percentile of the corresponding shuffle we considered that it had remapped between those two sessions. The rationale for shuffling within sequences was to ensure cells were only compared to others recorded under the same behavioural constraints. For every cell, this procedure yields four binary outcomes (remapping or no remapping between O1 & O2, O2 & C1, C1 & C2 and C2 & O3). For visualisation we generated a histogram of all 16 possible combinations.

### Remapping between foraging and goal-directed behaviour

After differentiating foraging from goal-directed behaviour (Methods - Behaviour discrimination) we sought to compare the firing of place cells between these two states. First, for each place cell we correlated its foraging and goal-directed firing rate map in each session (O1, O2, C1, C2 and O3). Remapping was defined as a change of spatial firing pattern between these at an above-chance level where chance was determined using a shuffle procedure similar to that described above (*Methods - Remapping between sessions*), but comparing foraging maps to goal-directed maps of random cells within the same session. We then compared the distribution of observed correlation values from all place cells to these combined shuffles for each session. If the median observed correlation value fell above the 95^th^ percentile of the corresponding shuffle distribution we considered that cells were more stable between foraging and goal-directed modes than chance.

### Firing rate remapping

In addition to the correlation-based analyses described in the main text, we also looked at rate remapping (changes in place cell firing rate independent of spatial changes) using a method described previously (Fuhs et al. 2005). Briefly, this followed the same procedure as the correlation analyses but using the following difference metric in place of the Pearson *r* correlation:

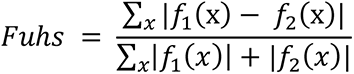

where the variable *x* ranges over map locations, and *f*_1_(*x*) and *f*_2_(*x*) are two place field firing rate maps that have been zero-normalized by subtracting off their respective mean firing rates. As outlined by Fuhs and colleagues (2005), the metric calculates the ratio of the difference between rate maps to the difference between each rate map and its mean. Thus, the metric equals 0 for identical fields and approaches a maximum of 1 when maps differ either spatially or in terms of firing rate. We used this ‘Fuhs metric’ to compare firing rate maps between sessions and only for those place cells which were found not to significantly spatially remap between any sessions (see *Methods - Remapping between sessions*).

### Place fields

When detecting place fields, we used smoothed, speed-filtered (>5cm/s) firing rate maps generated only from foraging data, as described in Methods: *Firing rate maps and spatial information*. Firing rate maps were thresholded to 20% of their maximum value, place fields were defined as regions within the thresholded maps with:

1. an area greater than 9 contiguous bins and;
2. a peak firing rate greater than 1Hz.

For each field we extracted its area (number of bins in the region), weighted centroid (center of the region based on bin locations and firing rates), convex hull (coordinates of minimum enclosing polygon) and mean firing rate (average bin value, all values calculated using Matlab *regionprops*).

### Place field repetition

To quantify the prevalence of place field repetition, for every place cell we calculated the average correlation between all possible pairs of boxes for each session independently. We compared the resulting distributions to chance, where chance was determined using a shuffle procedure similar to that described above (*Methods - Remapping between sessions*) but comparing box firing rate maps to box maps of random cells within the same session. Additionally, for session O2 only, we repeated this procedure after dividing place cells into groups based on the number of place cells they exhibited in this session.

For all sessions the distribution of observed correlations differed from the shuffles and was significantly shifted towards 1. This result suggests greater place field repetition than chance but does not quantify the degree of place field repetition. For this we designed shuffles to test if the observed values reflected place field repetition in 2, 3 or 4 boxes. For each shuffle we took each cell in turn and collected all 4 compartments for session O1. We next selected a random compartment and duplicated this across multiple compartments (2, 3 or 4 depending on the shuffle type). For duplication we took the rate maps for that compartment across the different sessions. For example, a shuffle designed to reflect repetition in two compartments might include compartment A, B and C in session O1 and compartment A from session O2. Our reasoning was that the same compartment sampled in different sessions provides a good approximation for sampling the same field in multiple compartments (assuming place field repetition).

### Place field overrepresentation

Next, to test if more or fewer fields were found around the locked doors, for each session we found the total number of fields with their centroid less than 25cm from each door. For the Closed-Door sequence we then calculated the average number of fields around the 3 open doors and compared this to the number of fields around the locked door. If these were equally represented we would expect 50% of this total to fall around each door type in each session, which we tested using a chi-square test of equal proportions (custom Matlab function, *p*-values were corrected across sessions using the Holm-Bonferroni method). For One-Way sequences we used a similar procedure except that we compared the number of fields within 25cm of the locked sides of the doors to the open sides.

To test more generally if doorways were overrepresented relative to the boxes and dummy doors, for each session and for each sequence type separately we counted the number of fields falling within 25cm of any door, any box centre or any dummy door and expressed these values as proportions of the total of the three (*N*). We then calculated the total surface of each test area by taking the median across all dwell maps and thresholding the resulting map to discard bins with a median of less than 0.01s. Surface area was then calculated as the total remaining bins within 25cm of the test areas. The expected proportion of fields for each test area was then estimated as the proportion of total surface area in the dwell map included in this test area multiplied by *N*. Lastly, we tested the observed proportions against the expected proportions using a chi-square test of equal proportions (custom Matlab function as above).

### Place field firing rate and area

To further test if field properties such as mean in-field firing rate and total field area (Methods: *Place fields*) changed in response to a change in connectivity we extracted all place fields with a weighted centroid within 25cm of a doorway and calculated average values for each session.

Next, we separated fields into 2 groups; those with a weighted centroid within 25cm of a door (or side of a door in the One-Way sequence) which remained open throughout the sequence and those within 25cm of one which was locked during the sequence. For visualisation we rotated fields in the Closed-Door sequences so that the locked doorway was always at the bottom and for the One-Way sequences we flipped the fields along the x-axis so that fields on the locked or open side of a door would overlap. For each spatial bin we then counted the total number of overlapping fields observed in that bin and plotted the result as a density heatmap. Finally, we calculated the average area and mean in-field firing rate for each group in each session and sequence.

### Place field extent across doors

To test if fields that extended across doorways continued to do so when these doors were locked we found all of the fields within 25cm of a door that was locked in C1. We then found the area (total bins) found on each side of the door (a and b) and calculated a ‘bridge index’ as:

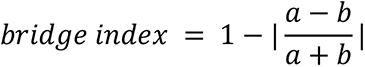

This index varies from 0 when a field’s area is entirely on one side of the door to 1 when a field’s area is equally distributed on each side of the door. For visualisation we then projected the field convex hull onto an axis perpendicular to the locked door and extracted the maximum limits of this projection and the centroid along this axis.

### Bayesian decoding of box quadrants

Position decoding was performed using a memoryless Bayesian algorithm that assumes spiking is Poissonian, independent across neurons, and compares the spiking vector of simultaneously recorded cells to their expected firing rates given by their rate maps (Kaefer et al., 2020; Zhang et al., 1998). We used rate maps constructed from O1 when decoding position from neural activity in session O2 and O2 rate maps were used for C1 decoding, meaning the ‘test’ and ‘training’ data always belong to different datasets.

The position and spike data of place cells were discretized into τ = 300 ms windows. Windows in which the animal had a velocity less than 5 cm/s were removed to reduce contamination by the non-local reactivations that can arise during hippocampal replay. Only sessions with at least 15 simultaneously recorded place cells were considered (Closed-Door, n = 9; One-Way, n = 3). The number of spikes that occurred in a given cell during a decoding window is denoted as *σ_i_* and thus the vector of spiking activity all simultaneously recorded cells 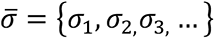. For every position *x* and place cell *i*, the expected value for the firing rate 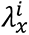 is retrieved from the rate map. Using the Poissonian assumption, the probability of observing spike vector 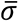 given *x* is:

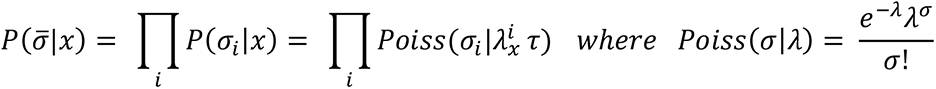

And via Bayes’ rule:

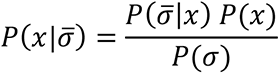

By using a uniform prior distribution *p*(*x*), and enforcing 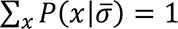, the decoded position becomes:

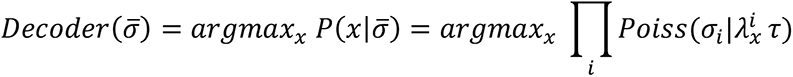

The decoded positions and true positions were assigned to a box and a quadrant using a set of inequalities based on linear equations centred around the centroid coordinates of the compartments. For each session we generated a 16 x 16 confusion matrix of actual vs predicted locations, wherein each cell is populated using:

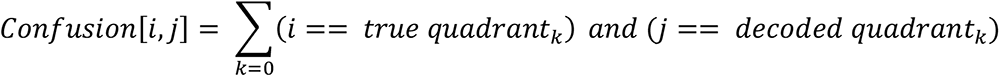

A mean confusion matrix was computed for each session type (O2 Closed-Door, O2 One-Way, C1 Closed-Door, C1 One-Way). The rows of the session confusion matrices were converted to probability distributions by dividing by the sum of the row.

### Cluster quality measures

Cluster quality (Sup. Table2) was estimated by calculating isolation distance (N. Schmitzer-Torbert et al. 2005; Neil Schmitzer-Torbert and Redish 2004), *L_ratio_* and peak waveform amplitude, taken as the highest amplitude reached by the four mean cluster waveforms in the session with the highest firing rate. For cluster *C*, containing *n_C_* spikes, isolation distance is defined as the squared Mahalanobis distance of the *n_C_* ^th^ closest non-*C* spike to the center of *C*. The squared Mahalanobis distance was calculated as:

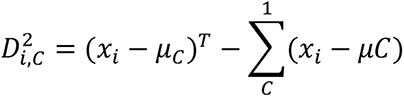

where *x_i_* is the vector containing features for spike *i*, and *μ_C_* is the mean feature vector for cluster *C*. A higher value indicates better isolation from non-cluster spikes. The *L* quantity was defined as:

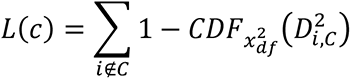

where *i ∉ C* is the set of spikes which are not members of the cluster and 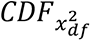 is the cumulative distribution function of the distribution with 8 degrees of freedom. The cluster quality measure, L_ratio_ was thus defined as *L* divided by the total number of spikes in the cluster.

### Statistical analyses

All statistics were performed in Matlab (The Mathworks, 2019a). Omnibus tests included two-way repeated measures ANOVAs (Matlab *ANOVAN* with repeated measure included as a random variable) and two-way ANOVAs (Matlab *ANOVAN*). The most common two-way repeated measures ANOVA design consisted of door type (closed/open) as the between subject variable, session (O1, O2, O3 …) as the within subject variable and animal as the repeated measure. Example responses included door pushes or correlations. These tests included two-way interactions. Two-way ANOVA designs were similar but without the repeated measure; often these were used for electrophysiology data containing missing values (i.e. cells not active in every session). In all cases, post-hoc tests were based on estimated marginal means and corrected using the Tukey-Kramer method (Matlab *multcompare*). For post-hoc tests the repeated measure was treated as a fixed instead of random variable but we only considered comparisons between groups and not across the repeated measure of sessions.

Other tests included one-way ANOVAs (Matlab *anova1*), one- and two-sample t-tests (Matlab *ttest* and *ttest2*), two-sample Kolmogorov-Smirnov tests (Matlab *kstest2*) and chi-square tests of expected proportions. Chi-square tests were calculated as:

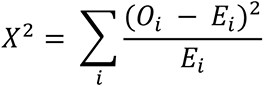

where O are the observed values and *E* are the expected values. The degrees of freedom, *c*, are N-1 where N is the number of expected values; *p* was calculated as one minus the value of the Chi-square distribution at *X*^2^ with degrees of freedom *c* (Matlab *chi2cdf*).

All tests were two-sided unless otherwise stated. Where multiple t-tests or chi-square tests were used to compare grouped values to chance we controlled the family-wise error rate by correcting the *p*-values using the Holm-Bonferroni method (Holm 1979). Briefly, *p*-values were ranked in ascending order and each value was then compared to a corresponding cutoff calculated as:

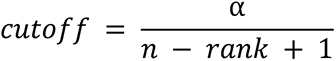

where α was the target significance threshold which was always set to .05 and *n* was the total number of *p*-values to correct. Each *p*-value was then compared to this cutoff in turn, the first *p*-value that exceeded its cutoff and all following *p*-values were considered to be non-significant and corrected to .99.

## Acknowledgements

We would like to thank Marta Huelin for providing one of the experimental subjects and members of the Institute of Behavioural Neuroscience in London and of the Department of Psychological and Brain Sciences in Dartmouth for support and discussions.

## Author contributions

Éléonore Duvelle: Project administration, Supervision, Writing—original draft preparation, Investigation, Methodology, Conceptualization, Writing—review & editing.

Roddy Grieves: Visualization, Data curation, Resources, Formal analysis, Software, Writing—original draft preparation, Writing—review & editing.

Anyi Liu: Investigation, Methodology, Visualization, Writing—original draft preparation, Writing—review & editing.

Selim Jedidi-Ayoub: Investigation.

Joanna Holeniewska: Investigation.

Adam Harris: Formal analysis, Writing—review & editing.

Nils Nyberg: Investigation.

Julie Lefort: Methodology, Writing—review & editing.

Francesco Donnarumma: Formal analysis.

Kate Jeffery: Resources, Methodology, Writing—review & editing.

Christopher Summerfield: Funding acquisition, Methodology, Conceptualization, Writing—review & editing.

Giovanni Pezzulo: Funding acquisition, Methodology, Conceptualization, Writing—review & editing.

Hugo J. Spiers: Funding acquisition, Supervision, Methodology, Conceptualization, Writing—review & editing.

## Data and code availability

The data and analysis code used for this experiment will be stored and made available after publication on an online repository.

## Supplementary data

A supplementary video of door-pushing and detour behaviour can be found <u>here</u>.

Sup. Fig.1 shows a schematic and pictures of the experimental room.

**Supplementary Figure 1:**
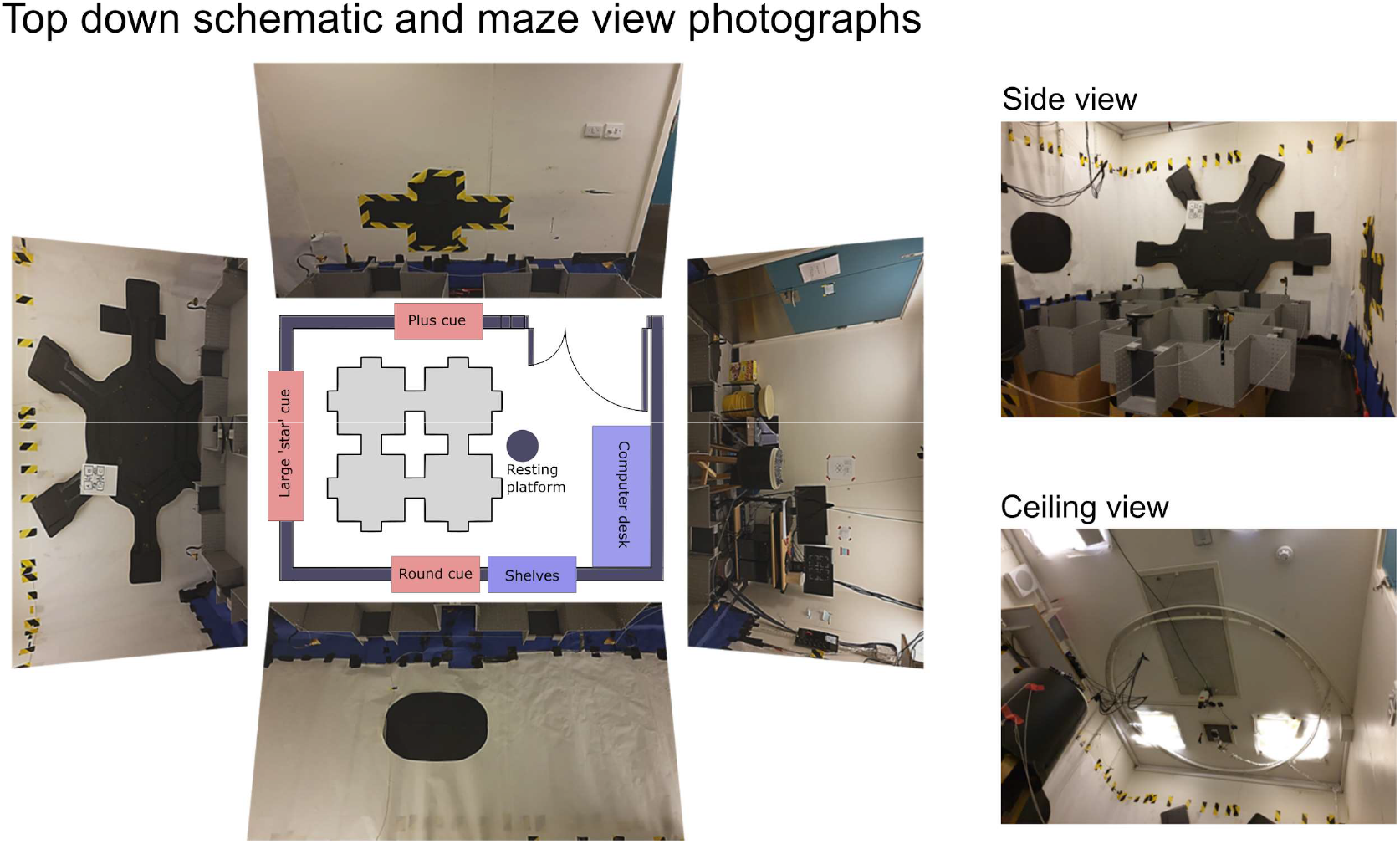
Experimental room layout; Left: top down schematic of the experimental room, main pieces of furniture and visual cues are labelled. The position and shape of the maze is given by the grey shape in the middle. The location of the resting platform is also given by the nearby blue circle. Surrounding this schematic are photos taken from the perspective of the maze along each major axis. Right: additional photographs showing a more natural side view of the setup and a view of the ceiling.

### Individual rat behaviour

Door-pushing behaviour organised as in Fig. 2A-D but averaged over individual rats instead of sessions is shown in Sup. Fig.2. The same effects still hold (Sup. Table1 for statistics). Sup. Fig.3 shows first door choice and first foraging choice organised as in Fig. 2E-J but with Closed-Door and One-Way sessions separately. Results do not obviously differ between sequence types except that performance seems generally increased for One-Way.

**Supplementary Figure 2:**
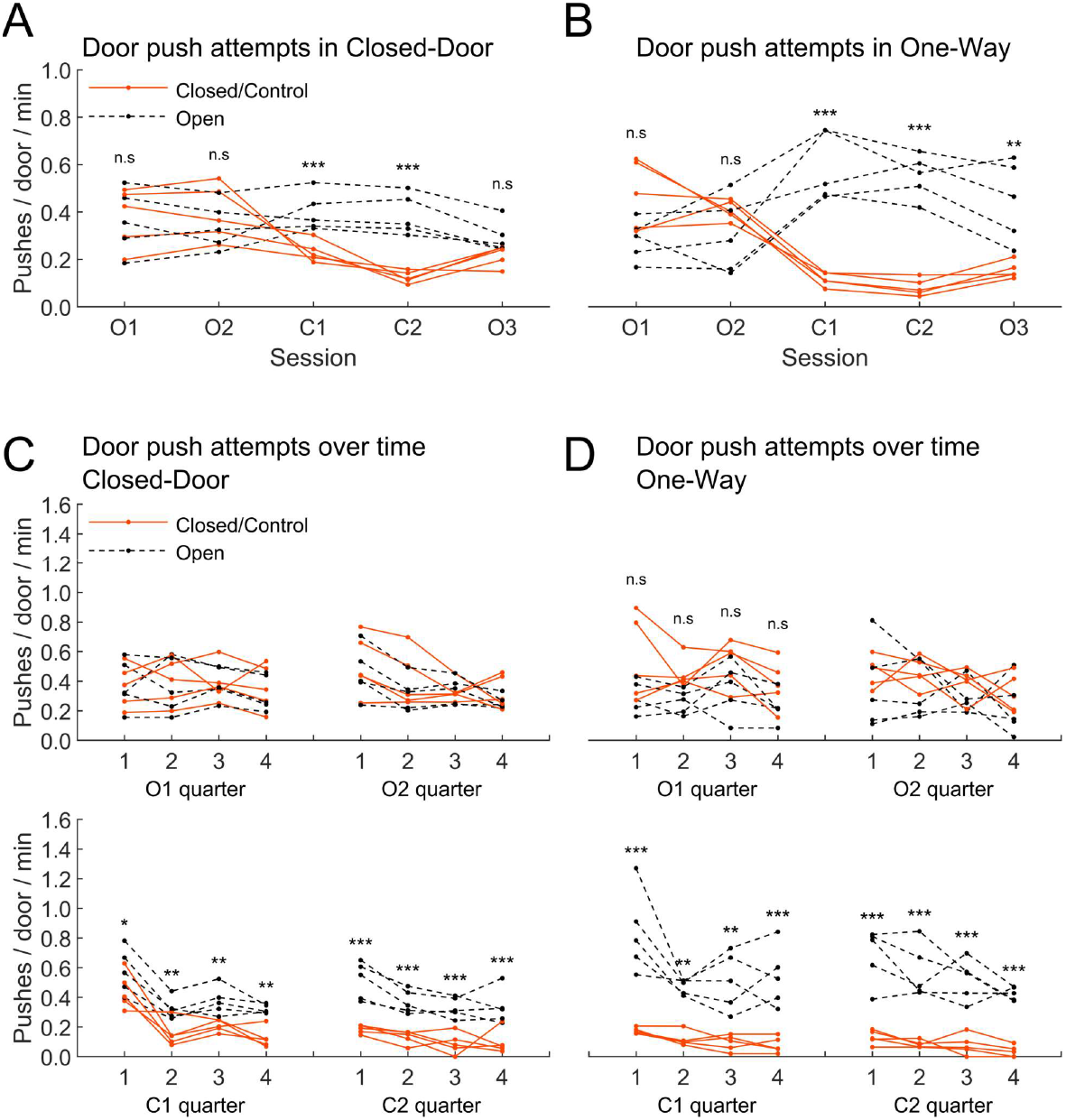
Knowledge of the door status and partial knowledge of the task. This is a supplement to Fig. 2A-D showing data averaged by rat instead of sessions. Here and later, * :*p* < .05, ** : *p* < .01, ***: *p* < .001. Statistics can be seen in Sup. Table1. **(A)** Normalised (per door and minute) number of door pushes for either the control doors (in O1-3, doors that will be or were locked), closed doors (same doors but in C1 and C2), or open doors, in the Closed-Door sequence, averaged for each rat. **(B)** Same, for the One-Way sequence. We usually closed the direction preferred by the rats, if they had a bias, which is why the numbers of pushes on the control side often seem higher in O1 and O2. **(C)** Same as A but separated by session quarters and only showing sessions O1, O2, C1 and C2. (D) Same as C but for the One-Way sequence.

**Supplementary Figure 3:**
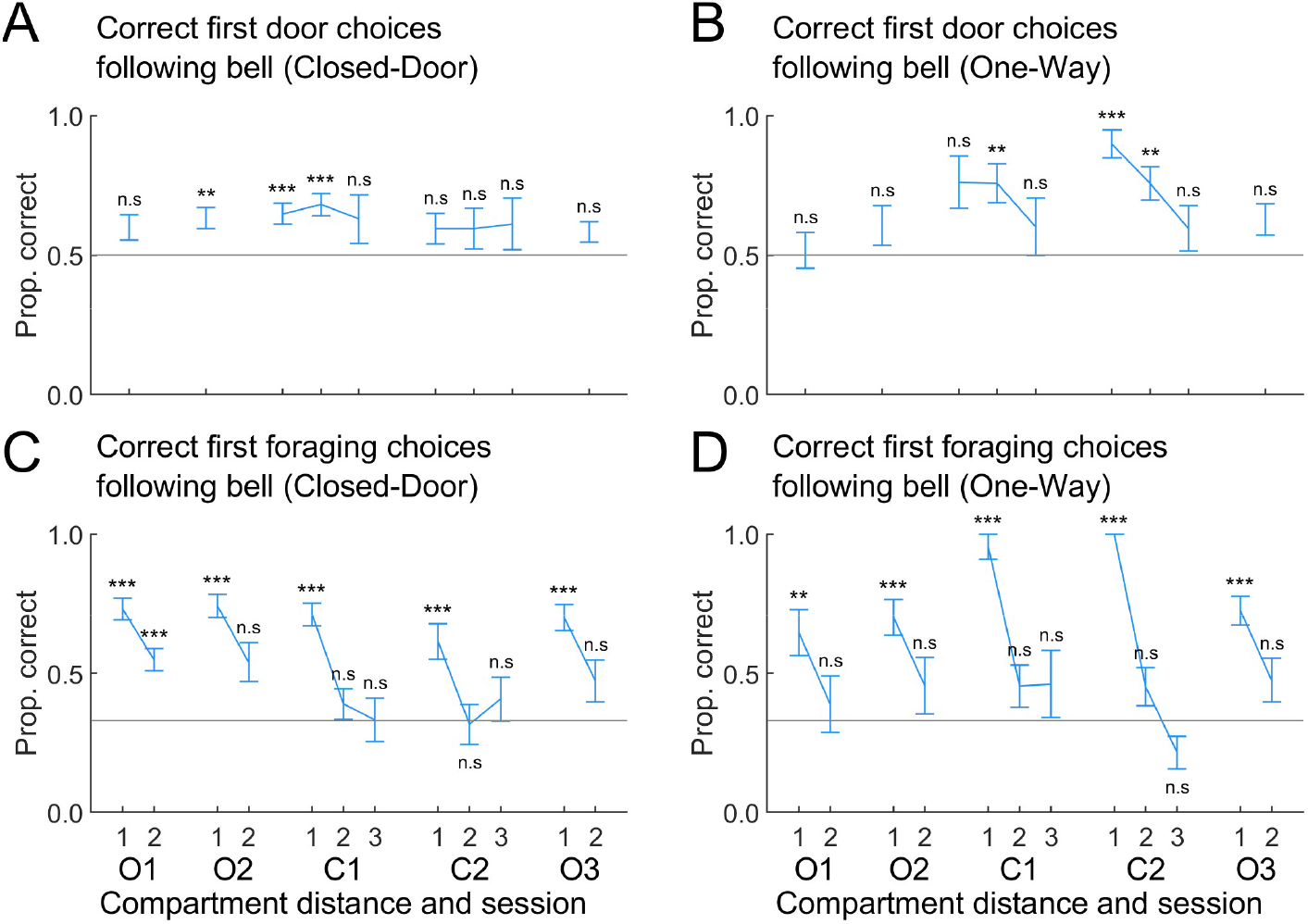
Knowledge of task parameters, separated by sequence type and averaged across sessions. This is a supplement to Fig. 2I-J. Statistics can be seen in Sup. Table1. Here and elsewhere, error bars indicate standard error of the mean. **(A)** Proportion of correct first door push after all bell sounds. Chance level is 50% (2 doors per box). Values are separated by distance between the current box and the goal box (distance of 1 = adjacent box). Distances of 2 and 3 are only available when a door was locked. **(B)** Same as A but for the One-Way condition. **(C)** Proportion of correct first foraging choices (where the rat forages first after a bell sound). Chance level is 33% when excluding the initial box. Rats foraged in the correct box for a goal distance of 1 (and 2 in O1), otherwise the proportions were not significantly different from chance. **(D)** Same as C but for the One-Way condition.

**Supplementary Table 1:**
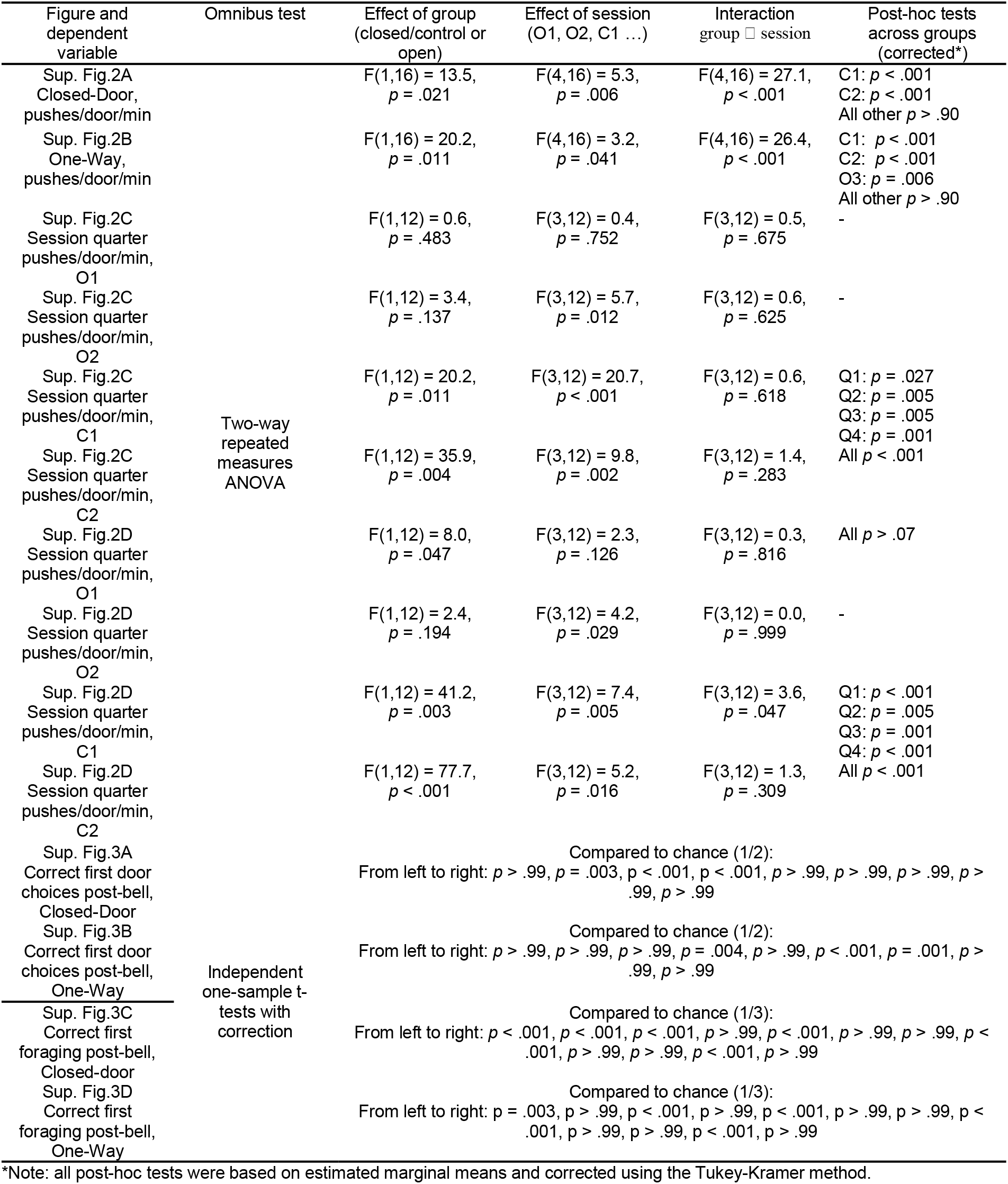
Statistical test results corresponding to Sup. Fig.2 and Sup. Fig.3.

### Occupancy maps

For each session in each sequence we calculated the average dwell time map, in which bins with an average occupancy less than 0.01s were considered empty. Before averaging, maps were rotated so that the closed door was always found in the south position (in the case of Closed-Door sequences) or reflected around the y-axis so that all doors were locked in a clockwise direction (in the case of One-Way sequences). We repeated this procedure for dwell time maps created using only foraging and goal-directed data respectively (see *Methods - Behaviour discrimination* for the automated classification method of trajectories). Dwell time maps averaged over all sessions are shown in Sup. Fig.4. Note the apparent similarity of maps across conditions for foraging but the difference between open and closed sessions in goal-directed maps, especially for Closed-Door.

**Supplementary Figure 4:**
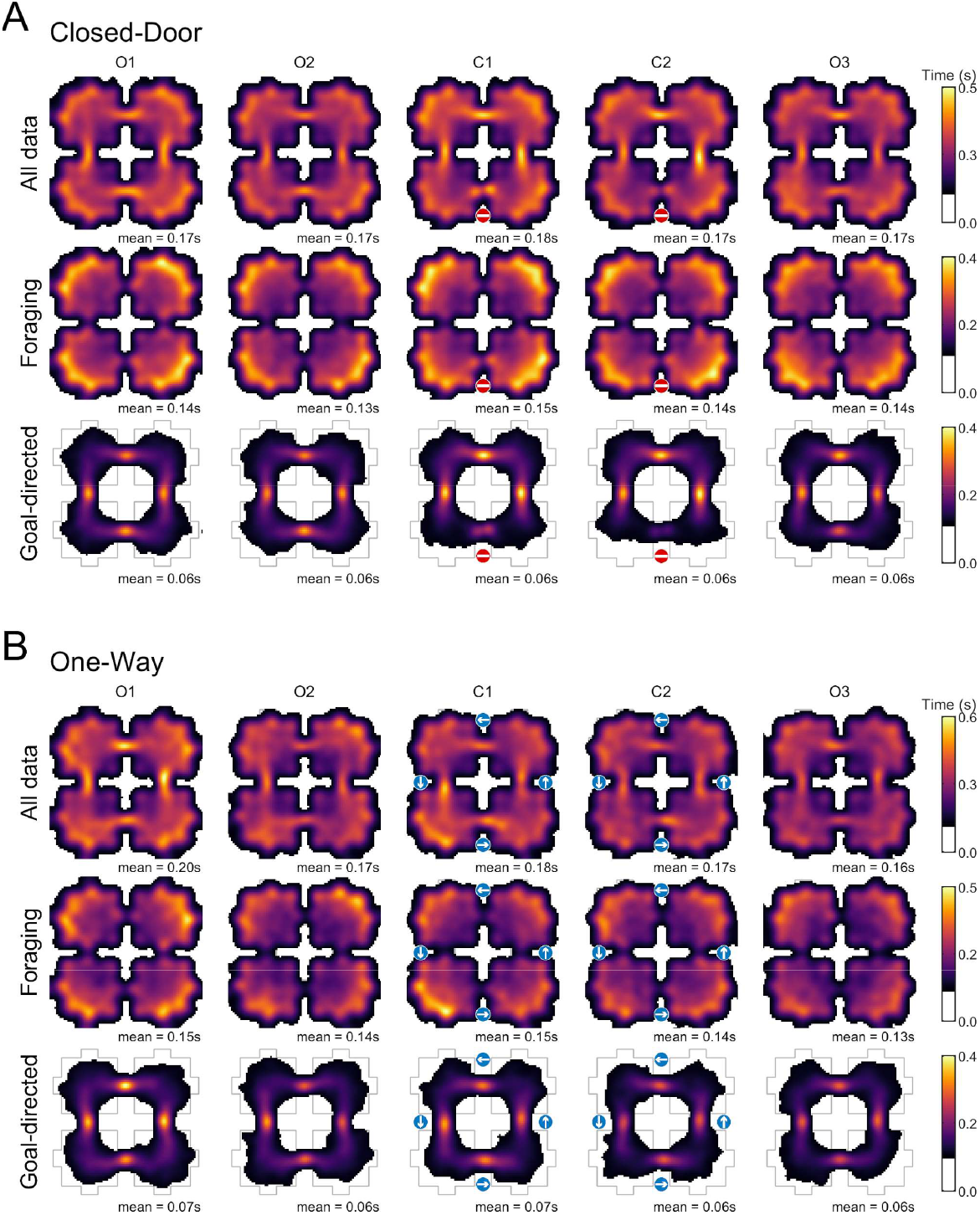
Occupancy maps, averaged across sessions. **(A)** Occupancy in the Closed-Door sequence for all data (top), foraging data (middle) and goal-directed data (bottom). **(B)** Same as A but for the One-Way sequence. In both cases note the difference in time distribution between foraging and goal-directed behaviour epochs, especially around the closed door for Closed-Door.

### Cluster quality measures

See *Methods - Cluster quality measures* for detailed methods regarding these metrics.

**Supplementary Table 2:** Cluster quality measures for putative place cells

| Condition | Isolation distance | L ratio | Waveform height ($\mu\text{V}$) | Waveform width ($\mu\text{s}$) | Mean firing rate (Hz) |
| --- | --- | --- | --- | --- | --- |
| Closed-Door | $23.6 \pm 24.2$ | $0.03 \pm 0.06$ | $48.4 \pm 39.5$ | $485.1 \pm 65$ | $0.73 \pm 0.76$ |
| One-way | $20.2 \pm 13.4$ | $0.04 \pm 0.1$ | $45.5 \pm 32.4$ | $481.9 \pm 73$ | $0.65 \pm 0.63$ |

Note: values are mean ± standard deviation.

### Field distance to doorway

To track the position of place fields across sessions (Sup. Fig.5) we applied k-means clustering to the field weighted centroids (Matlab *evalclusters* with gap evaluation at [1…N] clusters where N was the greatest number of unique fields observed in a session). When analysing the shift of fields between sessions we excluded fields that moved more than 60cm (the side length of a box).

Lastly, we tested if fields shifted their positions relative to the locked door between sessions. To test this we calculated the distance from every field in each session to every doorway. For Closed-Door sequences, we calculated the mean and standard deviation of the distance to the closest 16 fields around the three unchanging doors and compared this to the mean and standard deviation of the distances to the closest 16 fields around the locked door. For One-Way sequences we calculated the mean and standard deviation of the distance to the closest 8 fields on the closed side of each door and compared this to the mean and standard deviation of the distances to the closest 8 fields on the open side of each door. In a similar, but wider approach, we compared the distribution of field distances from the locked doors and from the open doors.

**Supplementary Figure 5:**
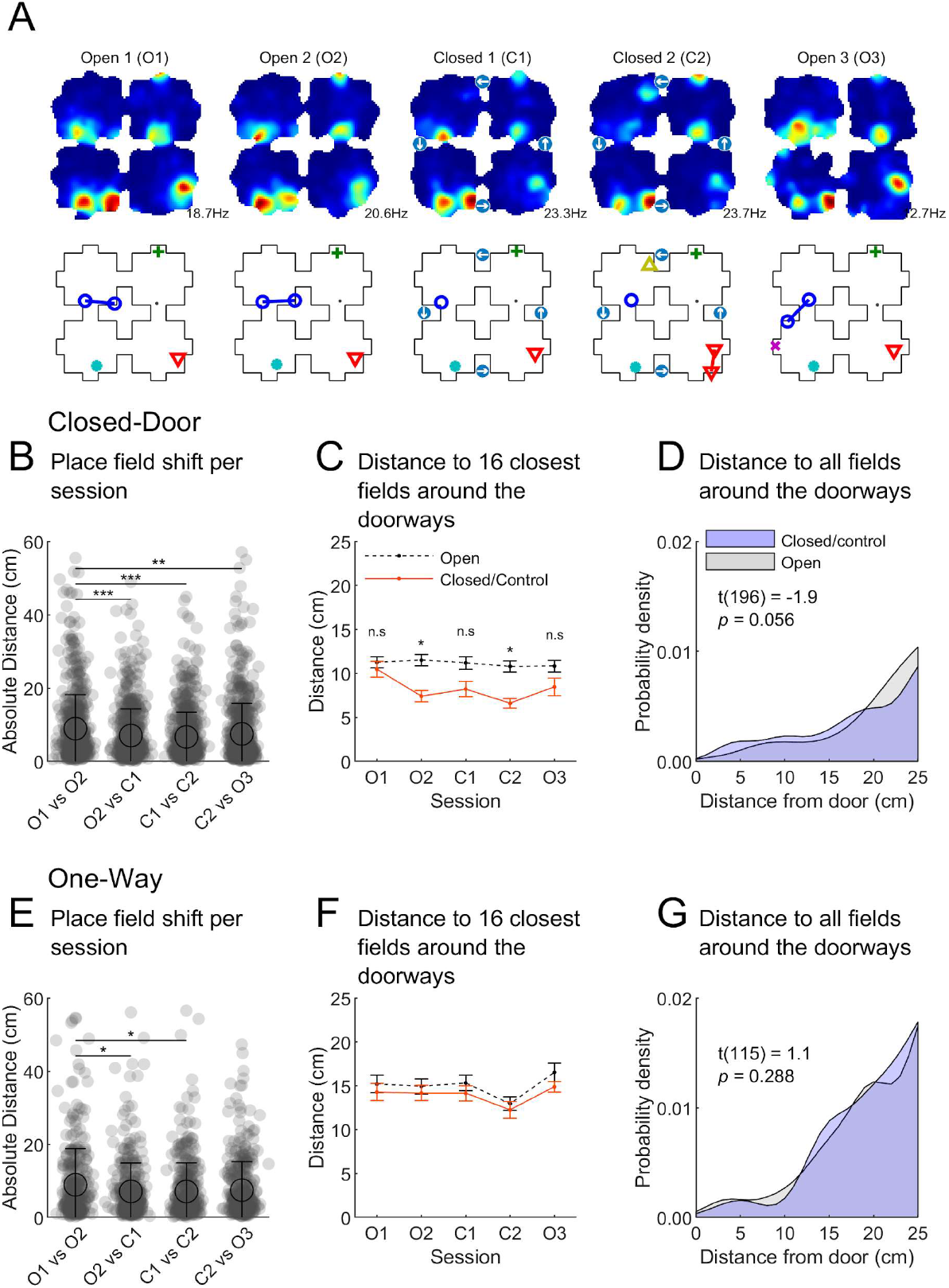
No change in distance of fields to doors with connectivity. See Sup. Table3 for statistics. **(A)** An example place cell recorded across sessions. Top: firing rate maps, blue denotes low or no firing, red denotes high firing. Maximum firing rates are given as text. Bottom: the centroids of place fields detected in each map. Each field is tracked across sessions and represented by the same coloured/shaped marker in each. Where a field was detected to split in one or more sessions the same marker is joined by a line of the same colour. **(B)** The absolute distance that place field centroids shifted between consecutive sessions. Fields that shifted more than 60cm (the side length of a box) were excluded. Place fields moved the most between sessions O1 and O2 where there was no connectivity change. **(C)** The distance to the closest 16 fields from the doors in each closed-door session. In O2 and C2 fields are significantly closer to the locked doors, although this is a general trend present in almost all of the sessions, including O2 and O3 where there was no connectivity change. **(D)** The distance to all fields from the doors in closed-door session C1 only. Text gives the result of a two-sample t-test comparing the distributions. **(E-G)** The same as B-D but for the One-Way condition with similar results.

**Supplementary Table 3:** Statistical test results corresponding to Sup. Fig.5.

| Figure and dependent variable | Omnibus test | Effect of group (closed/control or open) | Effect of session (O1, O2, C1 ...) | Interaction group □ session | Post-hoc tests across groups (corrected*) |
| --- | --- | --- | --- | --- | --- |
| Sup. Fig.5B<br>Field movements between sessions<br>Closed-Door | one-way ANOVA | - | $F(3,2684) = 10.3$,<br>$p < .001$ | - | O1vsO2 vs O2vsC1:<br>$p < .001$<br>O1vsO2 vs C1vsC2:<br>$p < .001$<br>O1vsO2 vs C2vsO3:<br>$p = .004$<br>All other $p > .20$ |
| Sup. Fig.5C<br>Field distance around doorways Closed-Door | two-way ANOVA | $F(1,310) = 27.2$,<br>$p < .001$ | $F(4,310) = 1.6$,<br>$p = .176$ | $F(4,310) = 1.3$,<br>$p = .281$ | O2: $p = .031$<br>O4: $p = .025$<br>All other $p > .30$ |
| Sup. Fig.5E<br>Field movements between sessions<br>One-Way | one-way ANOVA | - | $F(3,1537) = 4.0$,<br>$p = .008$ | - | O1vsO2 vs O2vsC1:<br>$p = .017$<br>O1vsO2 vs C1vsC2:<br>$p = .014$<br>All other $p > .07$ |
| Sup. Fig.5F<br>Field distance around doorways One-Way | two-way ANOVA | $F(1,310) = 3.4$,<br>$p = .067$ | $F(4,310) = 3.2$,<br>$p = .014$ | $F(4,310) = 0.1$,<br>$p = .984$ | - |

\*Note: all post-hoc tests were based on estimated marginal means and corrected using the Tukey-Kramer method.

### Minimal place field repetition & no connectivity effect

Sup. Fig.6 shows cross-boxes correlation distributions compared to shuffled data (*Methods - Place field repetition*) as described in *Results - Place cells encoded position in a global reference frame* for Closed-Door and One-Way. Results are similar for both sequence types.

**Supplementary Figure 6:**
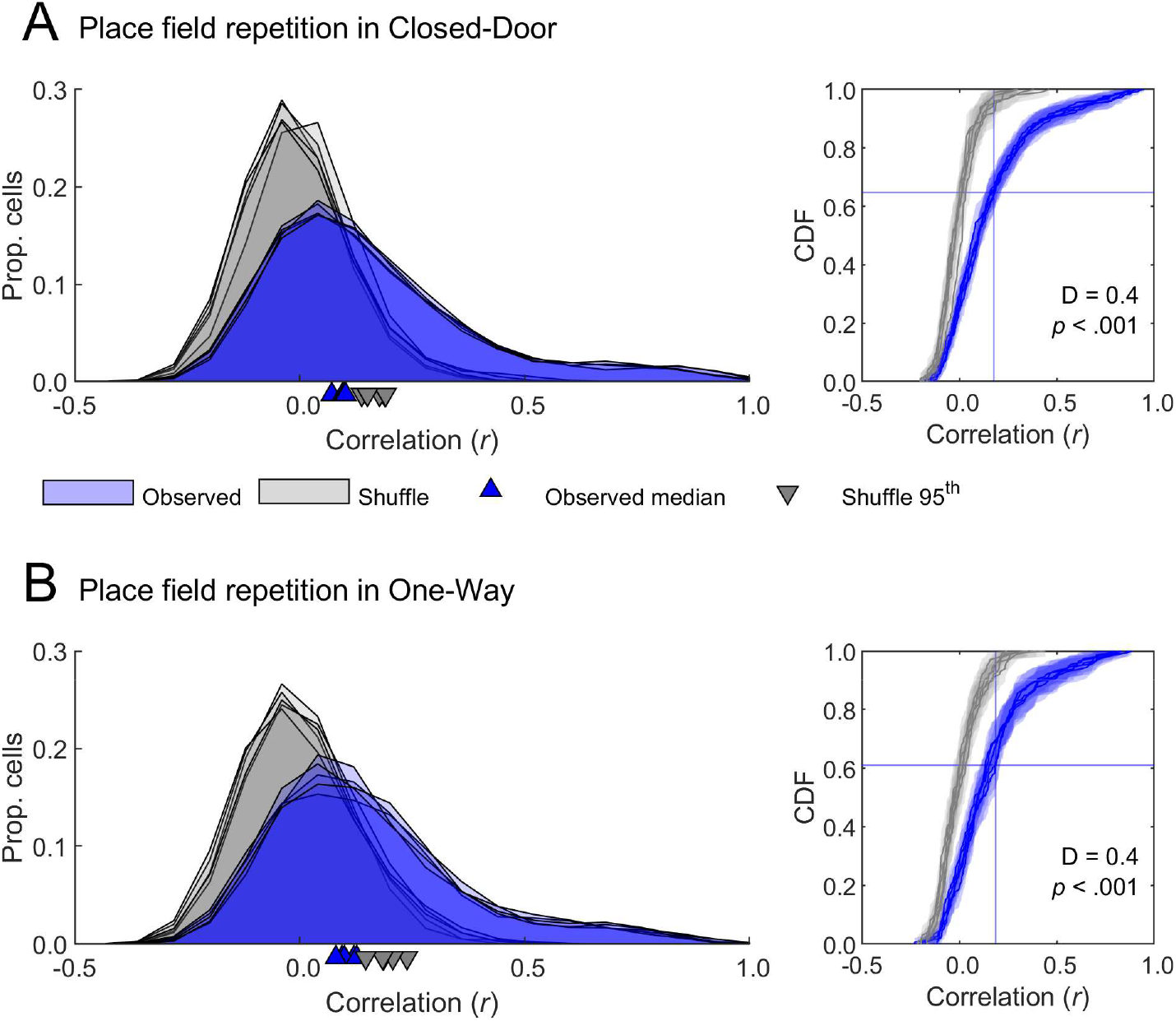
**(A)** Assessment of place field repetition for Closed-Door condition, *Left*, distributions of cross-box correlation values for all sessions (O1, O2, C1, C2, O3) in blue, with higher correlation values indicating place field repetition; in grey, distributions of shuffled correlation values for each session. Medians for all distributions are indicated by corresponding markers along the x-axis. The data median was always below the 95^th^ percentile of the corresponding shuffle, indicating similarity to the shuffle. *Right*, same data but showing the empirical cumulative distribution functions, shaded areas represent lower and upper confidence intervals. The 95^th^ percentile of the shuffle distribution (for O2) is indicated by a vertical blue line and the intersection of this with the data distribution is indicated by a horizontal line, showing that more than 60% of the cells do not repeat more than chance. Text gives the result of a two-sample Kolmogorov-Smirnov test comparing the two distributions in session O2. **(B)** Same as A but for One-Way sequences.

Sup. Fig.7 shows the equivalent of Fig. 9D-E for One-Way, with equivalent results.

**Supplementary Figure 7:**
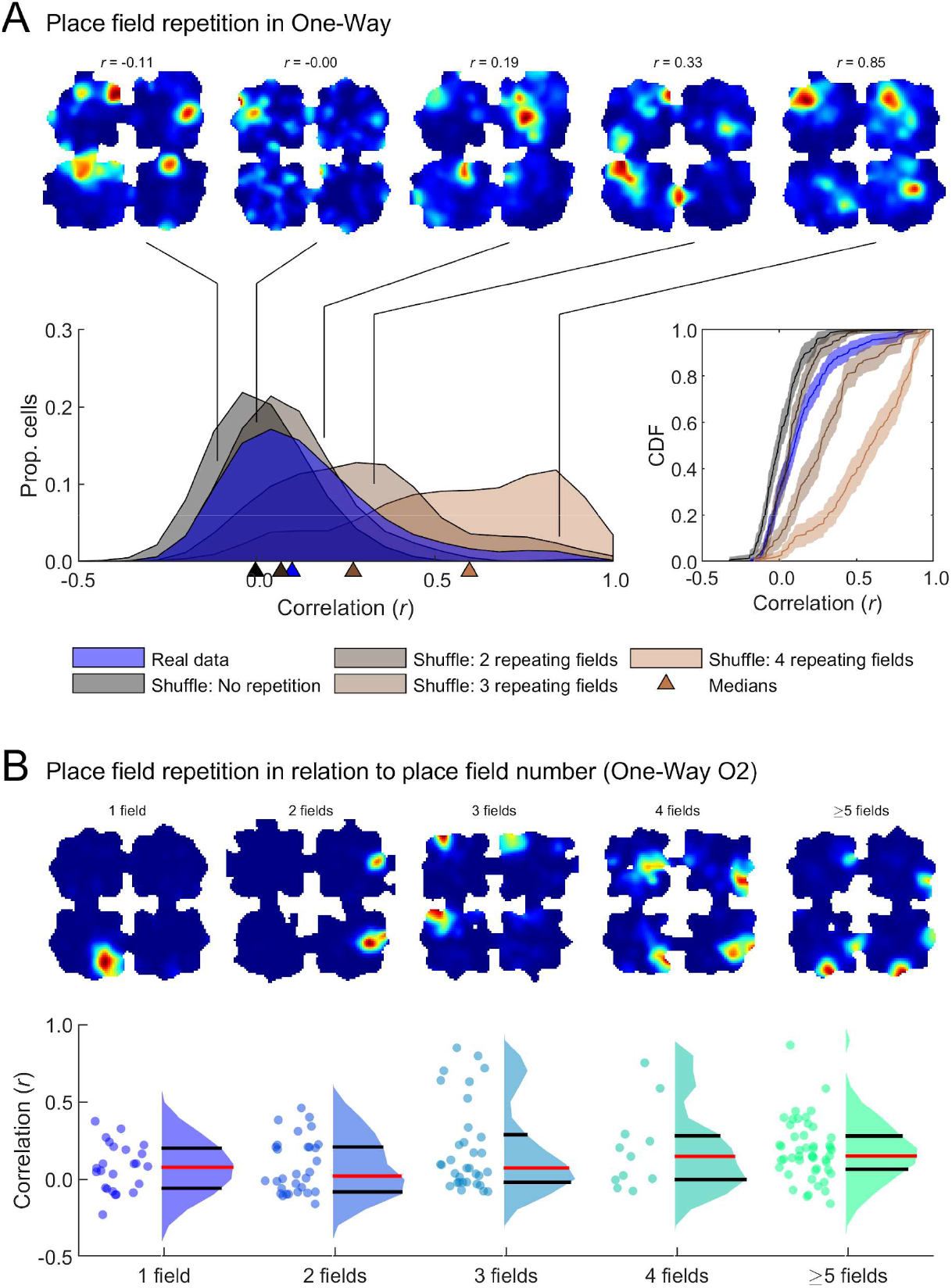
Place cells do not fully repeat in the One-Way condition. Supplement for Fig. 9D-E showing data for the One-Way condition. (A) *Top*: example rate maps (all data, speed-filtered, red indicates high firing rate, dark blue indicates low firing rate) with indicated correlation values on the O2 data distribution (blue). *Left*, blue distribution represents real data cross-box correlation values for session O2. All other distributions represent different shuffles designed to simulate the correlations expected if cells did not have any repeating fields or repeated fields in 2, 3 or 4 compartments. Medians for all distributions are indicated by corresponding markers along the x-axis. **Right**, same data as left but showing the empirical cumulative distribution functions, shaded areas represent lower and upper confidence intervals. The data most closely matched the distribution expected if cells had 2 repeating fields on average. **(B)** only for session O2. Top: example rate maps (as described in A) sorted by the number of detected place fields. *Bottom*: box correlations scores also separated by number of detected fields; dots show individual place cell data, distribution is shown as violin plot, with median in red and quartiles in black. High correlations indicative of place field repetition (>0.5) can be observed for only a small population of cells.

### Individual session results for Bayesian quadrant decoding

Sup. Fig.8 represents confusion matrices for all individual sessions used in the decoding analysis (*Methods - Bayesian decoding of box quadrants*), i.e., only sessions with at least 15 simultaneously recorded place cells. The diagonal pattern can be observed for most sessions, with some exceptions for those with lower numbers of cells.

**Supplementary Figure 8:**
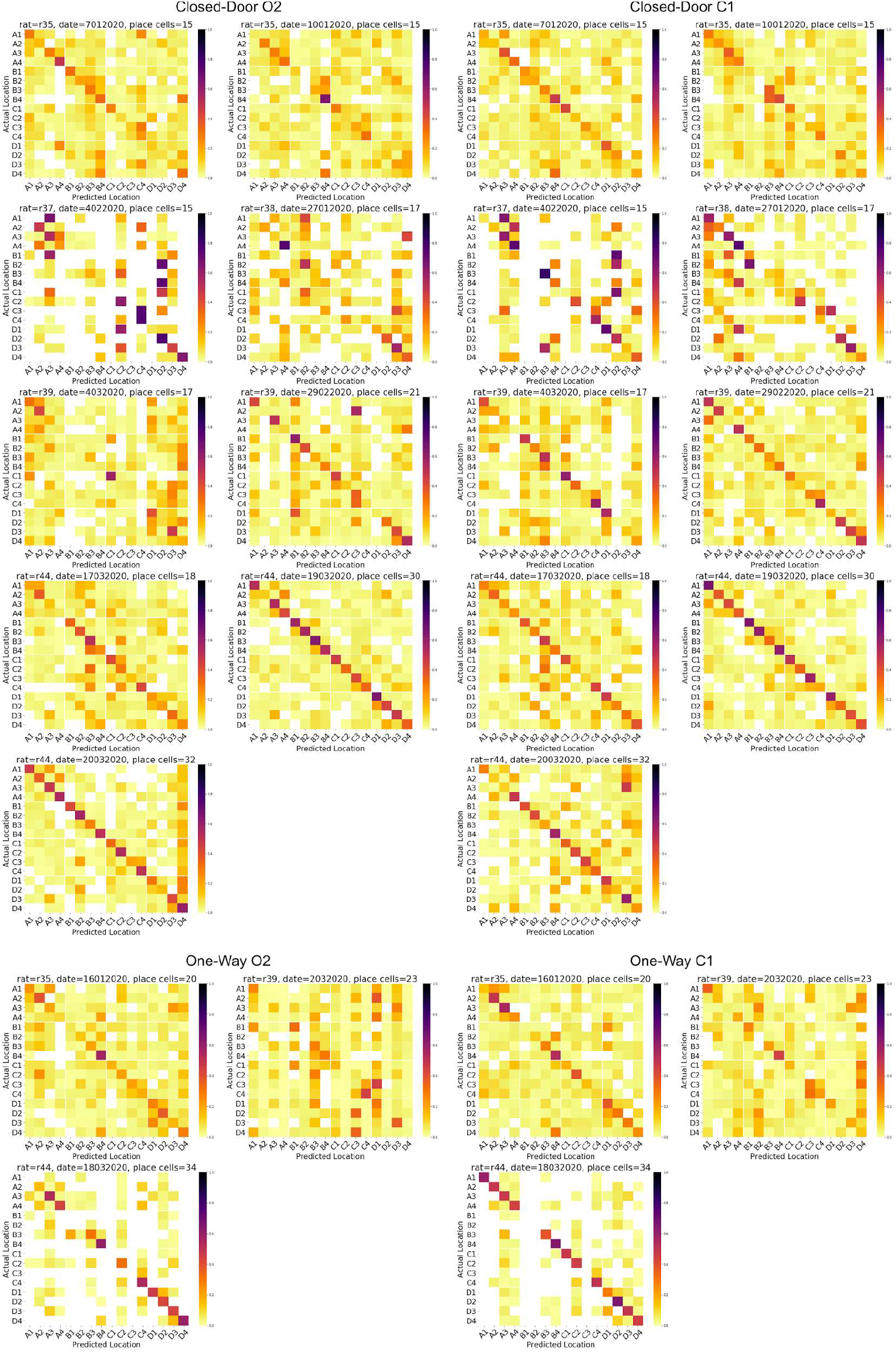
Individual confusion matrices for all sessions used in the quadrant decoding analysis, related to Fig. 10.

### The degree of connectivity of a box does not impact population activity

In a prior human neuroimaging study examining navigation of a simulation of Soho in London (UK), we found that participants entering a street that had greater connectivity than the preceding street had increased activity in their posterior right hippocampus, which decreased if the street had less connectivity (Javadi et al. 2017). We examined whether such a pattern of activity might be present in the activity of hippocampal place cells, comparing population activity when rats entered regions where the number of unlocked doors either increased or decreased. We found no evidence for modulation of the population activity driven by changes in connectivity (Sup. Fig.9).

**Supplementary Figure 9:**
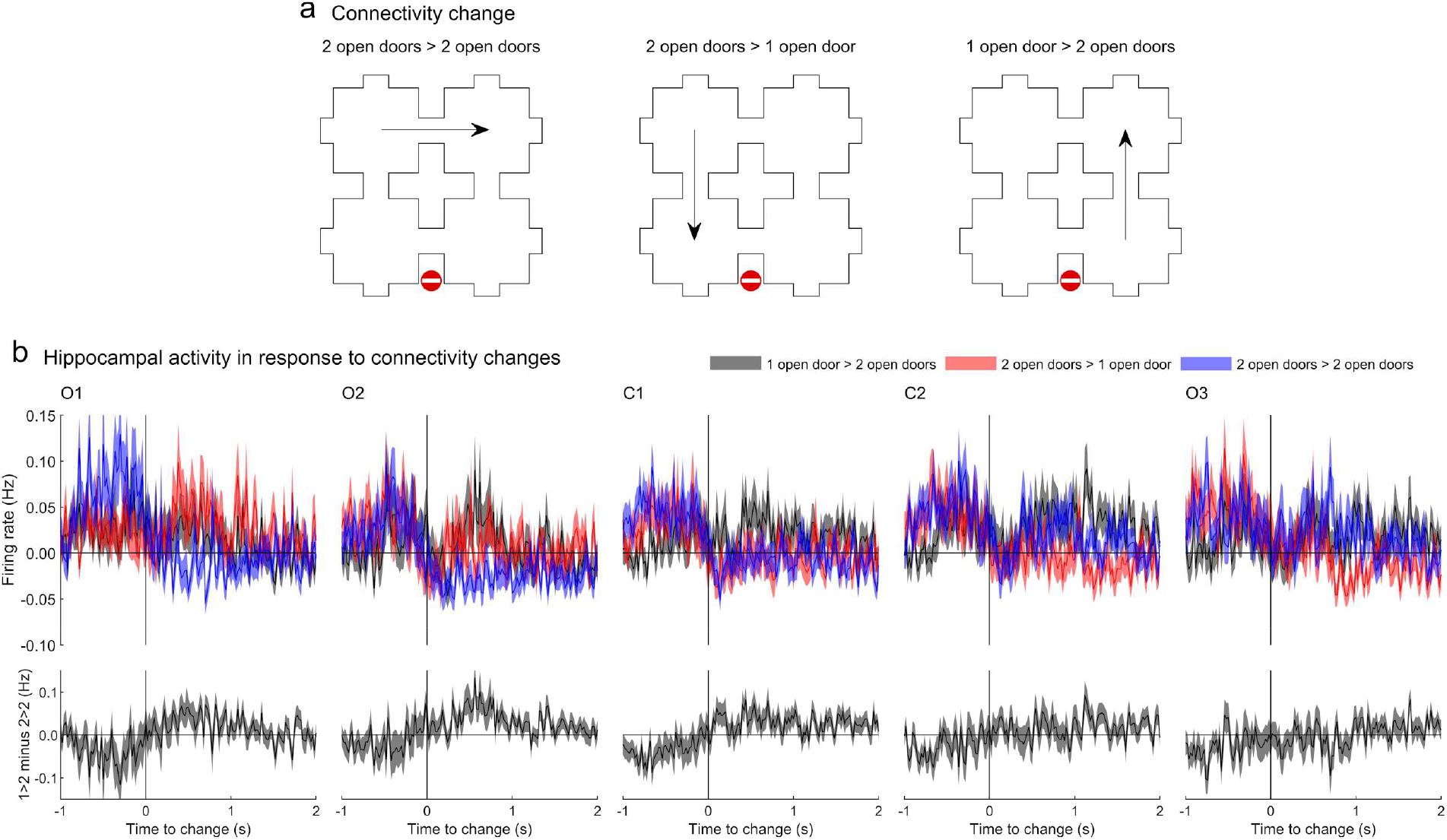
Activity when entering a box does not depend on the number of transitions available. (A) Schematic showing the possible connectivity transitions available to the rats when one door is closed. Rats can move to a box with equivalent connectedness (*left*), diminished connectedness (*middle*) or increased connectedness (*right*). (B) Top: mean and SEM firing rates for all place cells around these connectivity transitions (-1s to plus 2s). *Bottom*: the difference between the firing rate profile for increased connectedness and equivalent connectedness. Note that in the open sessions (O1, O2 & O3) all compartments are equally connected and these groups are arbitrary.

