## Supplementary figures 10 and 11 for "Hippocampal place cells encode global location but not changes in environmental connectivity in a 4-room navigation task"

**Duvelle et al., 2020**
**bioRxiv**

**Supplementary figures 10 and 11**

**Spike and rate maps for all used place cells**

The following figures show the spike plots and rate maps for all cells used in the analyses (only O2 and C1), built using only foraging, speed-filtered data, first for the Closed-Door condition, then for One-Way.

**Supplementary Figure 10**: Individual place cell plots for O2 and C1 in Closed-Door

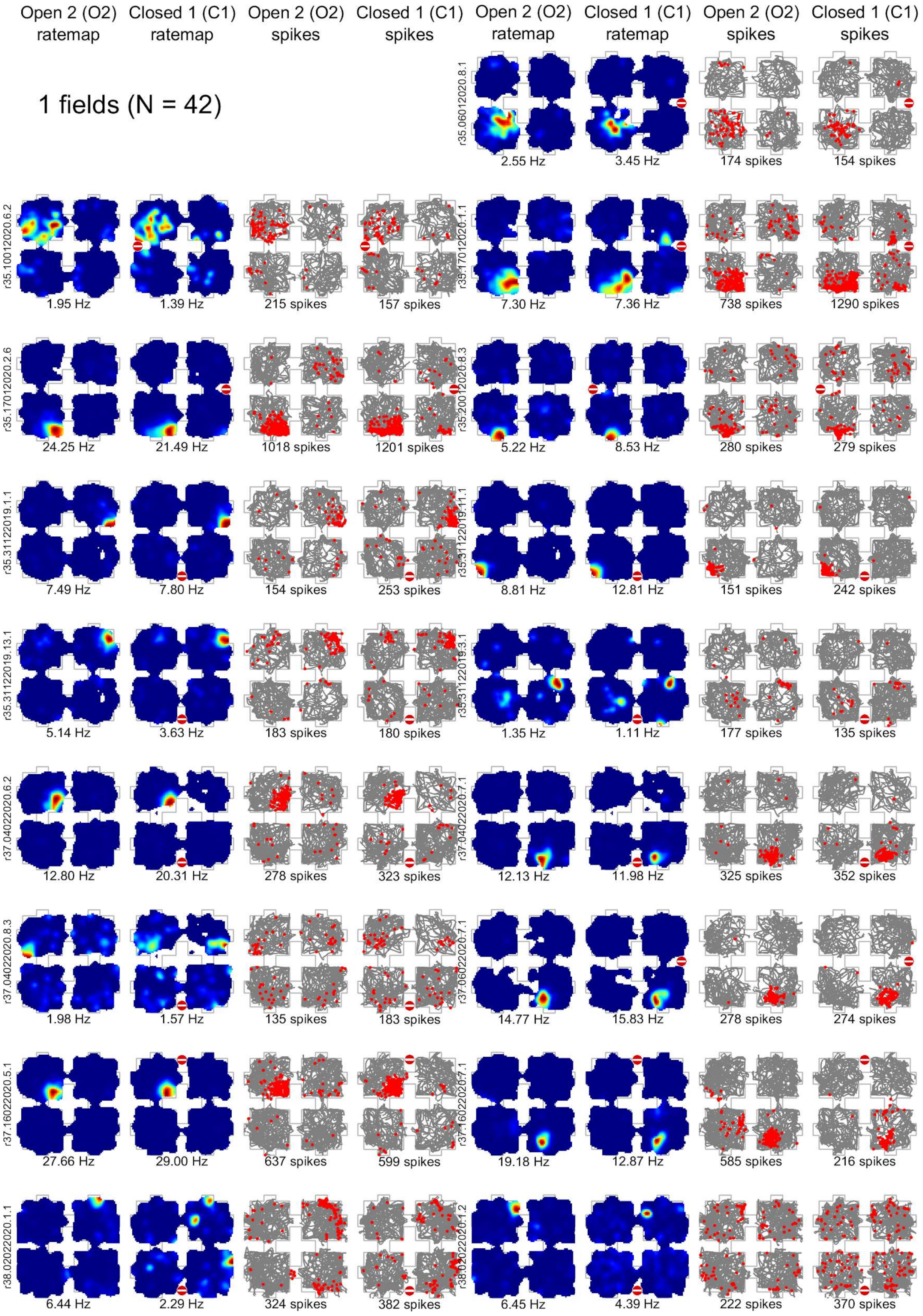

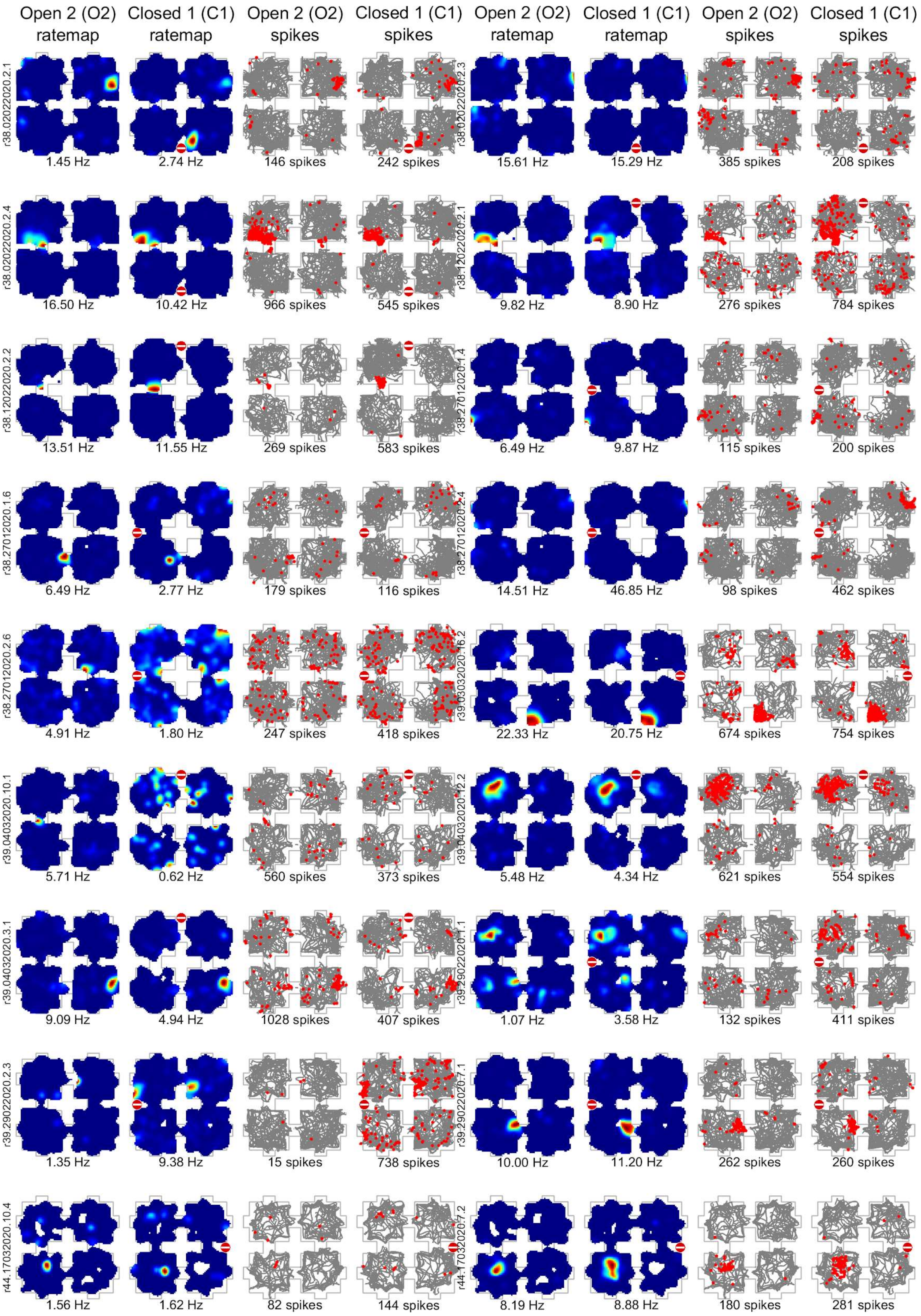

Open 2 (O2) ratemap
Closed 1 (C1) ratemap
Open 2 (O2) spikes
Closed 1 (C1) spikes
Open 2 (O2) ratemap
Closed 1 (C1) ratemap
Open 2 (O2) spikes
Closed 1 (C1) spikes
r38.02022020.2.1
1.45 Hz
2.74 Hz
146 spikes
242 spikes
r38.02022020.2.3
15.61 Hz
15.29 Hz
385 spikes
208 spikes
r38.02022020.2.4
16.50 Hz
10.42 Hz
966 spikes
545 spikes
r38.12022020.2.1
9.82 Hz
8.90 Hz
276 spikes
784 spikes
r38.12022020.2.2
13.51 Hz
11.55 Hz
269 spikes
583 spikes
r38.27012020.1.4
6.49 Hz
9.87 Hz
115 spikes
200 spikes
r38.27012020.1.6
6.49 Hz
2.77 Hz
179 spikes
116 spikes
r38.27012020.2.4
14.51 Hz
46.85 Hz
98 spikes
462 spikes
r38.27012020.2.6
4.91 Hz
1.80 Hz
247 spikes
418 spikes
r39.03032020.16.2
22.33 Hz
20.75 Hz
674 spikes
754 spikes
r39.04032020.10.1
5.71 Hz
0.62 Hz
560 spikes
373 spikes
r39.04032020.12.2
5.48 Hz
4.34 Hz
621 spikes
554 spikes
r39.04032020.3.1
9.09 Hz
4.94 Hz
1028 spikes
407 spikes
r39.29022020.1.1
1.07 Hz
3.58 Hz
132 spikes
411 spikes
r39.29022020.2.3
1.35 Hz
9.38 Hz
15 spikes
738 spikes
r39.29022020.7.1
10.00 Hz
11.20 Hz
262 spikes
260 spikes
r44.17032020.10.4
1.56 Hz
1.62 Hz
82 spikes
144 spikes
r44.17032020.7.2
8.19 Hz
8.88 Hz
180 spikes
281 spikes

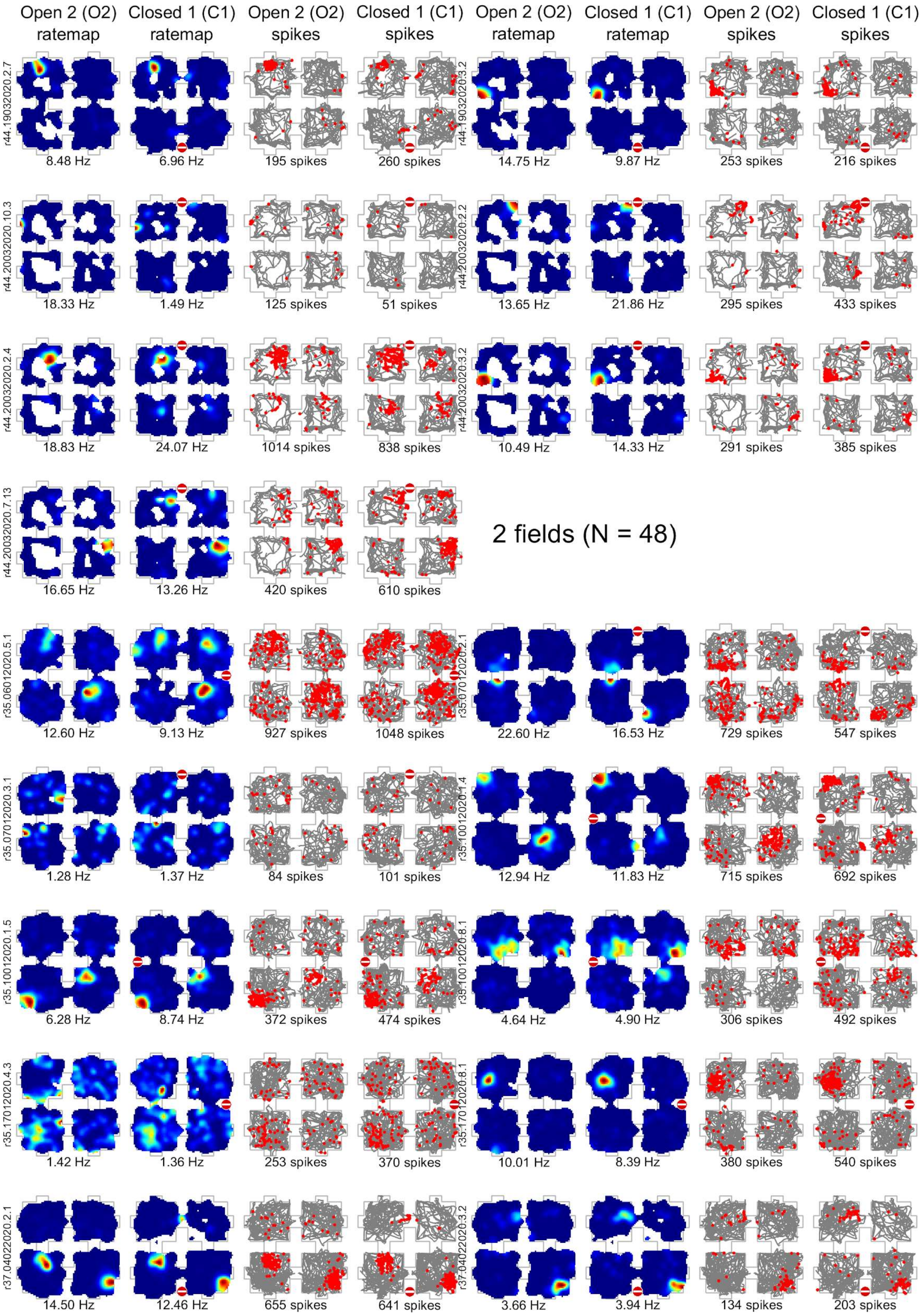
Open 2 (O2) ratemap
Closed 1 (C1) ratemap
Open 2 (O2) spikes
Closed 1 (C1) spikes
Open 2 (O2) ratemap
Closed 1 (C1) ratemap
Open 2 (O2) spikes
Closed 1 (C1) spikes
r44.19032020.2.7
8.48 Hz
6.96 Hz
195 spikes
260 spikes
r44.19032020.3.2
14.75 Hz
9.87 Hz
253 spikes
216 spikes
r44.20032020.10.3
18.33 Hz
1.49 Hz
125 spikes
51 spikes
r44.20032020.2.2
13.65 Hz
21.86 Hz
295 spikes
433 spikes
r44.20032020.2.4
18.83 Hz
24.07 Hz
1014 spikes
838 spikes
r44.20032020.3.2
10.49 Hz
14.33 Hz
291 spikes
385 spikes
r44.20032020.7.13
16.65 Hz
13.26 Hz
420 spikes
610 spikes
2 fields (N = 48)
r35.06012020.5.1
12.60 Hz
9.13 Hz
927 spikes
1048 spikes
r35.07012020.2.1
22.60 Hz
16.53 Hz
729 spikes
547 spikes
r35.07012020.3.1
1.28 Hz
1.37 Hz
84 spikes
101 spikes
r35.10012020.1.4
12.94 Hz
11.83 Hz
715 spikes
692 spikes
r35.10012020.1.5
6.28 Hz
8.74 Hz
372 spikes
474 spikes
r35.10012020.8.1
4.64 Hz
4.90 Hz
306 spikes
492 spikes
r35.17012020.4.3
1.42 Hz
1.36 Hz
253 spikes
370 spikes
r35.17012020.8.1
10.01 Hz
8.39 Hz
380 spikes
540 spikes
r37.04022020.2.1
14.50 Hz
12.46 Hz
655 spikes
641 spikes
r37.04022020.3.2
3.66 Hz
3.94 Hz
134 spikes
203 spikes

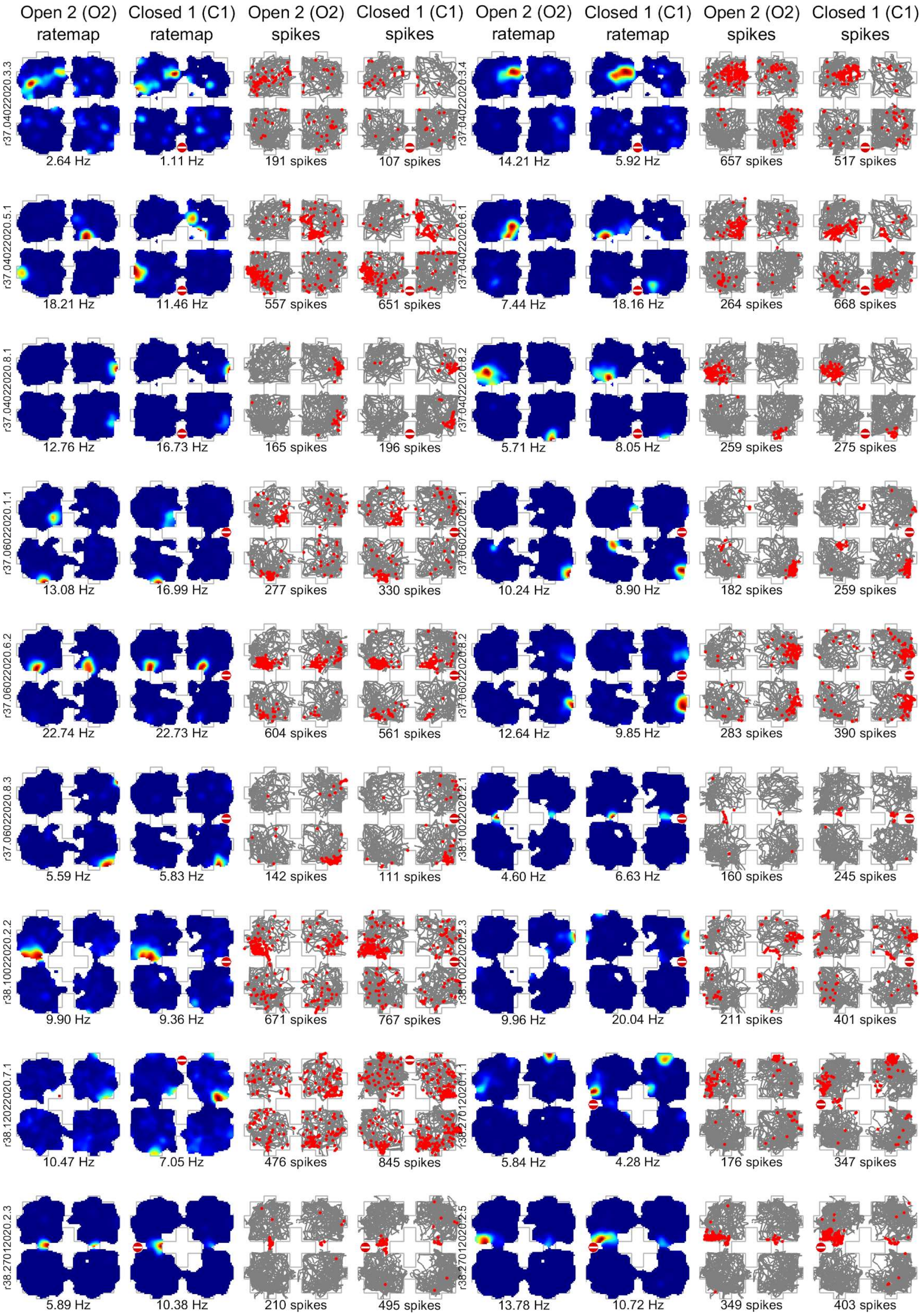

Open 2 (O2) ratemap
Closed 1 (C1) ratemap
Open 2 (O2) spikes
Closed 1 (C1) spikes
Open 2 (O2) ratemap
Closed 1 (C1) ratemap
Open 2 (O2) spikes
Closed 1 (C1) spikes
r37.04022020.3.3
2.64 Hz
1.11 Hz
191 spikes
107 spikes
r37.04022020.3.4
14.21 Hz
5.92 Hz
657 spikes
517 spikes
r37.04022020.5.1
18.21 Hz
11.46 Hz
557 spikes
651 spikes
r37.04022020.6.1
7.44 Hz
18.16 Hz
264 spikes
668 spikes
r37.04022020.8.1
12.76 Hz
16.73 Hz
165 spikes
196 spikes
r37.04022020.8.2
5.71 Hz
8.05 Hz
259 spikes
275 spikes
r37.06022020.1.1
13.08 Hz
16.99 Hz
277 spikes
330 spikes
r37.06022020.2.1
10.24 Hz
8.90 Hz
182 spikes
259 spikes
r37.06022020.6.2
22.74 Hz
22.73 Hz
604 spikes
561 spikes
r37.06022020.8.2
12.64 Hz
9.85 Hz
283 spikes
390 spikes
r37.06022020.8.3
5.59 Hz
5.83 Hz
142 spikes
111 spikes
r38.10022020.2.1
4.60 Hz
6.63 Hz
160 spikes
245 spikes
r38.10022020.2.2
9.90 Hz
9.36 Hz
671 spikes
767 spikes
r38.10022020.2.3
9.96 Hz
20.04 Hz
211 spikes
401 spikes
r38.12022020.7.1
10.47 Hz
7.05 Hz
476 spikes
845 spikes
r38.27012020.1.1
5.84 Hz
4.28 Hz
176 spikes
347 spikes
r38.27012020.2.3
5.89 Hz
10.38 Hz
210 spikes
495 spikes
r38.27012020.2.5
13.78 Hz
10.72 Hz
349 spikes
403 spikes

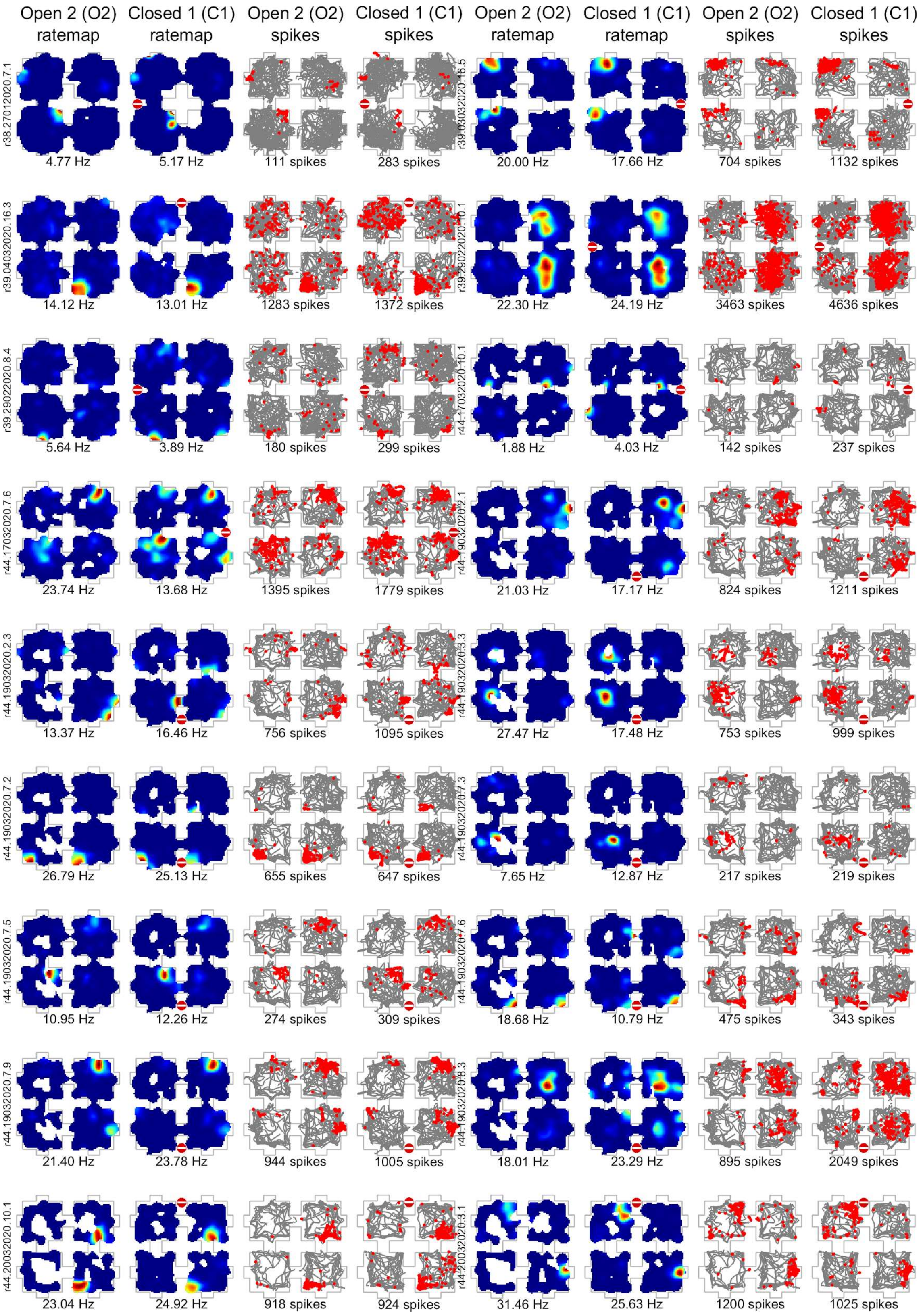
Open 2 (O2) ratemap
Closed 1 (C1) ratemap
Open 2 (O2) spikes
Closed 1 (C1) spikes
Open 2 (O2) ratemap
Closed 1 (C1) ratemap
Open 2 (O2) spikes
Closed 1 (C1) spikes
r38.27012020.7.1
4.77 Hz
5.17 Hz
111 spikes
283 spikes
r39.03032020.16.5
20.00 Hz
17.66 Hz
704 spikes
1132 spikes
r39.04032020.16.3
14.12 Hz
13.01 Hz
1283 spikes
1372 spikes
r39.29022020.10.1
22.30 Hz
24.19 Hz
3463 spikes
4636 spikes
r39.29022020.8.4
5.64 Hz
3.89 Hz
180 spikes
299 spikes
r44.17032020.10.1
1.88 Hz
4.03 Hz
142 spikes
237 spikes
r44.17032020.7.6
23.74 Hz
13.68 Hz
1395 spikes
1779 spikes
r44.19032020.2.1
21.03 Hz
17.17 Hz
824 spikes
1211 spikes
r44.19032020.2.3
13.37 Hz
16.46 Hz
756 spikes
1095 spikes
r44.19032020.3.3
27.47 Hz
17.48 Hz
753 spikes
999 spikes
r44.19032020.7.2
26.79 Hz
25.13 Hz
655 spikes
647 spikes
r44.19032020.7.3
7.65 Hz
12.87 Hz
217 spikes
219 spikes
r44.19032020.7.5
10.95 Hz
12.26 Hz
274 spikes
309 spikes
r44.19032020.7.6
18.68 Hz
10.79 Hz
475 spikes
343 spikes
r44.19032020.7.9
21.40 Hz
23.78 Hz
944 spikes
1005 spikes
r44.19032020.8.3
18.01 Hz
23.29 Hz
895 spikes
2049 spikes
r44.20032020.10.1
23.04 Hz
24.92 Hz
918 spikes
924 spikes
r44.20032020.3.1
31.46 Hz
25.63 Hz
1200 spikes
1025 spikes

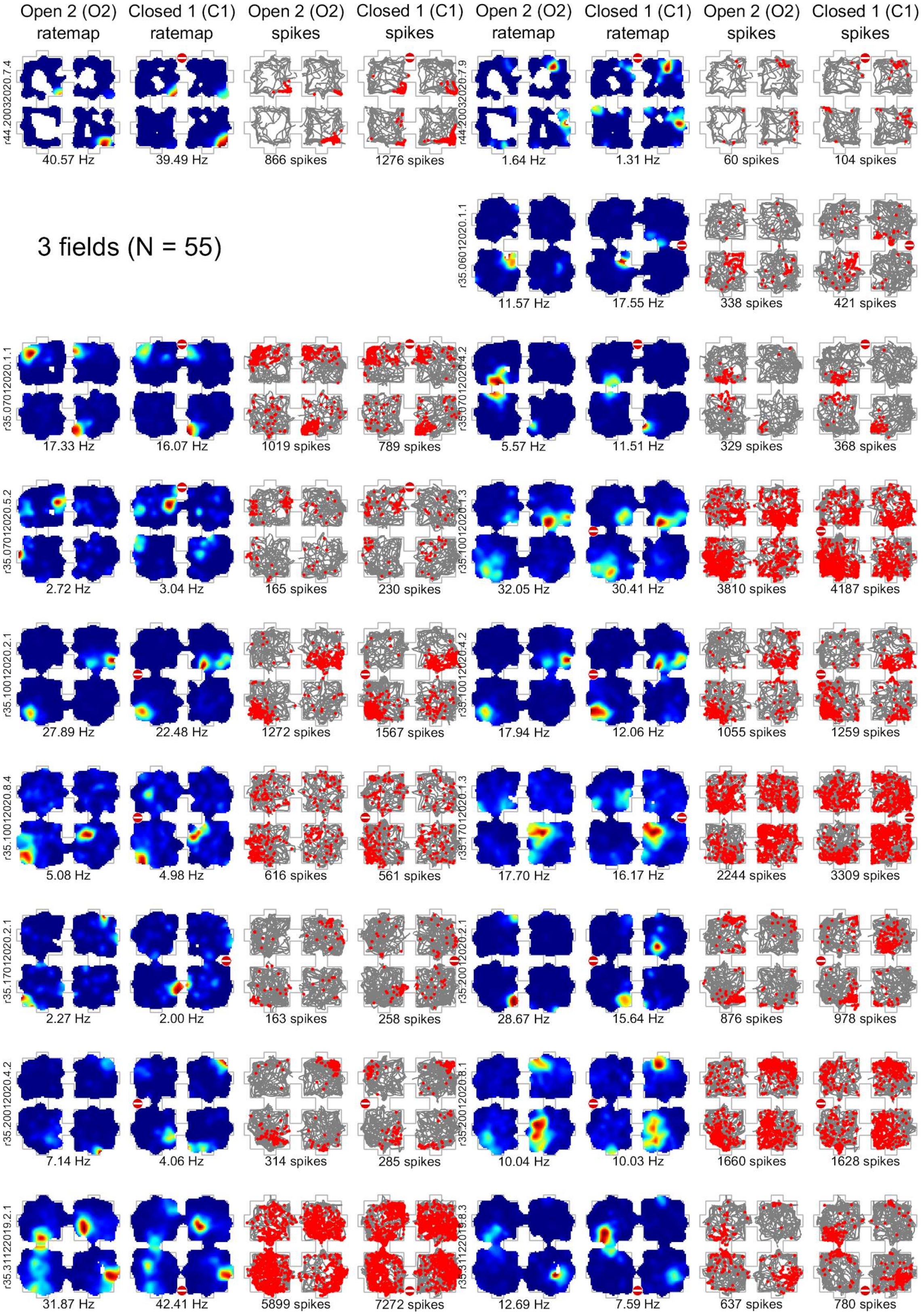
Open 2 (O2) ratemap
Closed 1 (C1) ratemap
Open 2 (O2) spikes
Closed 1 (C1) spikes
Open 2 (O2) ratemap
Closed 1 (C1) ratemap
Open 2 (O2) spikes
Closed 1 (C1) spikes
r44.20032020.7.4
40.57 Hz
39.49 Hz
866 spikes
1276 spikes
r44.20032020.7.9
1.64 Hz
1.31 Hz
60 spikes
104 spikes
3 fields (N = 55)
r35.06012020.1.1
11.57 Hz
17.55 Hz
338 spikes
421 spikes
r35.07012020.1.1
17.33 Hz
16.07 Hz
1019 spikes
789 spikes
r35.07012020.4.2
5.57 Hz
11.51 Hz
329 spikes
368 spikes
r35.07012020.5.2
2.72 Hz
3.04 Hz
165 spikes
230 spikes
r35.10012020.1.3
32.05 Hz
30.41 Hz
3810 spikes
4187 spikes
r35.10012020.2.1
27.89 Hz
22.48 Hz
1272 spikes
1567 spikes
r35.10012020.4.2
17.94 Hz
12.06 Hz
1055 spikes
1259 spikes
r35.10012020.8.4
5.08 Hz
4.98 Hz
616 spikes
561 spikes
r35.17012020.1.3
17.70 Hz
16.17 Hz
2244 spikes
3309 spikes
r35.17012020.2.1
2.27 Hz
2.00 Hz
163 spikes
258 spikes
r35.20012020.2.1
28.67 Hz
15.64 Hz
876 spikes
978 spikes
r35.20012020.4.2
7.14 Hz
4.06 Hz
314 spikes
285 spikes
r35.20012020.8.1
10.04 Hz
10.03 Hz
1660 spikes
1628 spikes
r35.31122019.2.1
31.87 Hz
42.41 Hz
5899 spikes
7272 spikes
r35.31122019.8.3
12.69 Hz
7.59 Hz
637 spikes
780 spikes

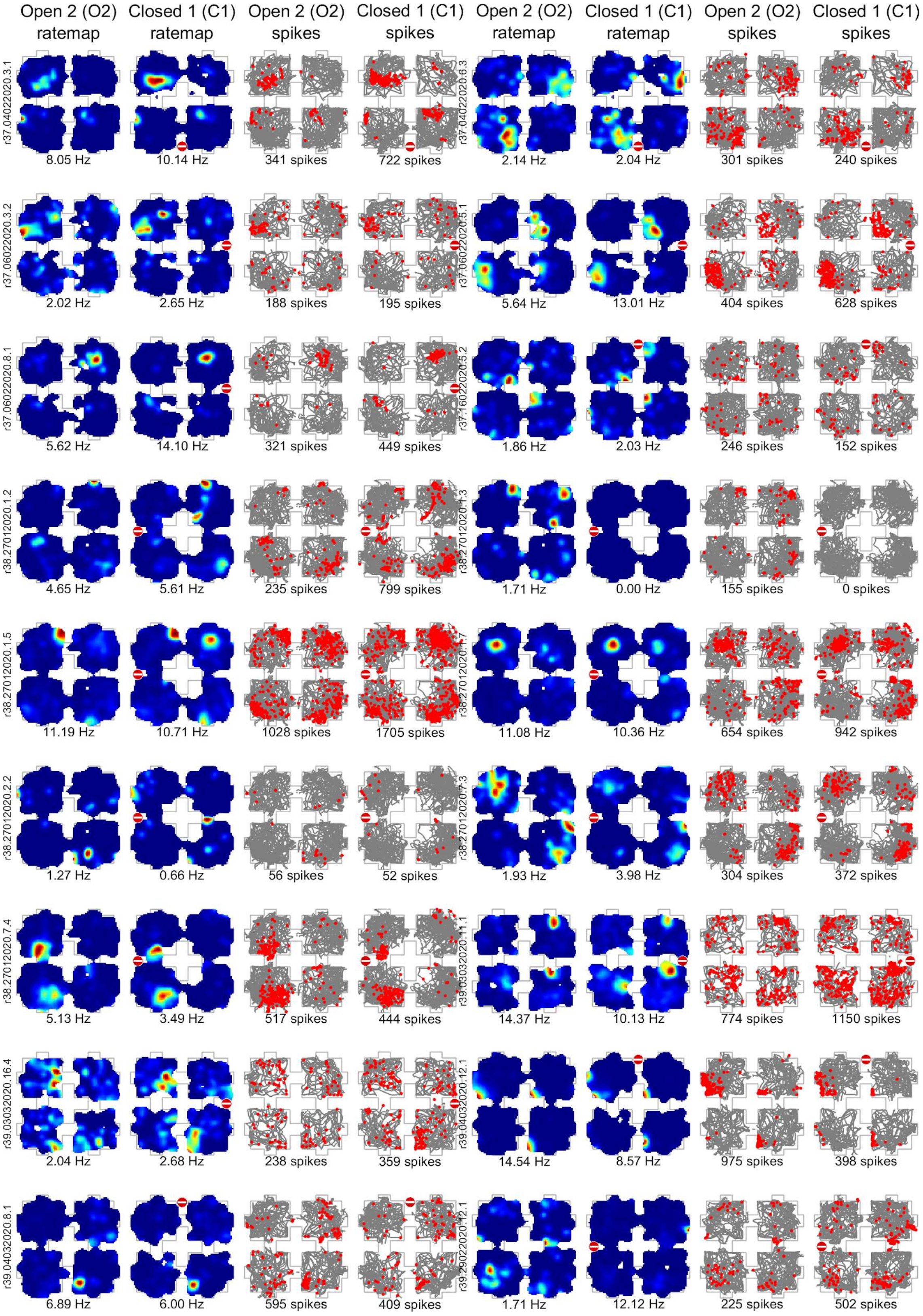
Open 2 (O2) ratemap
Closed 1 (C1) ratemap
Open 2 (O2) spikes
Closed 1 (C1) spikes
Open 2 (O2) ratemap
Closed 1 (C1) ratemap
Open 2 (O2) spikes
Closed 1 (C1) spikes
r37.04022020.3.1
8.05 Hz
10.14 Hz
341 spikes
722 spikes
r37.04022020.6.3
2.14 Hz
2.04 Hz
301 spikes
240 spikes
r37.06022020.3.2
2.02 Hz
2.65 Hz
188 spikes
195 spikes
5.64 Hz
13.01 Hz
404 spikes
628 spikes
r37.06022020.8.1
5.62 Hz
14.10 Hz
321 spikes
449 spikes
1.86 Hz
2.03 Hz
246 spikes
152 spikes
r38.27012020.1.2
4.65 Hz
5.61 Hz
235 spikes
799 spikes
1.71 Hz
0.00 Hz
155 spikes
0 spikes
r38.27012020.1.5
11.19 Hz
10.71 Hz
1028 spikes
1705 spikes
11.08 Hz
10.36 Hz
654 spikes
942 spikes
r38.27012020.2.2
1.27 Hz
0.66 Hz
56 spikes
52 spikes
1.93 Hz
3.98 Hz
304 spikes
372 spikes
r38.27012020.7.4
5.13 Hz
3.49 Hz
517 spikes
444 spikes
14.37 Hz
10.13 Hz
774 spikes
1150 spikes
r39.03032020.16.4
2.04 Hz
2.68 Hz
238 spikes
359 spikes
14.54 Hz
8.57 Hz
975 spikes
398 spikes
r39.04032020.8.1
6.89 Hz
6.00 Hz
595 spikes
409 spikes
1.71 Hz
12.12 Hz
225 spikes
502 spikes

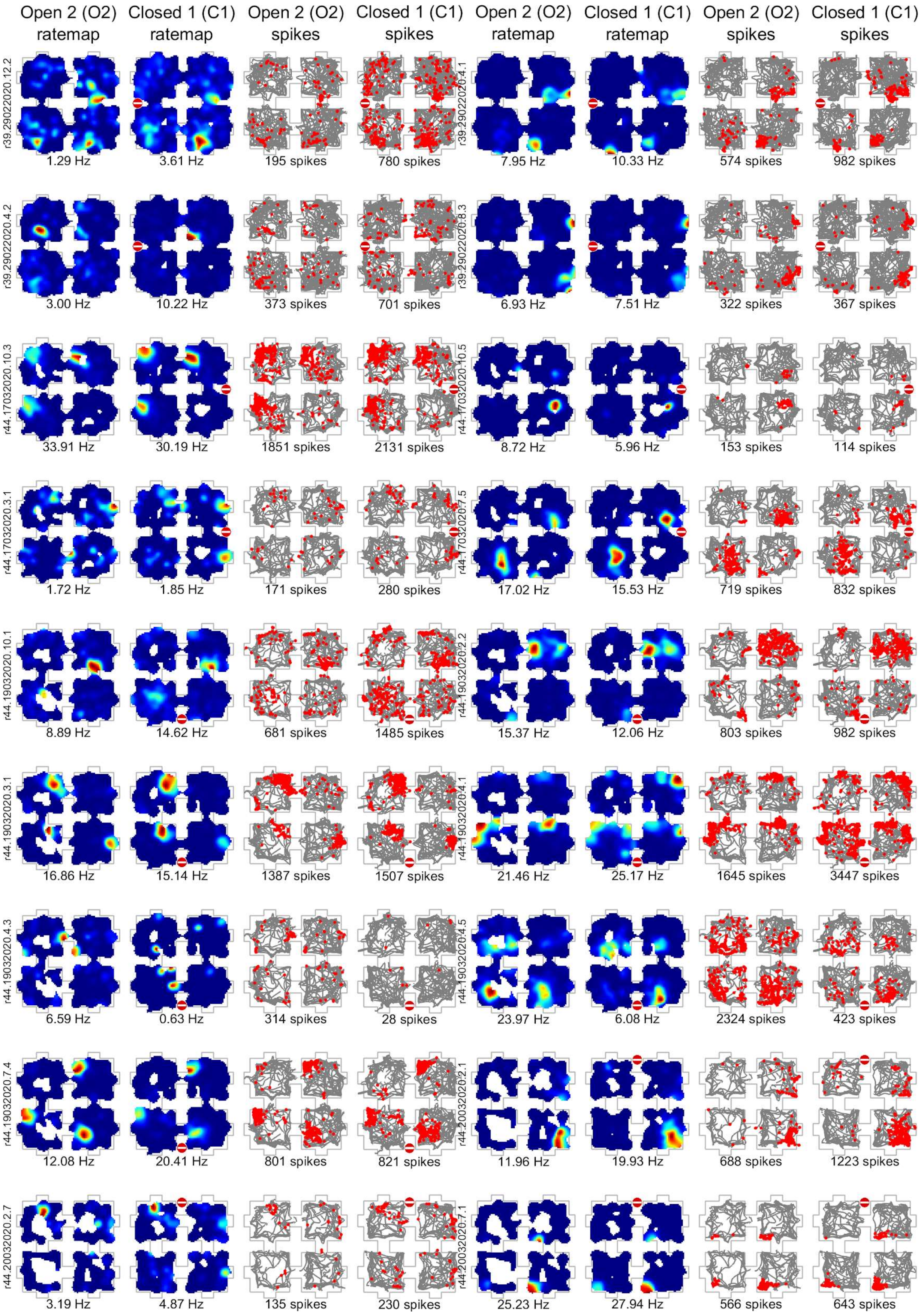
Open 2 (O2) ratemap
Closed 1 (C1) ratemap
Open 2 (O2) spikes
Closed 1 (C1) spikes
Open 2 (O2) ratemap
Closed 1 (C1) ratemap
Open 2 (O2) spikes
Closed 1 (C1) spikes
r39.29022020.12.2
1.29 Hz
3.61 Hz
195 spikes
780 spikes
r39.29022020.4.1
7.95 Hz
10.33 Hz
574 spikes
982 spikes
r39.29022020.4.2
3.00 Hz
10.22 Hz
373 spikes
701 spikes
r39.29022020.8.3
6.93 Hz
7.51 Hz
322 spikes
367 spikes
r44.17032020.10.3
33.91 Hz
30.19 Hz
1851 spikes
2131 spikes
r44.17032020.10.5
8.72 Hz
5.96 Hz
153 spikes
114 spikes
r44.17032020.3.1
1.72 Hz
1.85 Hz
171 spikes
280 spikes
r44.17032020.7.5
17.02 Hz
15.53 Hz
719 spikes
832 spikes
r44.19032020.10.1
8.89 Hz
14.62 Hz
681 spikes
1485 spikes
r44.19032020.2.2
15.37 Hz
12.06 Hz
803 spikes
982 spikes
r44.19032020.3.1
16.86 Hz
15.14 Hz
1387 spikes
1507 spikes
r44.19032020.4.1
21.46 Hz
25.17 Hz
1645 spikes
3447 spikes
r44.19032020.4.3
6.59 Hz
0.63 Hz
314 spikes
28 spikes
r44.19032020.4.5
23.97 Hz
6.08 Hz
2324 spikes
423 spikes
r44.19032020.7.4
12.08 Hz
20.41 Hz
801 spikes
821 spikes
r44.20032020.2.1
11.96 Hz
19.93 Hz
688 spikes
1223 spikes
r44.20032020.2.7
3.19 Hz
4.87 Hz
135 spikes
230 spikes
r44.20032020.7.1
25.23 Hz
27.94 Hz
566 spikes
643 spikes

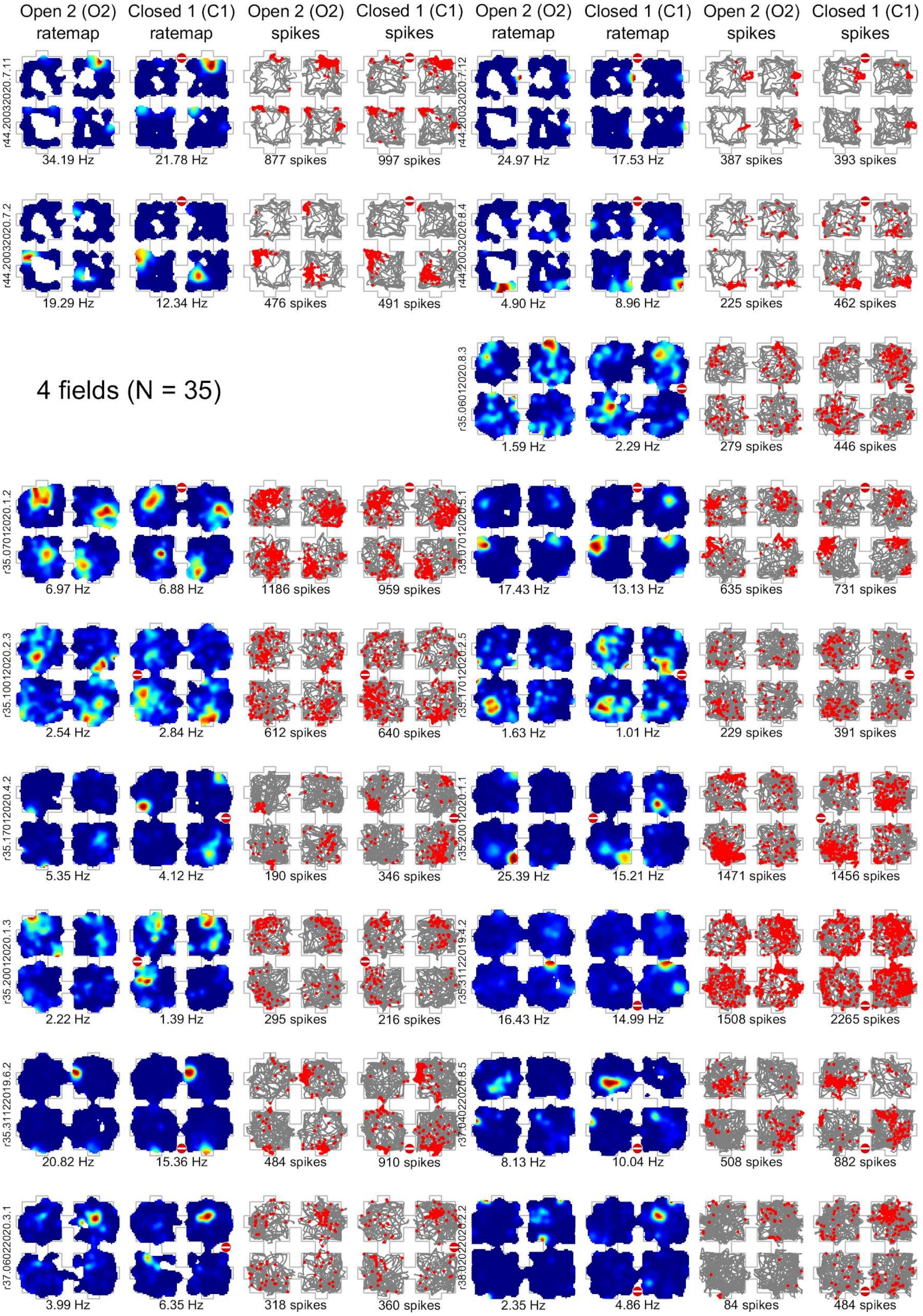

Open 2 (O2) ratemap
Closed 1 (C1) ratemap
Open 2 (O2) spikes
Closed 1 (C1) spikes
Open 2 (O2) ratemap
Closed 1 (C1) ratemap
Open 2 (O2) spikes
Closed 1 (C1) spikes
r44.20032020.7.11
34.19 Hz
21.78 Hz
877 spikes
997 spikes
r44.20032020.7.12
24.97 Hz
17.53 Hz
387 spikes
393 spikes
r44.20032020.7.2
19.29 Hz
12.34 Hz
476 spikes
491 spikes
4.90 Hz
8.96 Hz
225 spikes
462 spikes
4 fields (N = 35)
r35.06012020.8.3
1.59 Hz
2.29 Hz
279 spikes
446 spikes
r35.07012020.1.2
6.97 Hz
6.88 Hz
1186 spikes
959 spikes
17.43 Hz
13.13 Hz
635 spikes
731 spikes
r35.10012020.2.3
2.54 Hz
2.84 Hz
612 spikes
640 spikes
1.63 Hz
1.01 Hz
229 spikes
391 spikes
r35.17012020.4.2
5.35 Hz
4.12 Hz
190 spikes
346 spikes
25.39 Hz
15.21 Hz
1471 spikes
1456 spikes
r35.20012020.1.3
2.22 Hz
1.39 Hz
295 spikes
216 spikes
16.43 Hz
14.99 Hz
1508 spikes
2265 spikes
r35.31122019.6.2
20.82 Hz
15.36 Hz
484 spikes
910 spikes
8.13 Hz
10.04 Hz
508 spikes
882 spikes
r37.06022020.3.1
3.99 Hz
6.35 Hz
318 spikes
360 spikes
2.35 Hz
4.86 Hz
84 spikes
484 spikes

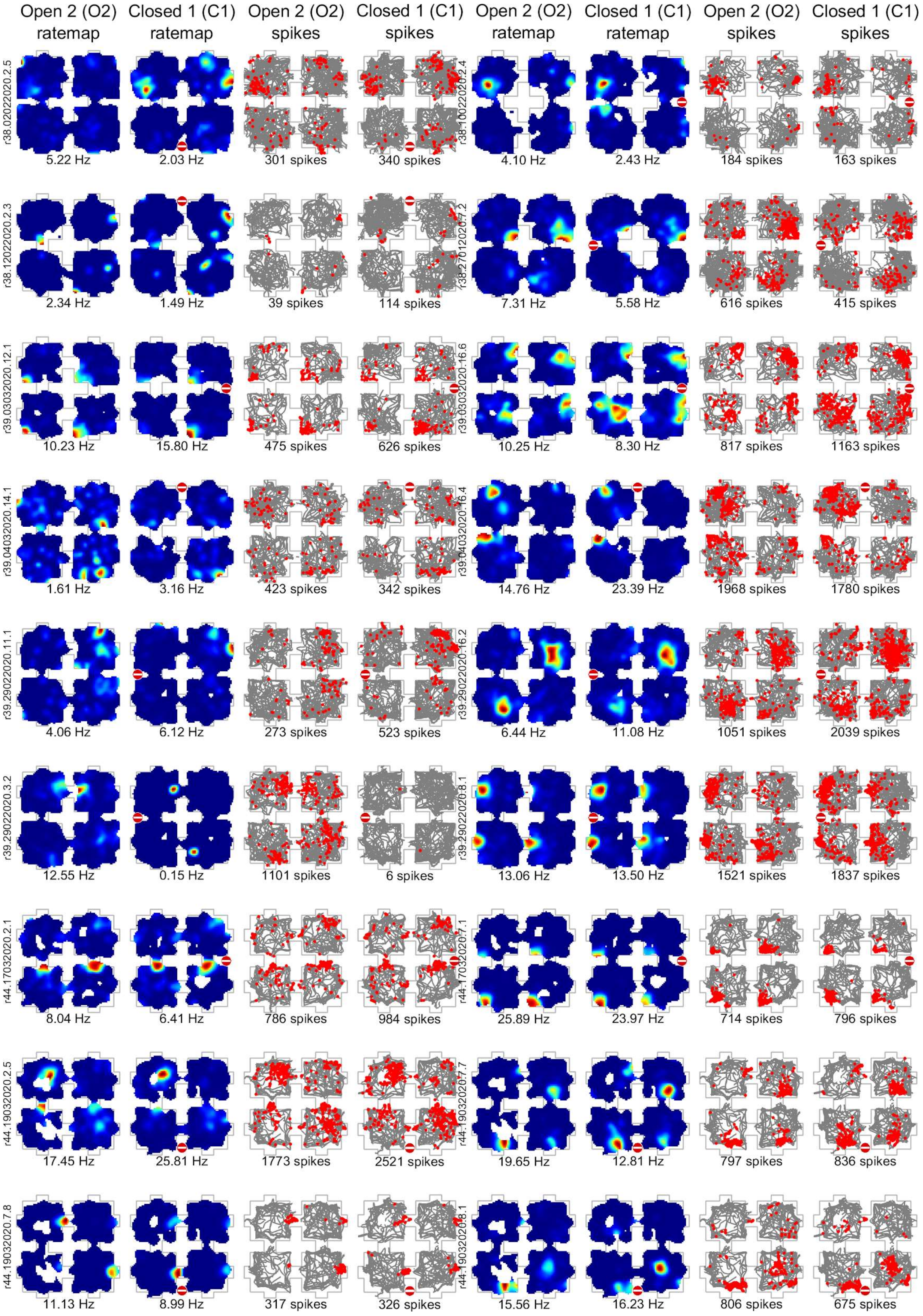
Open 2 (O2) ratemap
Closed 1 (C1) ratemap
Open 2 (O2) spikes
Closed 1 (C1) spikes
Open 2 (O2) ratemap
Closed 1 (C1) ratemap
Open 2 (O2) spikes
Closed 1 (C1) spikes
r38.02022020.2.5
5.22 Hz
2.03 Hz
301 spikes
340 spikes
4.10 Hz
2.43 Hz
184 spikes
163 spikes
r38.12022020.2.3
2.34 Hz
1.49 Hz
39 spikes
114 spikes
7.31 Hz
5.58 Hz
616 spikes
415 spikes
r39.03032020.12.1
10.23 Hz
15.80 Hz
475 spikes
626 spikes
10.25 Hz
8.30 Hz
817 spikes
1163 spikes
r39.04032020.14.1
1.61 Hz
3.16 Hz
423 spikes
342 spikes
14.76 Hz
23.39 Hz
1968 spikes
1780 spikes
r39.29022020.11.1
4.06 Hz
6.12 Hz
273 spikes
523 spikes
6.44 Hz
11.08 Hz
1051 spikes
2039 spikes
r39.29022020.3.2
12.55 Hz
0.15 Hz
1101 spikes
6 spikes
13.06 Hz
13.50 Hz
1521 spikes
1837 spikes
r44.17032020.2.1
8.04 Hz
6.41 Hz
786 spikes
984 spikes
25.89 Hz
23.97 Hz
714 spikes
796 spikes
r44.19032020.2.5
17.45 Hz
25.81 Hz
1773 spikes
2521 spikes
19.65 Hz
12.81 Hz
797 spikes
836 spikes
r44.19032020.7.8
11.13 Hz
8.99 Hz
317 spikes
326 spikes
15.56 Hz
16.23 Hz
806 spikes
675 spikes

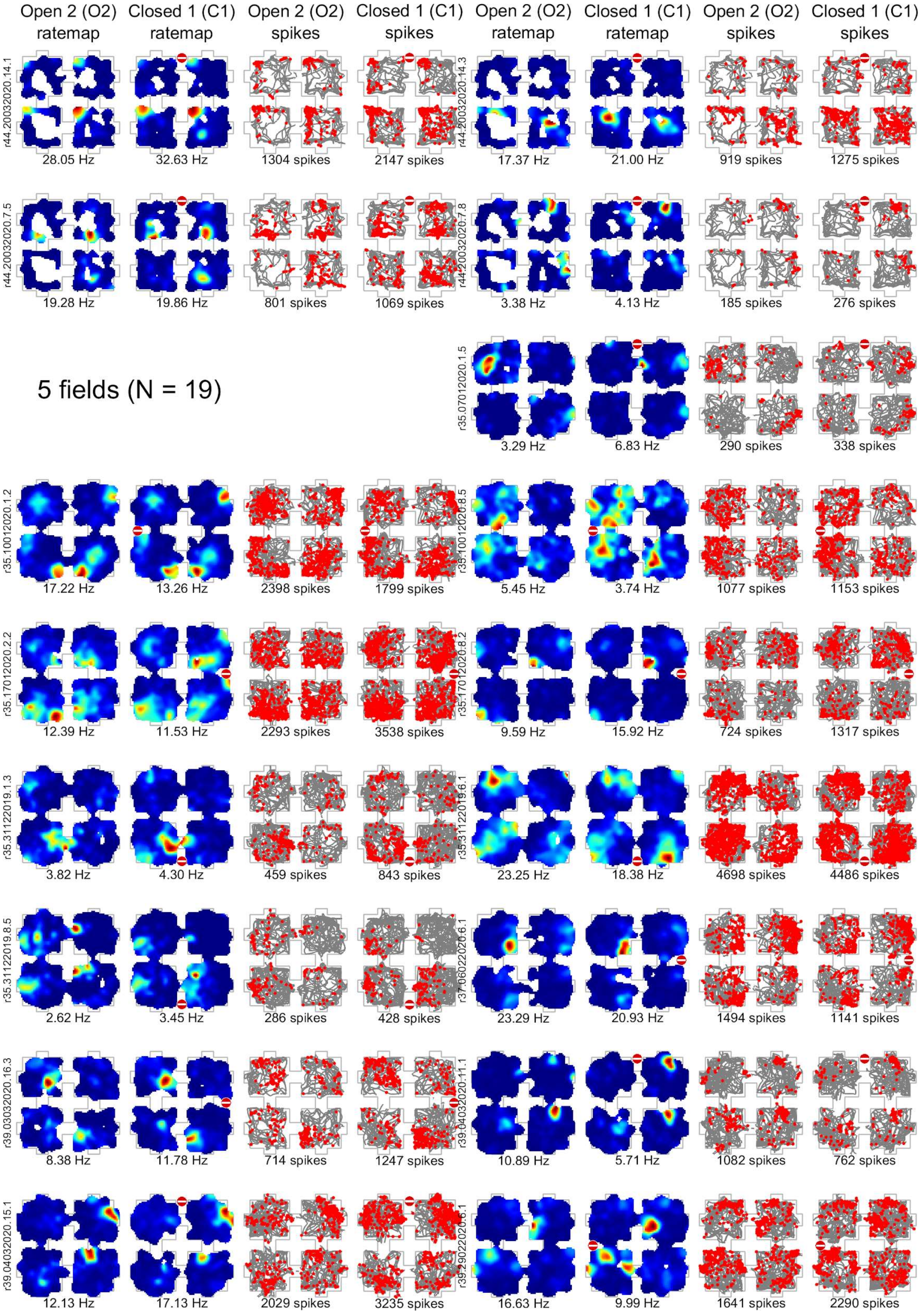
Open 2 (O2) ratemap
Closed 1 (C1) ratemap
Open 2 (O2) spikes
Closed 1 (C1) spikes
Open 2 (O2) ratemap
Closed 1 (C1) ratemap
Open 2 (O2) spikes
Closed 1 (C1) spikes
r44.20032020.14.1
28.05 Hz
32.63 Hz
1304 spikes
2147 spikes
r44.20032020.14.3
17.37 Hz
21.00 Hz
919 spikes
1275 spikes
r44.20032020.7.5
19.28 Hz
19.86 Hz
801 spikes
1069 spikes
3.38 Hz
4.13 Hz
185 spikes
276 spikes
5 fields (N = 19)
r35.07012020.1.5
3.29 Hz
6.83 Hz
290 spikes
338 spikes
r35.10012020.1.2
17.22 Hz
13.26 Hz
2398 spikes
1799 spikes
5.45 Hz
3.74 Hz
1077 spikes
1153 spikes
r35.17012020.2.2
12.39 Hz
11.53 Hz
2293 spikes
3538 spikes
9.59 Hz
15.92 Hz
724 spikes
1317 spikes
r35.31122019.1.3
3.82 Hz
4.30 Hz
459 spikes
843 spikes
23.25 Hz
18.38 Hz
4698 spikes
4486 spikes
r35.31122019.8.5
2.62 Hz
3.45 Hz
286 spikes
428 spikes
23.29 Hz
20.93 Hz
1494 spikes
1141 spikes
r39.03032020.16.3
8.38 Hz
11.78 Hz
714 spikes
1247 spikes
10.89 Hz
5.71 Hz
1082 spikes
762 spikes
r39.04032020.15.1
12.13 Hz
17.13 Hz
2029 spikes
3235 spikes
16.63 Hz
9.99 Hz
1641 spikes
2290 spikes

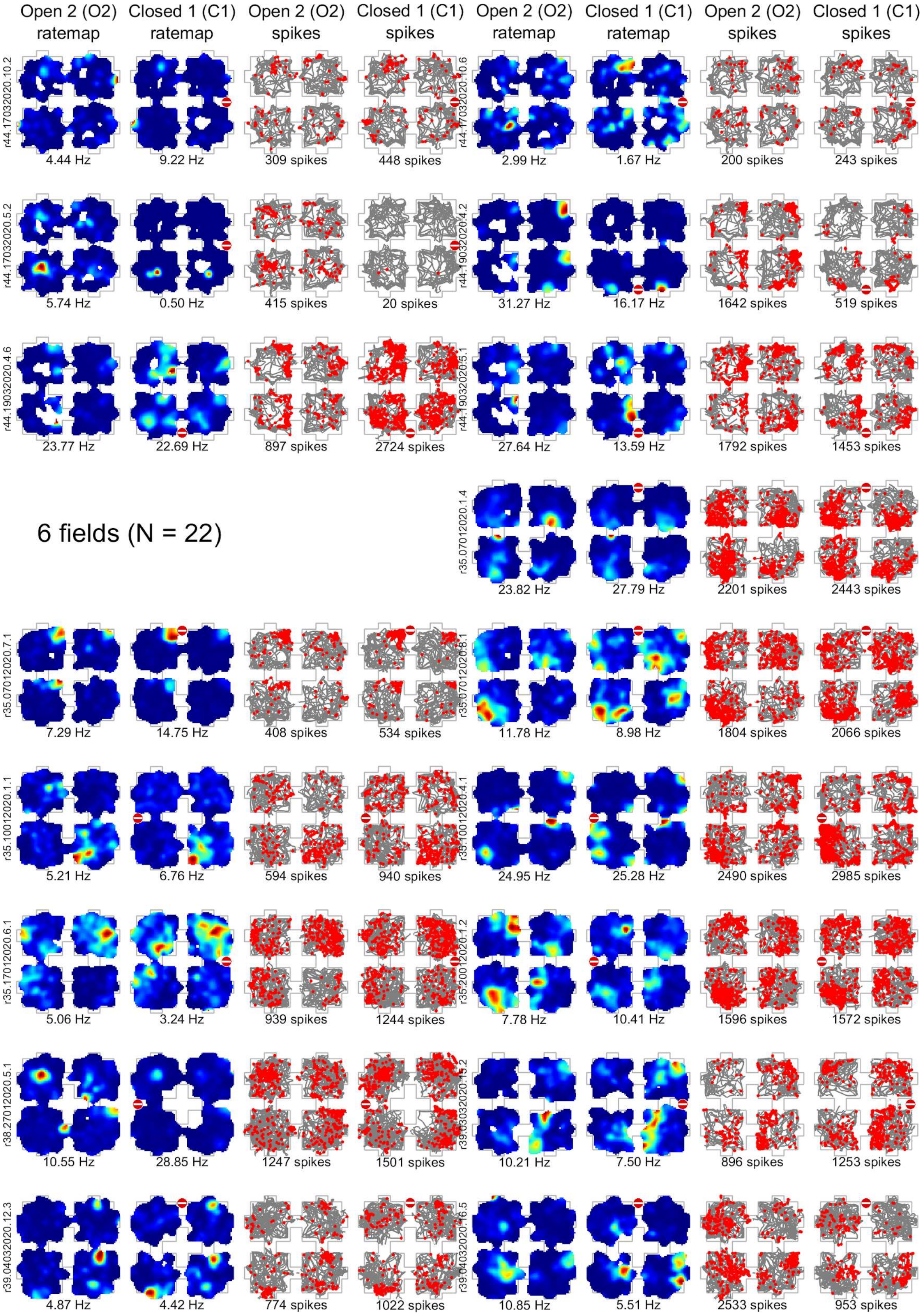

Open 2 (O2) ratemap
Closed 1 (C1) ratemap
Open 2 (O2) spikes
Closed 1 (C1) spikes
Open 2 (O2) ratemap
Closed 1 (C1) ratemap
Open 2 (O2) spikes
Closed 1 (C1) spikes
r44.17032020.10.2
4.44 Hz
9.22 Hz
309 spikes
448 spikes
2.99 Hz
1.67 Hz
200 spikes
243 spikes
r44.17032020.5.2
5.74 Hz
0.50 Hz
415 spikes
20 spikes
31.27 Hz
16.17 Hz
1642 spikes
519 spikes
r44.19032020.4.6
23.77 Hz
22.69 Hz
897 spikes
2724 spikes
27.64 Hz
13.59 Hz
1792 spikes
1453 spikes
6 fields (N = 22)
r35.07012020.1.4
23.82 Hz
27.79 Hz
2201 spikes
2443 spikes
r35.07012020.7.1
7.29 Hz
14.75 Hz
408 spikes
534 spikes
11.78 Hz
8.98 Hz
1804 spikes
2066 spikes
r35.10012020.1.1
5.21 Hz
6.76 Hz
594 spikes
940 spikes
24.95 Hz
25.28 Hz
2490 spikes
2985 spikes
r35.17012020.6.1
5.06 Hz
3.24 Hz
939 spikes
1244 spikes
7.78 Hz
10.41 Hz
1596 spikes
1572 spikes
r38.27012020.5.1
10.55 Hz
28.85 Hz
1247 spikes
1501 spikes
10.21 Hz
7.50 Hz
896 spikes
1253 spikes
r39.04032020.12.3
4.87 Hz
4.42 Hz
774 spikes
1022 spikes
10.85 Hz
5.51 Hz
2533 spikes
953 spikes

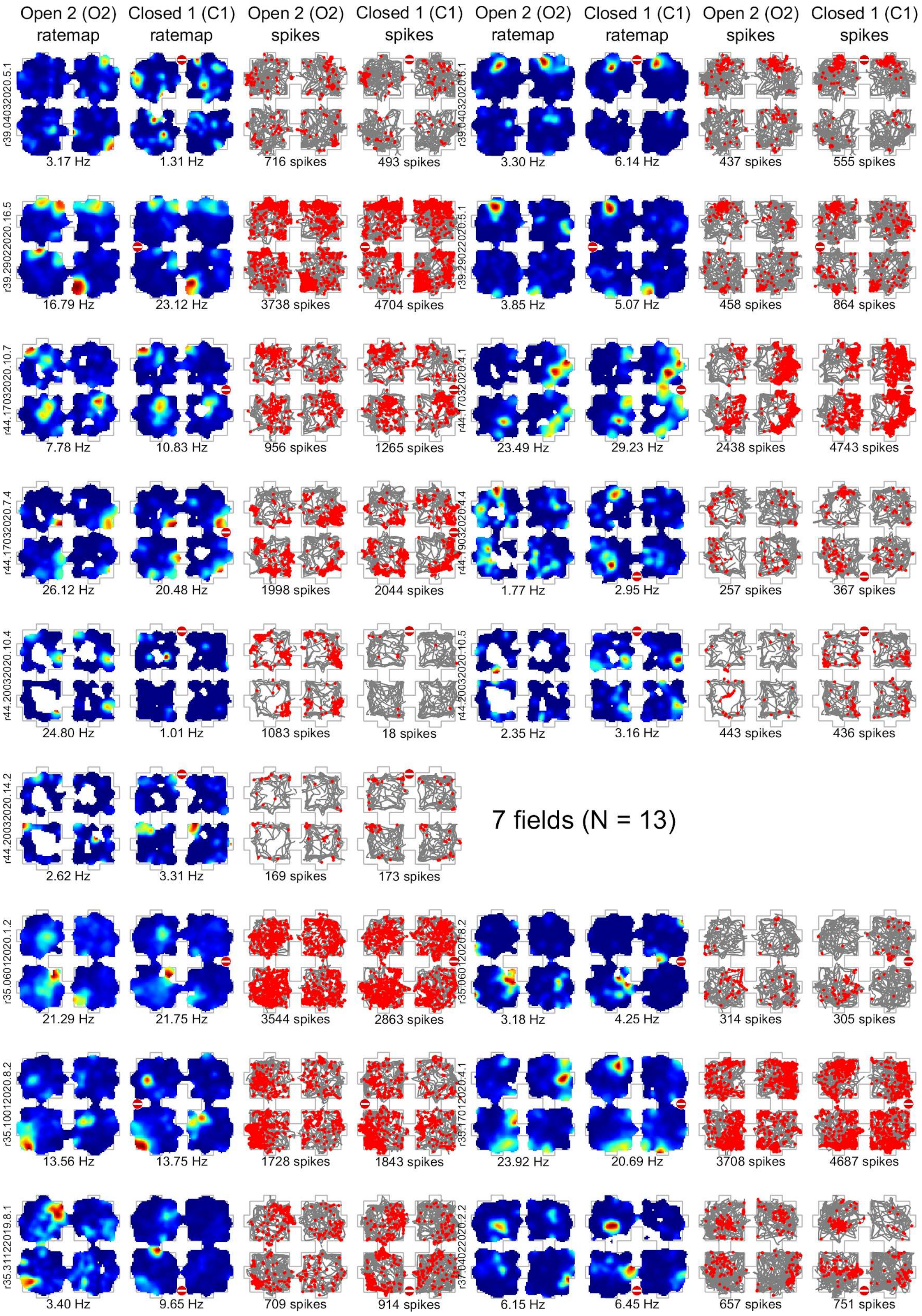
Open 2 (O2) ratemap
Closed 1 (C1) ratemap
Open 2 (O2) spikes
Closed 1 (C1) spikes
Open 2 (O2) ratemap
Closed 1 (C1) ratemap
Open 2 (O2) spikes
Closed 1 (C1) spikes
r39.04032020.5.1
3.17 Hz
1.31 Hz
716 spikes
493 spikes
r39.04032020.6.1
3.30 Hz
6.14 Hz
437 spikes
555 spikes
r39.29022020.16.5
16.79 Hz
23.12 Hz
3738 spikes
4704 spikes
r39.29022020.5.1
3.85 Hz
5.07 Hz
458 spikes
864 spikes
r44.17032020.10.7
7.78 Hz
10.83 Hz
956 spikes
1265 spikes
r44.17032020.4.1
23.49 Hz
29.23 Hz
2438 spikes
4743 spikes
r44.17032020.7.4
26.12 Hz
20.48 Hz
1998 spikes
2044 spikes
r44.19032020.4.4
1.77 Hz
2.95 Hz
257 spikes
367 spikes
r44.20032020.10.4
24.80 Hz
1.01 Hz
1083 spikes
18 spikes
r44.20032020.10.5
2.35 Hz
3.16 Hz
443 spikes
436 spikes
r44.20032020.14.2
2.62 Hz
3.31 Hz
169 spikes
173 spikes
7 fields (N = 13)
r35.06012020.1.2
21.29 Hz
21.75 Hz
3544 spikes
2863 spikes
r35.06012020.8.2
3.18 Hz
4.25 Hz
314 spikes
305 spikes
r35.10012020.8.2
13.56 Hz
13.75 Hz
1728 spikes
1843 spikes
r35.17012020.4.1
23.92 Hz
20.69 Hz
3708 spikes
4687 spikes
r35.31122019.8.1
3.40 Hz
9.65 Hz
709 spikes
914 spikes
r37.04022020.2.2
6.15 Hz
6.45 Hz
657 spikes
751 spikes

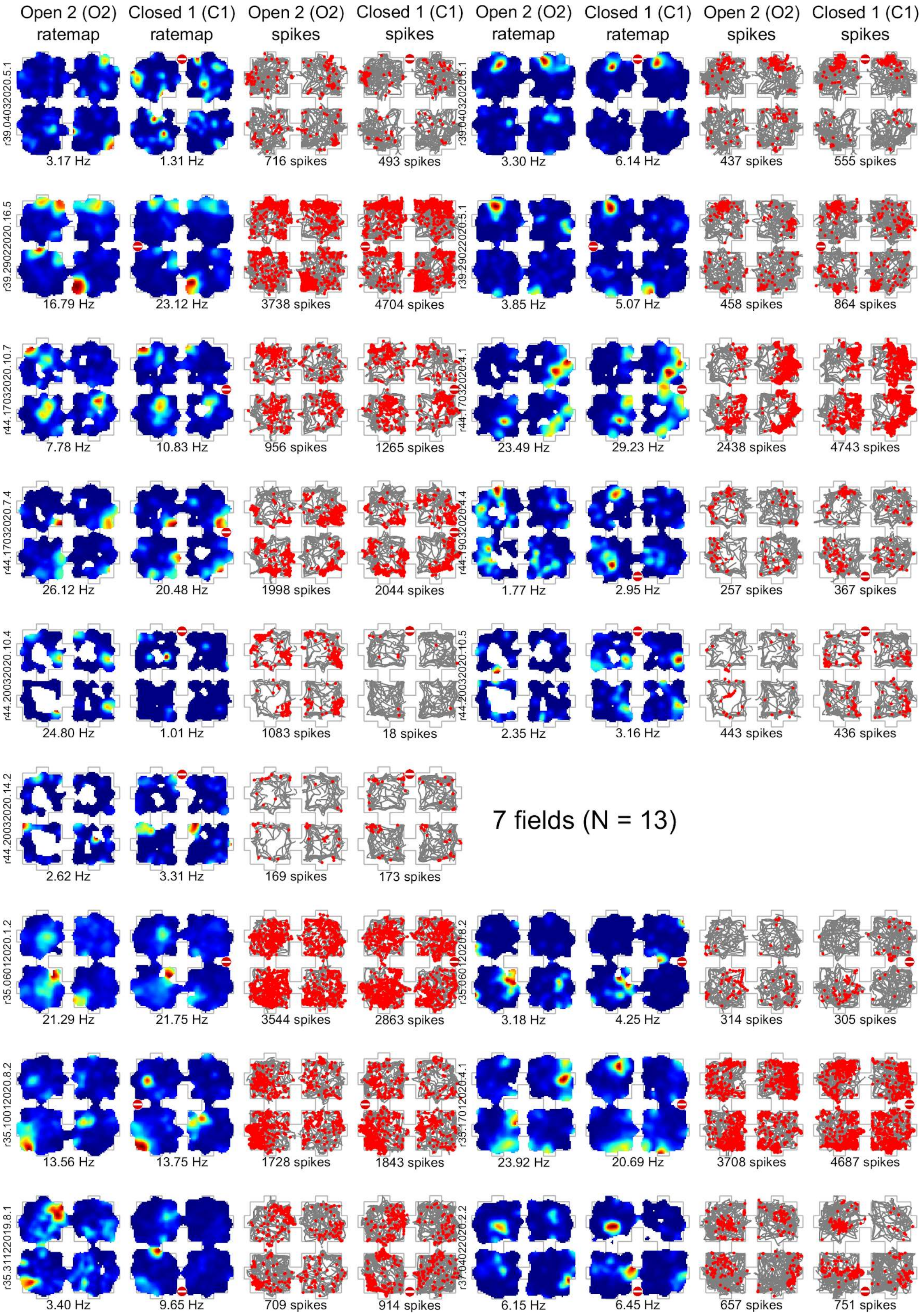
Open 2 (O2) ratemap
Closed 1 (C1) ratemap
Open 2 (O2) spikes
Closed 1 (C1) spikes
Open 2 (O2) ratemap
Closed 1 (C1) ratemap
Open 2 (O2) spikes
Closed 1 (C1) spikes
r39.04032020.5.1
3.17 Hz
1.31 Hz
716 spikes
493 spikes
r39.04032020.6.1
3.30 Hz
6.14 Hz
437 spikes
555 spikes
r39.29022020.16.5
16.79 Hz
23.12 Hz
3738 spikes
4704 spikes
r39.29022020.5.1
3.85 Hz
5.07 Hz
458 spikes
864 spikes
r44.17032020.10.7
7.78 Hz
10.83 Hz
956 spikes
1265 spikes
r44.17032020.4.1
23.49 Hz
29.23 Hz
2438 spikes
4743 spikes
r44.17032020.7.4
26.12 Hz
20.48 Hz
1998 spikes
2044 spikes
r44.19032020.4.4
1.77 Hz
2.95 Hz
257 spikes
367 spikes
r44.20032020.10.4
24.80 Hz
1.01 Hz
1083 spikes
18 spikes
r44.20032020.10.5
2.35 Hz
3.16 Hz
443 spikes
436 spikes
r44.20032020.14.2
2.62 Hz
3.31 Hz
169 spikes
173 spikes
7 fields (N = 13)
r35.06012020.1.2
21.29 Hz
21.75 Hz
3544 spikes
2863 spikes
r35.06012020.8.2
3.18 Hz
4.25 Hz
314 spikes
305 spikes
r35.10012020.8.2
13.56 Hz
13.75 Hz
1728 spikes
1843 spikes
r35.17012020.4.1
23.92 Hz
20.69 Hz
3708 spikes
4687 spikes
r35.31122019.8.1
3.40 Hz
9.65 Hz
709 spikes
914 spikes
r37.04022020.2.2
6.15 Hz
6.45 Hz
657 spikes
751 spikes

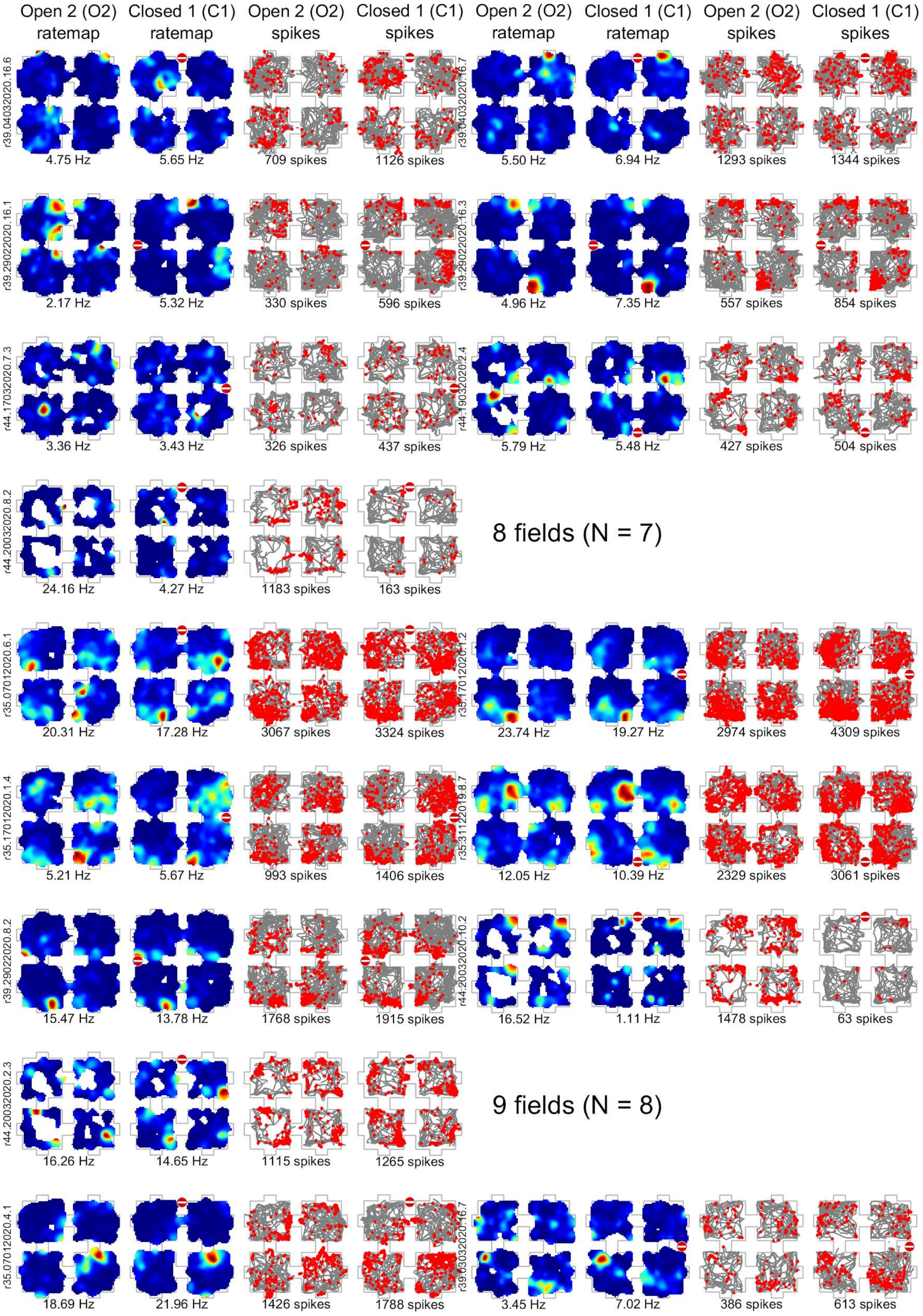
Open 2 (O2) ratemap
Closed 1 (C1) ratemap
Open 2 (O2) spikes
Closed 1 (C1) spikes
Open 2 (O2) ratemap
Closed 1 (C1) ratemap
Open 2 (O2) spikes
Closed 1 (C1) spikes
r39.04032020.16.6
4.75 Hz
5.65 Hz
709 spikes
1126 spikes
r39.04032020.16.7
5.50 Hz
6.94 Hz
1293 spikes
1344 spikes
r39.29022020.16.1
2.17 Hz
5.32 Hz
330 spikes
596 spikes
r39.29022020.16.3
4.96 Hz
7.35 Hz
557 spikes
854 spikes
r44.17032020.7.3
3.36 Hz
3.43 Hz
326 spikes
437 spikes
r44.19032020.2.4
5.79 Hz
5.48 Hz
427 spikes
504 spikes
r44.20032020.8.2
24.16 Hz
4.27 Hz
1183 spikes
163 spikes
8 fields (N = 7)
r35.07012020.6.1
20.31 Hz
17.28 Hz
3067 spikes
3324 spikes
r35.17012020.1.2
23.74 Hz
19.27 Hz
2974 spikes
4309 spikes
r35.17012020.1.4
5.21 Hz
5.67 Hz
993 spikes
1406 spikes
r35.31122019.8.7
12.05 Hz
10.39 Hz
2329 spikes
3061 spikes
r39.29022020.8.2
15.47 Hz
13.78 Hz
1768 spikes
1915 spikes
r44.20032020.10.2
16.52 Hz
1.11 Hz
1478 spikes
63 spikes
r44.20032020.2.3
16.26 Hz
14.65 Hz
1115 spikes
1265 spikes
9 fields (N = 8)
r35.07012020.4.1
18.69 Hz
21.96 Hz
1426 spikes
1788 spikes
r39.03032020.16.7
3.45 Hz
7.02 Hz
386 spikes
613 spikes

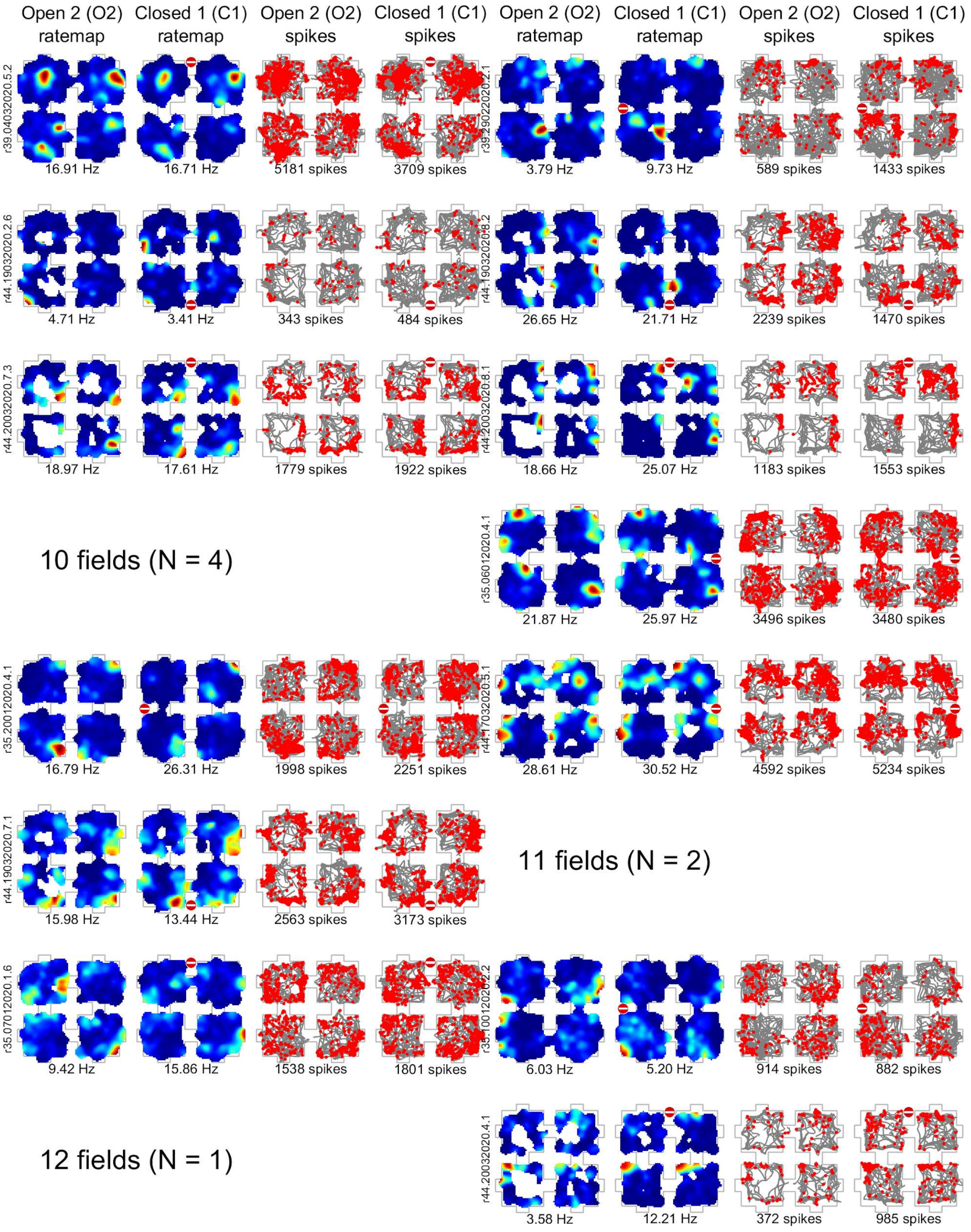
Open 2 (O2) ratemap
Closed 1 (C1) ratemap
Open 2 (O2) spikes
Closed 1 (C1) spikes
r39.04032020.5.2
16.91 Hz
16.71 Hz
5181 spikes
3709 spikes
3.79 Hz
9.73 Hz
589 spikes
1433 spikes
r44.19032020.2.6
4.71 Hz
3.41 Hz
343 spikes
484 spikes
26.65 Hz
21.71 Hz
2239 spikes
1470 spikes
r44.20032020.7.3
18.97 Hz
17.61 Hz
1779 spikes
1922 spikes
18.66 Hz
25.07 Hz
1183 spikes
1553 spikes
10 fields (N = 4)
r35.06012020.4.1
21.87 Hz
25.97 Hz
3496 spikes
3480 spikes
r35.20012020.4.1
16.79 Hz
26.31 Hz
1998 spikes
2251 spikes
28.61 Hz
30.52 Hz
4592 spikes
5234 spikes
r44.19032020.7.1
15.98 Hz
13.44 Hz
2563 spikes
3173 spikes
11 fields (N = 2)
r35.07012020.1.6
9.42 Hz
15.86 Hz
1538 spikes
1801 spikes
6.03 Hz
5.20 Hz
914 spikes
882 spikes
12 fields (N = 1)
r44.20032020.4.1
3.58 Hz
12.21 Hz
372 spikes
985 spikes

**Supplementary Figure 11**: Individual place cell plots for O2 and C1 in One-Way

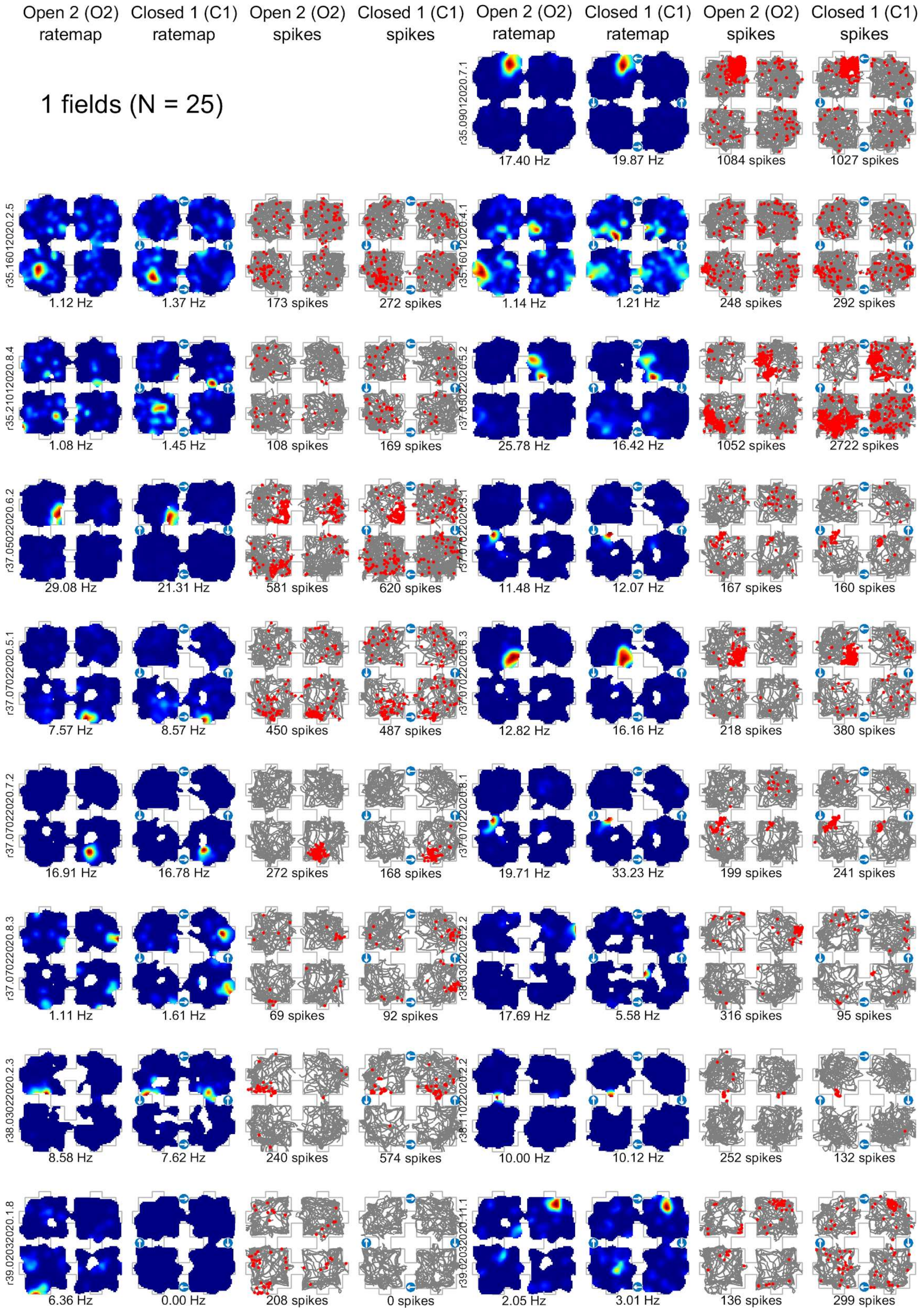

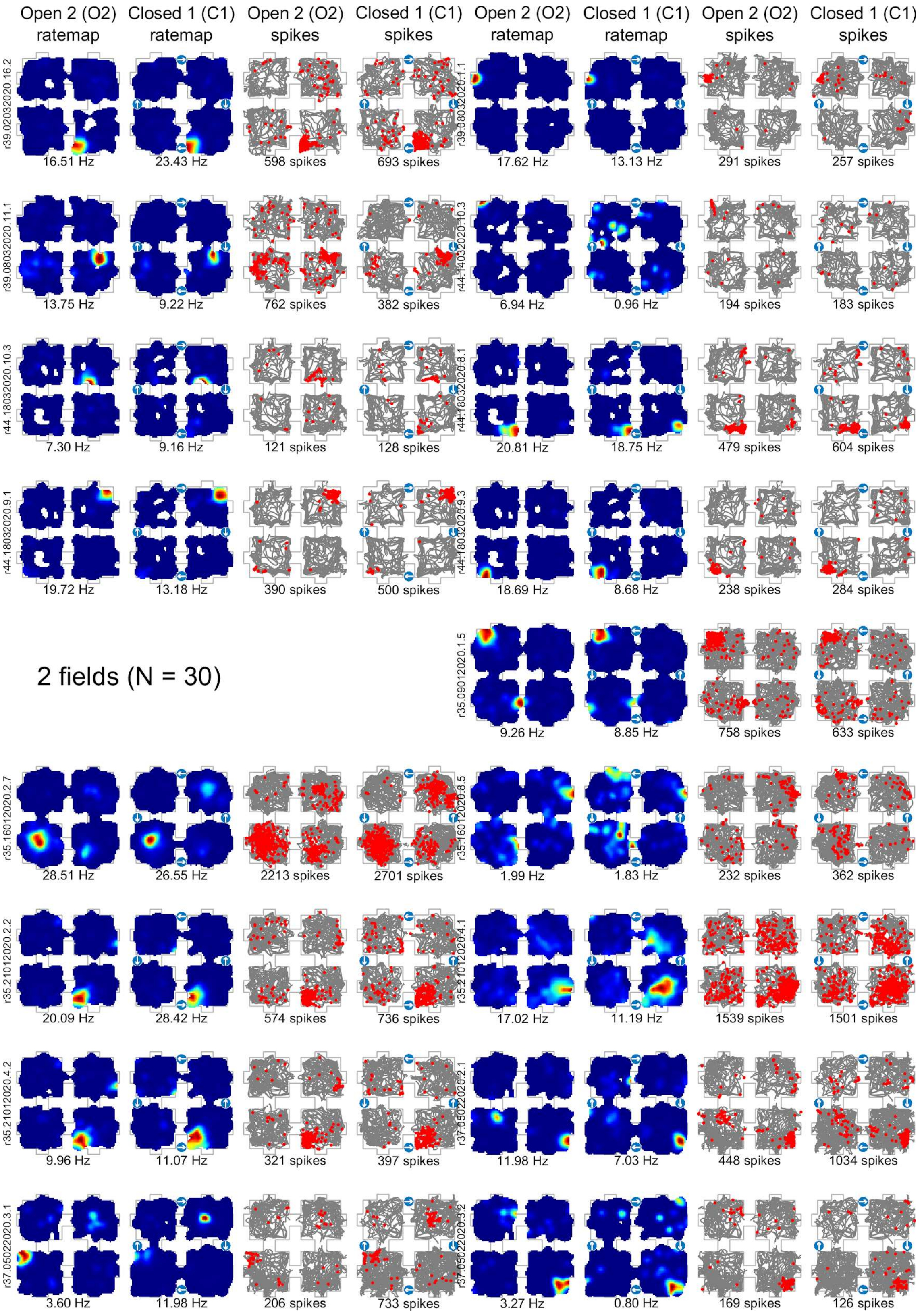
Open 2 (O2) ratemap
Closed 1 (C1) ratemap
Open 2 (O2) spikes
Closed 1 (C1) spikes
Open 2 (O2) ratemap
Closed 1 (C1) ratemap
Open 2 (O2) spikes
Closed 1 (C1) spikes
r39.02032020.16.2
16.51 Hz
23.43 Hz
598 spikes
693 spikes
r39.08032020.1.1
17.62 Hz
13.13 Hz
291 spikes
257 spikes
r39.08032020.11.1
13.75 Hz
9.22 Hz
762 spikes
382 spikes
r44.14032020.10.3
6.94 Hz
0.96 Hz
194 spikes
183 spikes
r44.18032020.10.3
7.30 Hz
9.16 Hz
121 spikes
128 spikes
r44.18032020.8.1
20.81 Hz
18.75 Hz
479 spikes
604 spikes
r44.18032020.9.1
19.72 Hz
13.18 Hz
390 spikes
500 spikes
r44.18032020.9.3
18.69 Hz
8.68 Hz
238 spikes
284 spikes
2 fields (N = 30)
r35.09012020.1.5
9.26 Hz
8.85 Hz
758 spikes
633 spikes
r35.16012020.2.7
28.51 Hz
26.55 Hz
2213 spikes
2701 spikes
r35.16012020.8.5
1.99 Hz
1.83 Hz
232 spikes
362 spikes
r35.21012020.2.2
20.09 Hz
28.42 Hz
574 spikes
736 spikes
r35.21012020.4.1
17.02 Hz
11.19 Hz
1539 spikes
1501 spikes
r35.21012020.4.2
9.96 Hz
11.07 Hz
321 spikes
397 spikes
r37.05022020.2.1
11.98 Hz
7.03 Hz
448 spikes
1034 spikes
r37.05022020.3.1
3.60 Hz
11.98 Hz
206 spikes
733 spikes
r37.05022020.3.2
3.27 Hz
0.80 Hz
169 spikes
126 spikes

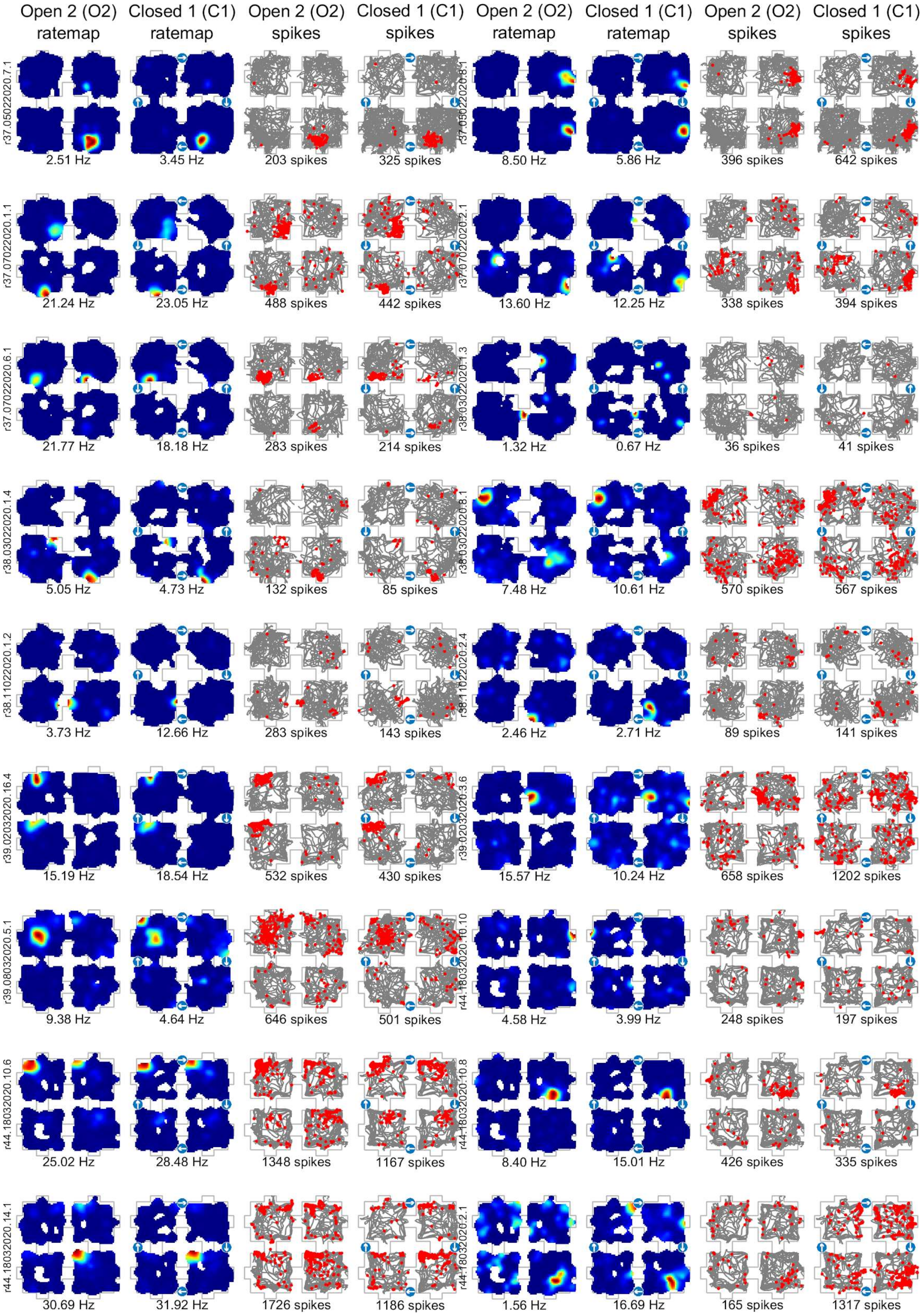
Open 2 (O2) ratemap
Closed 1 (C1) ratemap
Open 2 (O2) spikes
Closed 1 (C1) spikes
Open 2 (O2) ratemap
Closed 1 (C1) ratemap
Open 2 (O2) spikes
Closed 1 (C1) spikes
r37.05022020.7.1
2.51 Hz
3.45 Hz
203 spikes
325 spikes
r37.05022020.8.1
8.50 Hz
5.86 Hz
396 spikes
642 spikes
r37.07022020.1.1
21.24 Hz
23.05 Hz
488 spikes
442 spikes
r37.07022020.2.1
13.60 Hz
12.25 Hz
338 spikes
394 spikes
r37.07022020.6.1
21.77 Hz
18.18 Hz
283 spikes
214 spikes
r38.03022020.1.3
1.32 Hz
0.67 Hz
36 spikes
41 spikes
r38.03022020.1.4
5.05 Hz
4.73 Hz
132 spikes
85 spikes
r38.03022020.8.1
7.48 Hz
10.61 Hz
570 spikes
567 spikes
r38.11022020.1.2
3.73 Hz
12.66 Hz
283 spikes
143 spikes
r38.11022020.2.4
2.46 Hz
2.71 Hz
89 spikes
141 spikes
r39.02032020.16.4
15.19 Hz
18.54 Hz
532 spikes
430 spikes
r39.02032020.3.6
15.57 Hz
10.24 Hz
658 spikes
1202 spikes
r39.08032020.5.1
9.38 Hz
4.64 Hz
646 spikes
501 spikes
r44.18032020.10.10
4.58 Hz
3.99 Hz
248 spikes
197 spikes
r44.18032020.10.6
25.02 Hz
28.48 Hz
1348 spikes
1167 spikes
r44.18032020.10.8
8.40 Hz
15.01 Hz
426 spikes
335 spikes
r44.18032020.14.1
30.69 Hz
31.92 Hz
1726 spikes
1186 spikes
r44.18032020.2.1
1.56 Hz
16.69 Hz
165 spikes
1317 spikes

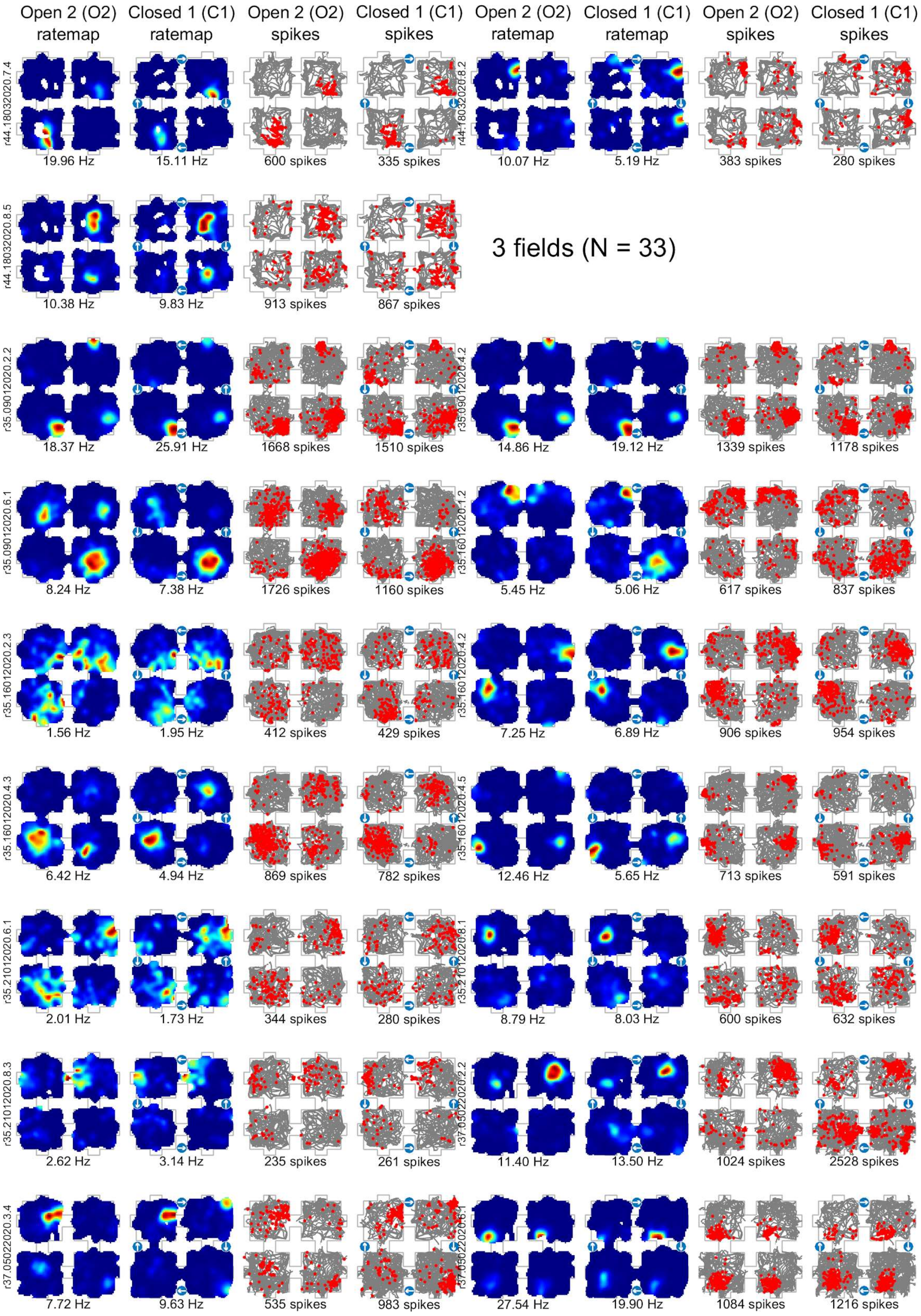
Open 2 (O2) ratemap
Closed 1 (C1) ratemap
Open 2 (O2) spikes
Closed 1 (C1) spikes
Open 2 (O2) ratemap
Closed 1 (C1) ratemap
Open 2 (O2) spikes
Closed 1 (C1) spikes
r44.18032020.7.4
19.96 Hz
15.11 Hz
600 spikes
335 spikes
r44.18032020.8.2
10.07 Hz
5.19 Hz
383 spikes
280 spikes
r44.18032020.8.5
10.38 Hz
9.83 Hz
913 spikes
867 spikes
3 fields (N = 33)
r35.09012020.2.2
18.37 Hz
25.91 Hz
1668 spikes
1510 spikes
r35.09012020.4.2
14.86 Hz
19.12 Hz
1339 spikes
1178 spikes
r35.09012020.6.1
8.24 Hz
7.38 Hz
1726 spikes
1160 spikes
r35.16012020.1.2
5.45 Hz
5.06 Hz
617 spikes
837 spikes
r35.16012020.2.3
1.56 Hz
1.95 Hz
412 spikes
429 spikes
r35.16012020.4.2
7.25 Hz
6.89 Hz
906 spikes
954 spikes
r35.16012020.4.3
6.42 Hz
4.94 Hz
869 spikes
782 spikes
r35.16012020.4.5
12.46 Hz
5.65 Hz
713 spikes
591 spikes
r35.21012020.6.1
2.01 Hz
1.73 Hz
344 spikes
280 spikes
r35.21012020.8.1
8.79 Hz
8.03 Hz
600 spikes
632 spikes
r35.21012020.8.3
2.62 Hz
3.14 Hz
235 spikes
261 spikes
r37.05022020.2.2
11.40 Hz
13.50 Hz
1024 spikes
2528 spikes
r37.05022020.3.4
7.72 Hz
9.63 Hz
535 spikes
983 spikes
r37.05022020.6.1
27.54 Hz
19.90 Hz
1084 spikes
1216 spikes

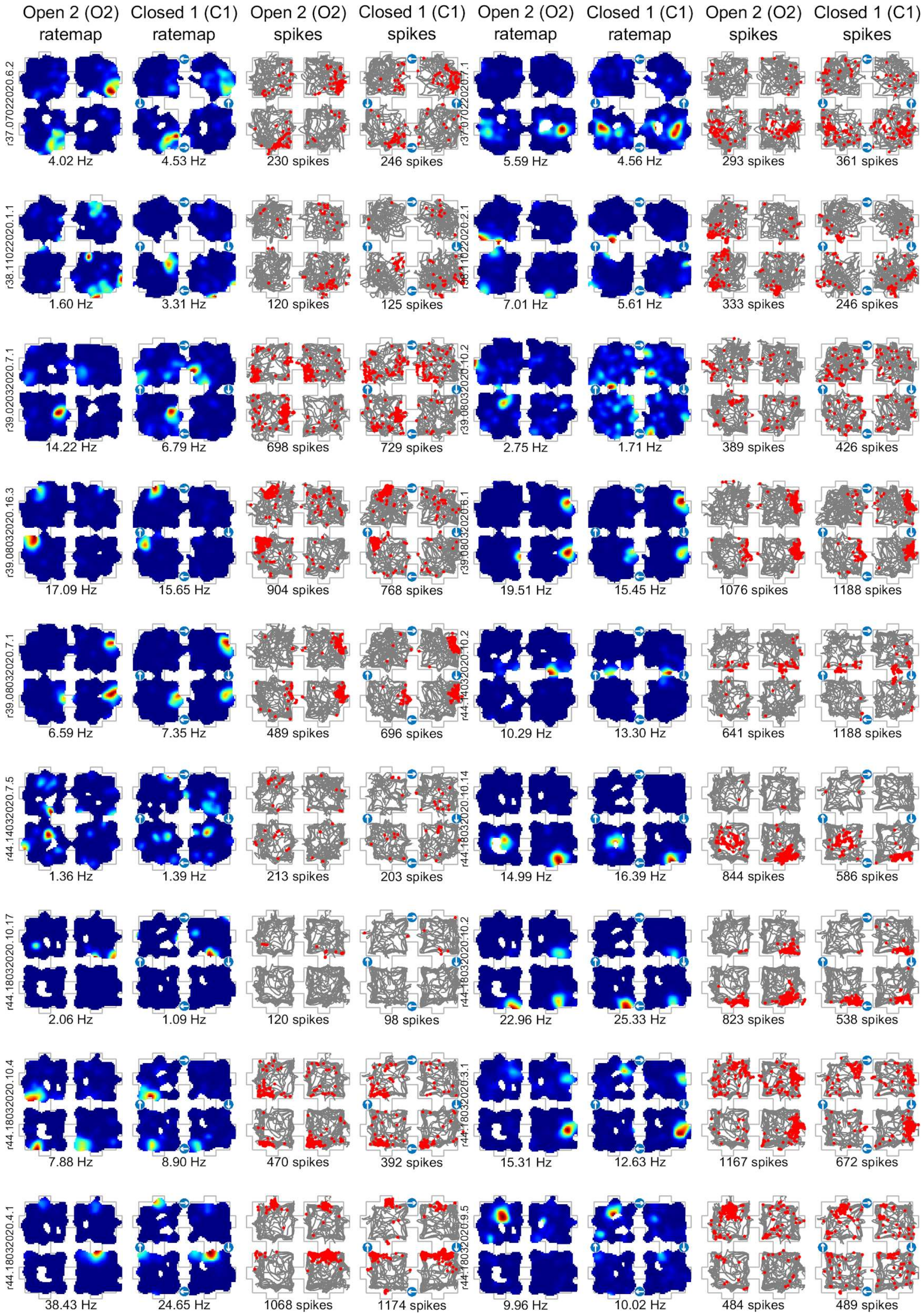
Open 2 (O2) ratemap
Closed 1 (C1) ratemap
Open 2 (O2) spikes
Closed 1 (C1) spikes
Open 2 (O2) ratemap
Closed 1 (C1) ratemap
Open 2 (O2) spikes
Closed 1 (C1) spikes
r37.07022020.6.2
4.02 Hz
4.53 Hz
230 spikes
246 spikes
r37.07022020.7.1
5.59 Hz
4.56 Hz
293 spikes
361 spikes
r38.11022020.1.1
1.60 Hz
3.31 Hz
120 spikes
125 spikes
r38.11022020.2.1
7.01 Hz
5.61 Hz
333 spikes
246 spikes
r39.02032020.7.1
14.22 Hz
6.79 Hz
698 spikes
729 spikes
r39.08032020.10.2
2.75 Hz
1.71 Hz
389 spikes
426 spikes
r39.08032020.16.3
17.09 Hz
15.65 Hz
904 spikes
768 spikes
r39.08032020.6.1
19.51 Hz
15.45 Hz
1076 spikes
1188 spikes
r39.08032020.7.1
6.59 Hz
7.35 Hz
489 spikes
696 spikes
r44.14032020.10.2
10.29 Hz
13.30 Hz
641 spikes
1188 spikes
r44.14032020.7.5
1.36 Hz
1.39 Hz
213 spikes
203 spikes
r44.18032020.10.14
14.99 Hz
16.39 Hz
844 spikes
586 spikes
r44.18032020.10.17
2.06 Hz
1.09 Hz
120 spikes
98 spikes
r44.18032020.10.2
22.96 Hz
25.33 Hz
823 spikes
538 spikes
r44.18032020.10.4
7.88 Hz
8.90 Hz
470 spikes
392 spikes
r44.18032020.3.1
15.31 Hz
12.63 Hz
1167 spikes
672 spikes
r44.18032020.4.1
38.43 Hz
24.65 Hz
1068 spikes
1174 spikes
r44.18032020.9.5
9.96 Hz
10.02 Hz
484 spikes
489 spikes

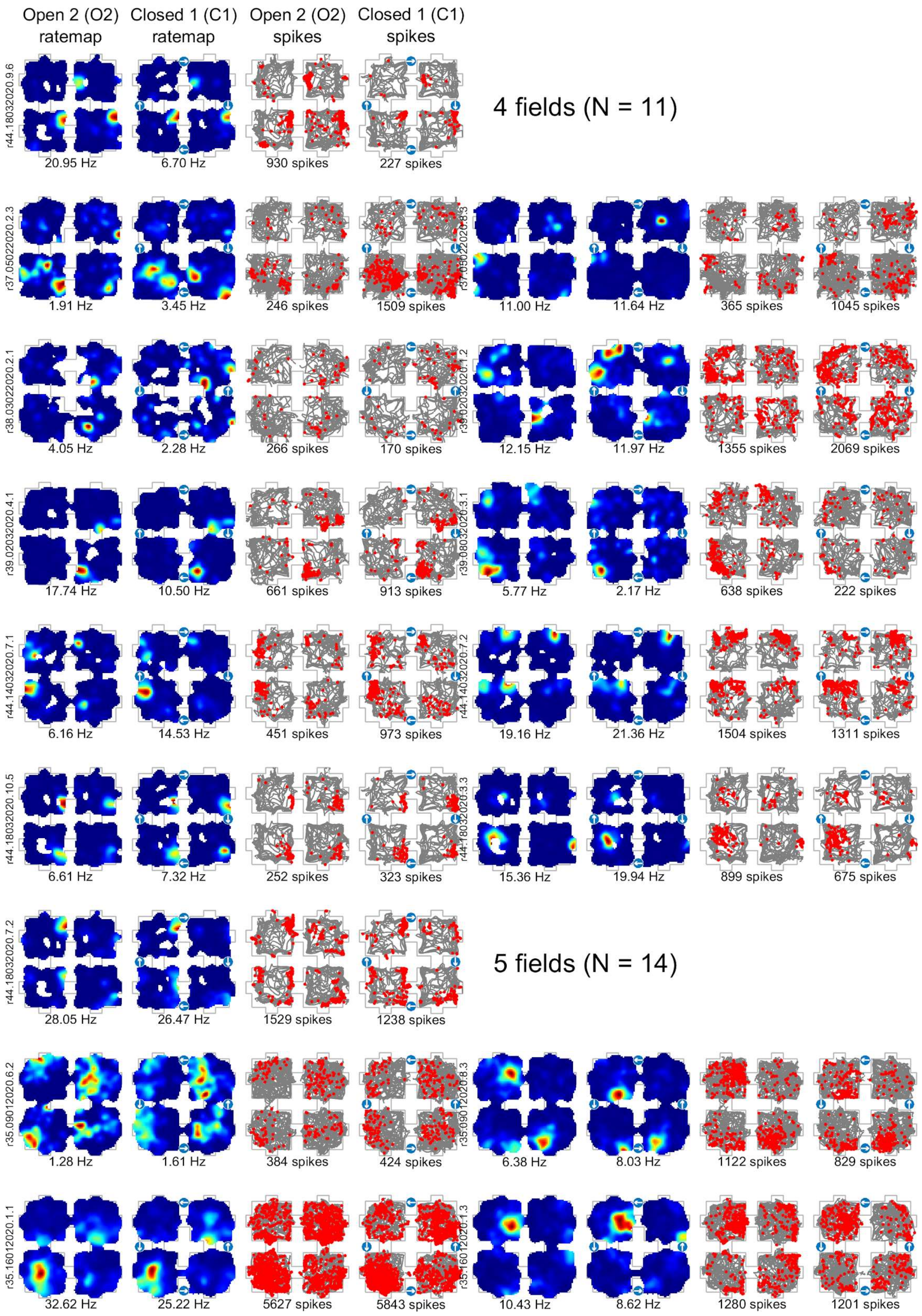
Open 2 (O2) ratemap
Closed 1 (C1) ratemap
Open 2 (O2) spikes
Closed 1 (C1) spikes
r44.18032020.9.6
20.95 Hz
6.70 Hz
930 spikes
227 spikes
4 fields (N = 11)
r37.05022020.2.3
1.91 Hz
3.45 Hz
246 spikes
1509 spikes
r37.05022020.8.3
11.00 Hz
11.64 Hz
365 spikes
1045 spikes
r38.03022020.2.1
4.05 Hz
2.28 Hz
266 spikes
170 spikes
r39.02032020.1.2
12.15 Hz
11.97 Hz
1355 spikes
2069 spikes
r39.02032020.4.1
17.74 Hz
10.50 Hz
661 spikes
913 spikes
r39.08032020.3.1
5.77 Hz
2.17 Hz
638 spikes
222 spikes
r44.14032020.7.1
6.16 Hz
14.53 Hz
451 spikes
973 spikes
r44.14032020.7.2
19.16 Hz
21.36 Hz
1504 spikes
1311 spikes
r44.18032020.10.5
6.61 Hz
7.32 Hz
252 spikes
323 spikes
r44.18032020.3.3
15.36 Hz
19.94 Hz
899 spikes
675 spikes
r44.18032020.7.2
28.05 Hz
26.47 Hz
1529 spikes
1238 spikes
5 fields (N = 14)
r35.09012020.6.2
1.28 Hz
1.61 Hz
384 spikes
424 spikes
r35.09012020.8.3
6.38 Hz
8.03 Hz
1122 spikes
829 spikes
r35.16012020.1.1
32.62 Hz
25.22 Hz
5627 spikes
5843 spikes
r35.16012020.1.3
10.43 Hz
8.62 Hz
1280 spikes
1201 spikes

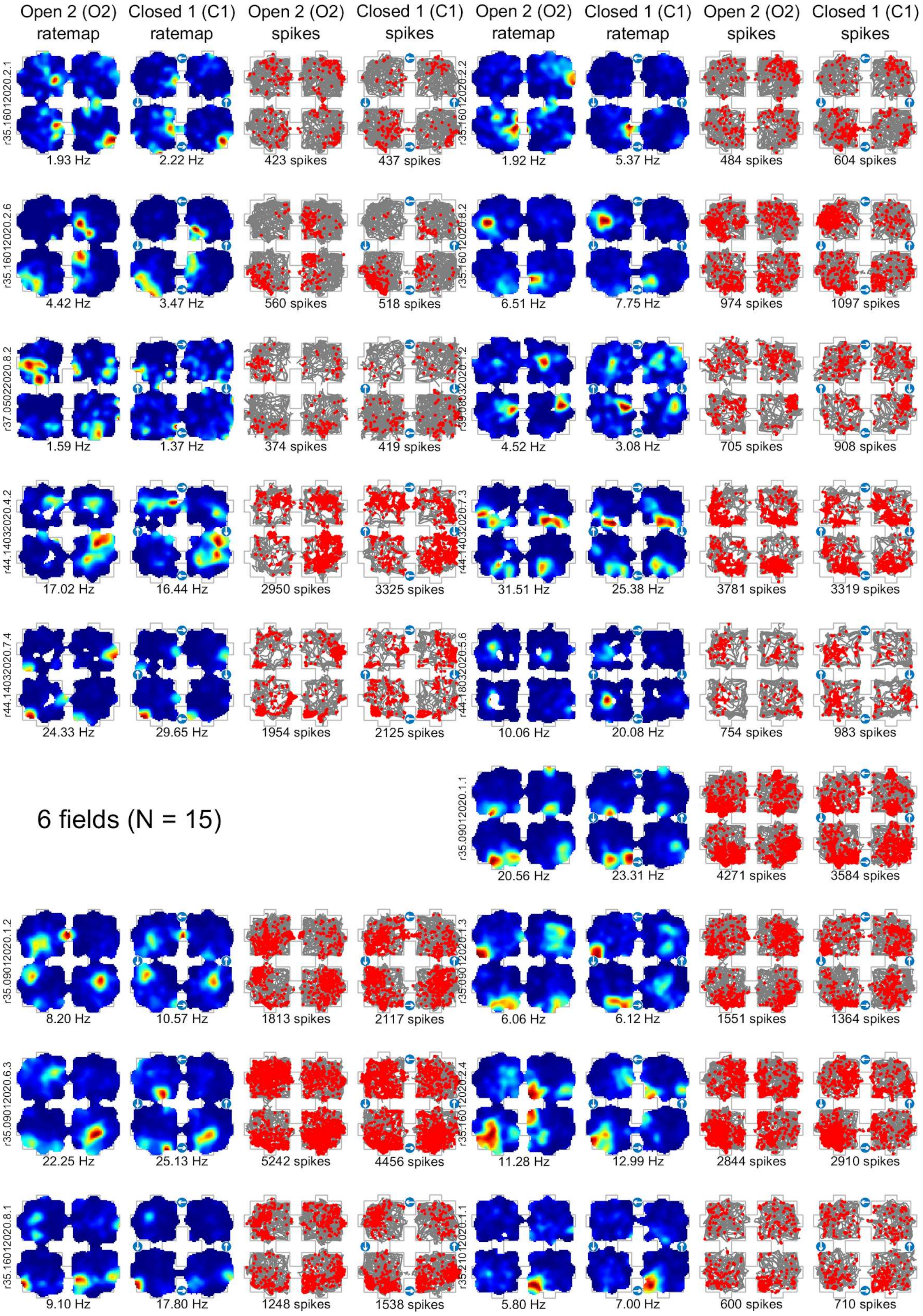
Open 2 (O2) ratemap
Closed 1 (C1) ratemap
Open 2 (O2) spikes
Closed 1 (C1) spikes
r35.16012020.2.1
1.93 Hz
2.22 Hz
423 spikes
437 spikes
r35.16012020.2.2
1.92 Hz
5.37 Hz
484 spikes
604 spikes
r35.16012020.2.6
4.42 Hz
3.47 Hz
560 spikes
518 spikes
r35.16012020.8.2
6.51 Hz
7.75 Hz
974 spikes
1097 spikes
r37.05022020.8.2
1.59 Hz
1.37 Hz
374 spikes
419 spikes
4.52 Hz
3.08 Hz
705 spikes
908 spikes
r44.14032020.4.2
17.02 Hz
16.44 Hz
2950 spikes
3325 spikes
r44.14032020.7.3
31.51 Hz
25.38 Hz
3781 spikes
3319 spikes
r44.14032020.7.4
24.33 Hz
29.65 Hz
1954 spikes
2125 spikes
r44.18032020.5.6
10.06 Hz
20.08 Hz
754 spikes
983 spikes
6 fields (N = 15)
r35.09012020.1.1
20.56 Hz
23.31 Hz
4271 spikes
3584 spikes
r35.09012020.1.2
8.20 Hz
10.57 Hz
1813 spikes
2117 spikes
r35.09012020.1.3
6.06 Hz
6.12 Hz
1551 spikes
1364 spikes
r35.09012020.6.3
22.25 Hz
25.13 Hz
5242 spikes
4456 spikes
r35.16012020.2.4
11.28 Hz
12.99 Hz
2844 spikes
2910 spikes
r35.16012020.8.1
9.10 Hz
17.80 Hz
1248 spikes
1538 spikes
r35.21012020.1.1
5.80 Hz
7.00 Hz
600 spikes
710 spikes

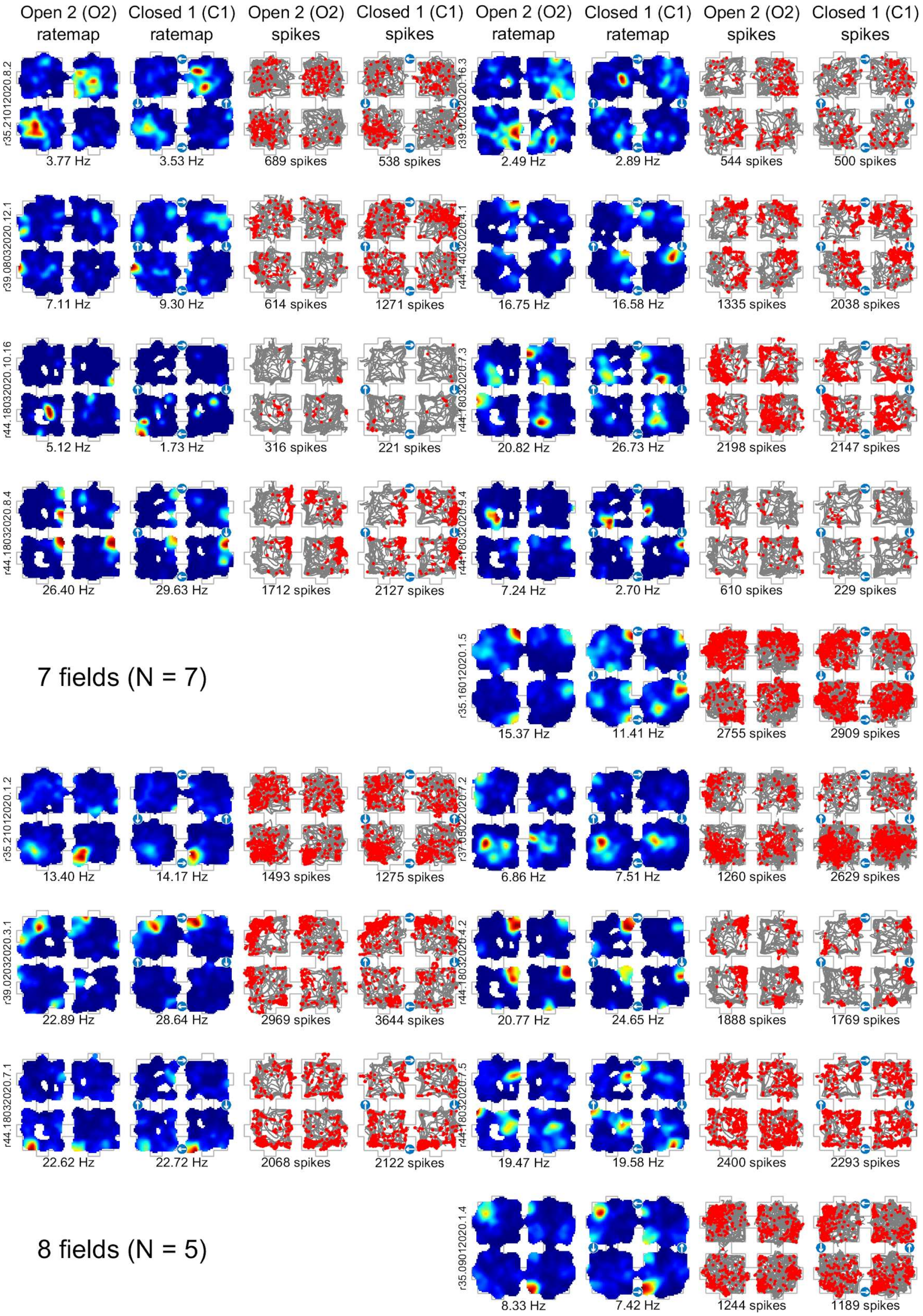

Open 2 (O2) ratemap
Closed 1 (C1) ratemap
Open 2 (O2) spikes
Closed 1 (C1) spikes
Open 2 (O2) ratemap
Closed 1 (C1) ratemap
Open 2 (O2) spikes
Closed 1 (C1) spikes
r35.21012020.8.2
3.77 Hz
3.53 Hz
689 spikes
538 spikes
r39.02032020.16.3
2.49 Hz
2.89 Hz
544 spikes
500 spikes
r39.08032020.12.1
7.11 Hz
9.30 Hz
614 spikes
1271 spikes
r44.14032020.4.1
16.75 Hz
16.58 Hz
1335 spikes
2038 spikes
r44.18032020.10.16
5.12 Hz
1.73 Hz
316 spikes
221 spikes
r44.18032020.7.3
20.82 Hz
26.73 Hz
2198 spikes
2147 spikes
r44.18032020.8.4
26.40 Hz
29.63 Hz
1712 spikes
2127 spikes
r44.18032020.9.4
7.24 Hz
2.70 Hz
610 spikes
229 spikes
7 fields (N = 7)
r35.16012020.1.5
15.37 Hz
11.41 Hz
2755 spikes
2909 spikes
r35.21012020.1.2
13.40 Hz
14.17 Hz
1493 spikes
1275 spikes
6.86 Hz
7.51 Hz
1260 spikes
2629 spikes
r39.02032020.3.1
22.89 Hz
28.64 Hz
2969 spikes
3644 spikes
r44.18032020.4.2
20.77 Hz
24.65 Hz
1888 spikes
1769 spikes
r44.18032020.7.1
22.62 Hz
22.72 Hz
2068 spikes
2122 spikes
r44.18032020.7.5
19.47 Hz
19.58 Hz
2400 spikes
2293 spikes
8 fields (N = 5)
r35.09012020.1.4
8.33 Hz
7.42 Hz
1244 spikes
1189 spikes

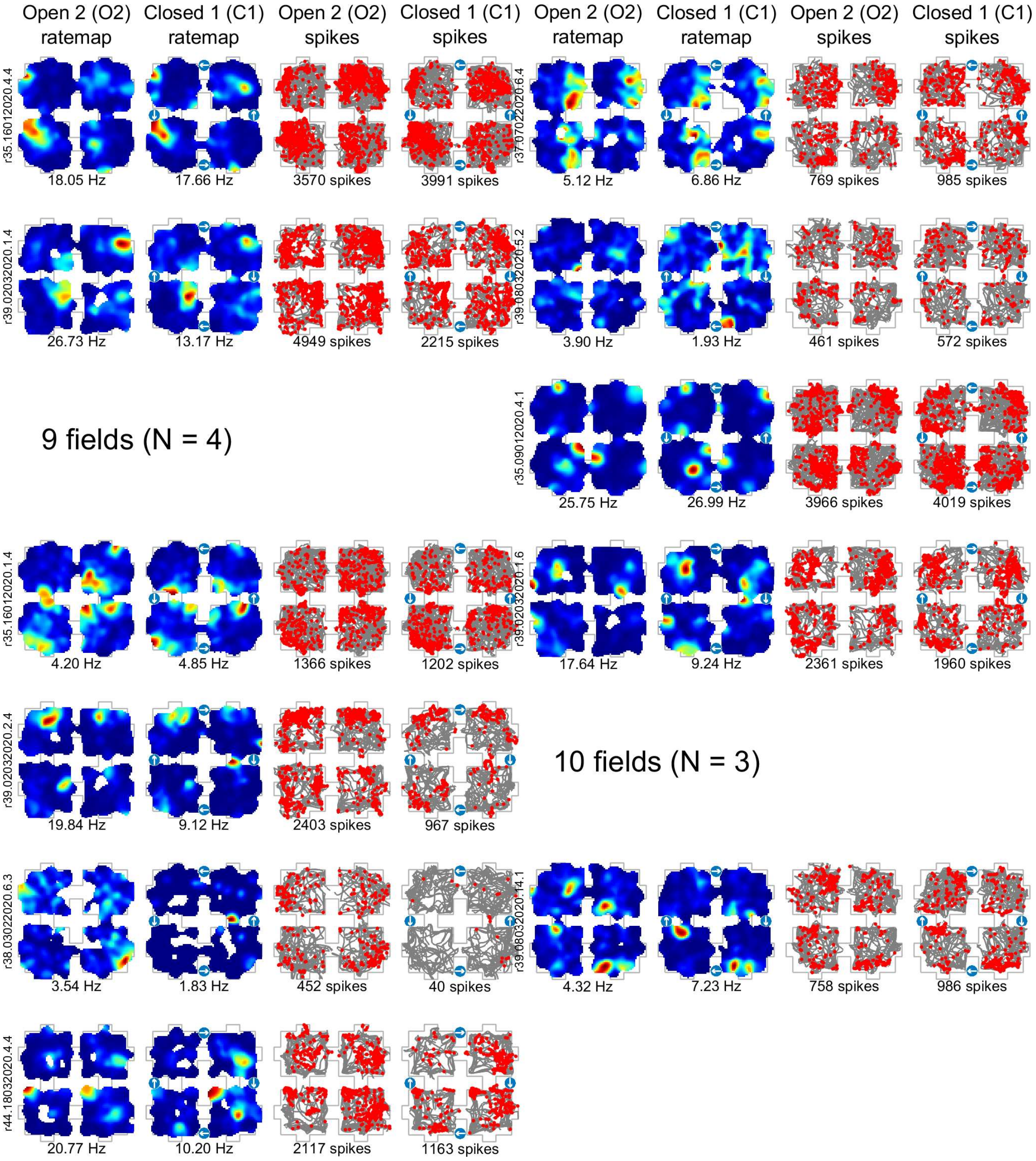

Open 2 (O2) ratemap
Closed 1 (C1) ratemap
Open 2 (O2) spikes
Closed 1 (C1) spikes
Open 2 (O2) ratemap
Closed 1 (C1) ratemap
Open 2 (O2) spikes
Closed 1 (C1) spikes
r35.16012020.4.4
18.05 Hz
17.66 Hz
3570 spikes
3991 spikes
r37.07022020.6.4
5.12 Hz
6.86 Hz
769 spikes
985 spikes
r39.02032020.1.4
26.73 Hz
13.17 Hz
4949 spikes
2215 spikes
r39.08032020.5.2
3.90 Hz
1.93 Hz
461 spikes
572 spikes
9 fields (N = 4)
r35.09012020.4.1
25.75 Hz
26.99 Hz
3966 spikes
4019 spikes
r35.16012020.1.4
4.20 Hz
4.85 Hz
1366 spikes
1202 spikes
r39.02032020.1.6
17.64 Hz
9.24 Hz
2361 spikes
1960 spikes
r39.02032020.2.4
19.84 Hz
9.12 Hz
2403 spikes
967 spikes
10 fields (N = 3)
r38.03022020.6.3
3.54 Hz
1.83 Hz
452 spikes
40 spikes
r39.08032020.14.1
4.32 Hz
7.23 Hz
758 spikes
986 spikes
r44.18032020.4.4
20.77 Hz
10.20 Hz
2117 spikes
1163 spikes
